# Localization of Mutant Huntingtin with HTT Exon1 P90 C-terminal Neoepitope Antibodies in Relation to Regional and Neuronal Vulnerability in Forebrain in Q175 Mice and Human Huntington’s Disease

**DOI:** 10.1101/2025.08.06.668984

**Authors:** Yunping Deng, Marion Joni, Hongbing Wang, Rachel Cox, Ellen Sapp, Marian DiFiglia, Anton Reiner

**Affiliations:** Department of Anatomy and Neurobiology, The University of Tennessee Health Science Center, Memphis, TN 38163; Department of Neurology, Massachusetts General Hospital, Charlestown, MA 02129, USA

**Keywords:** pathogenesis, mutant HTT1a, basal ganglia, immunohistochemistry, regional vulnerability

## Abstract

**Background:** Recent evidence suggests that accumulation of mutant exon 1 protein (HTT1a) may be critical to HD pathogenesis, but the relation of this to differential regional and cellular vulnerability in HD is unknown.

**Objective:** We assessed the contribution of the accumulation of the mutant huntingtin HTT1a to the regional and cellular variation in HD brain pathology by determining if more vulnerable regions and neuron types were relatively enriched.

**Methods:** We performed immunolabeling using the novel monoclonal antibodies 11G2 and 1B12 against the C-terminal proline 90 (P90) neoepitope of huntingtin HTT1a, which detect accumulation of monomeric, oligomeric and aggregated mutant HTT1a, on forebrain of Q175 and R6/2 mice and human HD cases.

**Results:** Diffuse nuclear and aggregate immunolabeling increased in abundance in Q175 with age, with striatal projection neurons showing immunolabeling earlier than cortical neurons, and only neuropil immunolabeling prominent in pallidal regions. Nonetheless, some regions less affected in HD, such as hippocampus, were rich in mutant HTT1a as well. In humans, striatal immunolabeling was sparser than in mouse, and mainly in the neuropil, but sparser in striatal target areas. In human HD cortex, the P90 antibodies detected predominantly neuropil aggregates, which appeared to, in part, localize to dendrites. Immunostaining in mouse and human could be blocked with HTT1a target peptide, demonstrating antibody specificity.

**Conclusions:** Our results indicate that mutant HTT1a burden appears to partly account for overall differential forebrain regional vulnerability in HD, but additional factors may contribute to vulnerability differences among forebrain regions and between specific neuron types.

**Plain Language Summary:** Huntington’s disease (HD) is caused by a mutant gene that is passed from one generation to the next. The mutant gene causes production of a mutant variant of an otherwise valuable protein called huntingtin. This mutant protein is thought to gradually damage neurons in the brain, leading to such extensive loss in a part of the brain called the basal ganglia that movement is impaired. The disability is eventually so severe, it proves fatal. It has been a mystery as to why the mutant protein might cause brain damage, but more so to some brain areas than others. One important recent clue has emerged from studies of the abnormal huntingtin protein that cells with the mutation make. Namely, rather than make a huntingtin protein of full size, as occurs normally, cells instead make only an abbreviated form of the huntingtin protein, but one that contains the abnormality. We hypothesized that if this mutant fragment is particularly toxic and the cause of the neuron damage in HD, then those brain regions that are most injured in the disease should accumulate more of this mutant fragment early in disease. We evaluated this hypothesis by examining accumulation of this mutant fragment, using a staining method selective for the fragment, in histological specimens from the brains of humans with HD and mice engineered to possess the same mutant gene as in the human disease. We found to a large extent that it is the case that those brain regions that showed the most disease-related damage, such as the basal ganglia, also accumulated the mutant huntingtin protein fragment the most. This fragment thus does appear to be toxic for neurons. Our work suggests that therapeutic efforts for HD should be directed at preventing its production or accumulation.

## Introduction

Although the greatest atrophy and neuron loss occurs within the striatal part of the basal ganglia in Huntington’s disease (HD) (Deng et al., 2004; Reiner et al., 2011), substantial atrophy and neuron loss also occur in the pallidal part of the basal ganglia, as well as in the cerebral cortex (Reiner et al., 2011). Within striatum, striatal projection neurons (SPNs) are much more vulnerable than are cholinergic interneurons (Albin et al., 1990; Reiner et al., 2011; Reiner and Deng, 2018). SPNs are, however, not all alike in their vulnerability, since enkephalinergic indirect pathway SPNs (iSPNs) projecting to globus pallidus externus (GPe) show greater loss during HD than substance P-containing direct pathway SPNs (dSPNs) projecting to globus pallidus internus (GPi) (Reiner et al., 1988; Sapp et al., 1995; Glass et al., 2000; Deng et al., 2004; Reiner and Deng, 2018). In contrast to cholinergic interneurons, GABAergic interneurons containing parvalbumin (PARV) are as vulnerable as iSPNs (Reiner et al., 2011, 2013).

The basis of the greater vulnerability of some neuron types in striatum, and the overall regional heterogeneity in HD pathology, has been uncertain, but one possibility is that it is related to heterogeneity in the production and accumulation of mutant protein (Wang et al., 2008). Surprisingly, however, prior studies have found mutant protein aggregates in only a small percent of striatal neurons in human HD patients (DiFiglia et al., 1997; Maat-Schieman et al., 1999; Kuemmerle et al., 1999; Sapp et al., 1999), despite their prominence in striatal neurons in mouse models of HD (Davies et al., 1997; Meade et al., 2002; Brooks and Dunnett, 2015).

Recent evidence has accrued that mutant huntingtin toxicity seems to occur due to production of a mutant exon 1 protein (HTT1a) stemming from a CAG-dependent splicing failure during the processing of immature mutant huntingtin transcripts, with this propensity increasing as the CAG repeat expands (Sathasivam et al., 2013; Neueder et al., 2017). This has focused attention on the accumulation of mutant HTT1a and the relationship of that accumulation to differential regional and cellular vulnerability in HD.

Our present immunolabeling study reports on the first use of recently generated antibodies directed against the C-terminal neoepitope at proline 90 (P90 based on polyQ 23 numbering) of HTT1a, namely the P90 monoclonal antibody clones 11G2 and 1B12 (Missineo et al., 2024), to characterize the forebrain distribution of mutant HTT1a in human HD brain, in the Q175 mouse (Heikkinen et al., 2012; Menalled et al., 2012; Deng et al., 2021), which like in human produces mutant HTT1a (Franich et al., 2019; Smith et al., 2023), and in R6/2 mice, which selectively produce mutant HTT1a from their transgene. We relate our findings to differential regional vulnerability in HD (Reiner et al., 2011; Waldvogel et al., 2015; Reiner and Deng, 2018), and find that HTT1a burden is consistent with some aspects of differential forebrain vulnerability in HD, but additional factors may contribute to vulnerability differences among non-basal ganglia forebrain regions and between specific neuron types in HD.

## Materials and Methods

### Anti-Huntingtin Antibodies Used

The antibodies referred to as “P90 antibodies”, specifically clones 11G2 and 1B12, are rabbit monoclonal antibodies raised against the C-terminal 8 amino acids (AEEPLHRP-OH) of human HTT1a with a free carboxylate at P90 (numbering based on 23Q) (Table 1). The specificity for the C-terminal neoepitope of HTT1a was shown by surface plasmon resonance (SPR), western blotting, ELISA, and dot blotting with counter-screening against longer and shorter HTT peptides, N-terminal HTT fragments, and full length recombinant HTT (Missineo et al., 2024). We also assessed the relationship of the mutant HTT1a detection with the P90 antibodies to detection with the PHP1 and MW8 antibodies.

**Table 1.** Antibodies used in present study. The table provides details about the antigen, catalogue number, source, antibody host and IgG type, whether or not antigen retrieval was used, and the RRID (Research Resource Identifier) number.

| Name | Antigen | Catalog # | Company | Antibody Type | Antigen Retrieval Mouse | Antigen Retrieval Human | RRID |
| --- | --- | --- | --- | --- | --- | --- | --- |
| 11G2 | KLH conjugated - C-AEEPLHRP(OH) | CHDI-90004291-4 | CHDI | rabbit monoclonal IgG | No | Yes | None |
| 1B12 | KLH conjugated - C-AEEPLHRP(OH) | CHDI-90004290-4 | CHDI | rabbit monoclonal IgG | No | Yes | None |
| PHP1 | aa 57-65 of human huntingtin | CHDI-90001515-2 | CHDI | mouse monoclonal IgG1 | No | Yes | None |
| MW8 | human exon 1 protein with 67Q | CHDI-90000942-2 | CHDI | mouse monoclonal IgG2a | No | Yes | AB_528297 |
| D7F7 | residues surrounding aa 1220 of human huntingtin | 5656 | Cell Signaling | rabbit monoclonal IgG | No | N/A | AB_10827977 |
| NeuN | aa 1-100 of synthetic human NeuN | MAB377 | Chemicon | mouse monoclonal IgG | No | N/A | AB_177621 |
| ChAT | human placental ChAT | AB 144P | Millipore | goat polyclonal IgG | No | N/A | AB_2079751 |
| PARV | carp muscle PARV | P3171 | Sigma-Aldrich | mouse monoclonal IgG | No | N/A | AB_2313693 |
| D1 | 97 amino acid C-terminal fragment of human D1 | D-187 | Sigma-Aldrich | rat monoclonal IgG | NO | N/A | AB_1840789 |
| SMI32 | nonphosphorylated 200 kD neurofilament protein | 801701 | BioLegend | mouse monoclonal IgG | NO | Yes | AB_2315330 |

PHP1 was generated against a recombinant HTT Exon1 (20Q) GST-fusion protein and detects a polyproline region of HTT1a (Table 1) (Ko et al., 2018). The MW8 antibody was generated against an aggregated form of recombinant human exon 1 (67Q) protein and was shown by peptide array epitope mapping (dot blot) to recognize the sequence AEEPLHRP (Table 1), with an early study reporting that recognition was not ablated by addition of amino acids to the C-terminus of HTT1a (Ko et al., 2001), suggesting at that time that MW8 is not selective for HTT1a, but can detect huntingtin forms elongated at the C-terminus of HTT1a. Landles et al. (2010) later confirmed that MW8 does recognize the epitope AEEPLHRP, but also showed that MW8 does not do so in the context of HTT fragments longer than HTT1a, supporting the selectivity of MW8 for pure HTT1a, like the P90 antibodies. Consistent with the interpretation that MW8 is selective for HTT1a, Smith et al. (2023) and Landles et al. (2021) reported that MW8 does not detect a signal in the brains of N171-82Q transgenic mice, which cannot produce HTT1a from their mutant HD transgene.

### Mouse Studies – Subjects

Heterozygous male Q175 HD and wildtype (WT) littermate mice were obtained from JAX at approximately 2, 6, 12, and 18 months of age (Bar Harbor, ME). The CAG repeat length was assessed by the Laragen Corporation (Culver City, CA) and found to be about 190. By convention, these HD mutant mice are nonetheless referred to as Q175 mice (Menalled et al., 2014). We also performed immunolabeling studies on tissues from three additional lines of mutant mice – R6/2 mice, N171-82Q mice, and huntingtin*-*flox mice with a tamoxifen inducible Cre recombinase (Httflox/flox;CAG-CreERT). The R6/2 mice were bred by us and their transgene had a repeat length of about 125 CAG (Reiner et al., 2012a,b; Wang et al., 2021), and they ranged in age from 9-12 weeks. N171-82Q mice were examined at 4 months of age, when they are symptomatic and feature abundant mutant protein aggregates in brain (Schilling et al., 1999). These mice were bred at Johns Hopkins University (JHU), and their brains were harvested and shipped to us by Dr. Wenzhen Duan as fixed whole brain following transcardial perfusion. The Httflox/flox;CAG-CreERT mice were 3 months old and one month post vehicle or tamoxifen treatment (Bragg et al., 2024). These mice were prepared at the University of Washington, and their brains shipped to us by Dr. Jeff Carroll as fixed whole brain following transcardial perfusion. All animal use was carried out in accordance with the National Institutes of Health Guide for Care and Use of Laboratory Animals, local animal care and use committees, and Society for Neuroscience Guidelines. The animal studies at University of Tennessee Health Science Center (UTHSC) were conducted with the approval of the UTHSC Animal Care and Use Committee (approval 21-0308) on November 8, 2021.

### Mouse Studies – Immunohistochemical Methods

Immunohistochemical studies in mice were carried out on paraformaldehyde-fixed tissue (Deng et al., 2021). The localization in Q175 mouse and R6/2 mouse brain focused on mutant huntingtin load in basal ganglia and related regions (i.e., substantia nigra and subthalamic nucleus), pallial structures of the telencephalon (which include cerebral cortex, hippocampus, and amygdala), and thalamus and hypothalamus. The studies in N171-82Q evaluated the specificity of P90 antibodies for mutant HTT1a, while the studies in Httflox/flox;CAG-CreERT mice, in which expression of WT huntingtin is greatly diminished, evaluated the nature of the P90 immunolabeling we unexpectedly saw in WT mice from both Q175 and R6/2 litters, assessing whether the labeling in WT mice might reflect production of WT HTT1a from the endogenous mouse WT huntingtin.

To assess the specificity of the P90 immunolabeling in mutant and WT mice, we also performed blocked control studies using 11G2, as described in more detail below. The species host, source, immunogen and specificity of each of the primary antibodies used to detect mutant huntingtin, and all other antibodies used as described below, are shown in Table 1. Note that specificity of all antibodies used was confirmed by the source, by us in prior studies (Deng et al., 2021; Dragatsis et al., 2018), or in the present study.

For these studies, mice were deeply anesthetized, and perfused transcardially with paraformaldehyde fixative and sectioned frozen on a sliding microtome as described previously (Deng et al., 2021). To evaluate the regional distribution of P90 labeling, the peroxidase-antiperoxidase (PAP) procedure using diaminobenzidine (DAB) as the chromogen was employed to immunolabel several sets of sections, as described previously (Meade et al., 2002; Reiner et al., 2007; Dragatsis et al., 2009; Deng et al., 2021). For blocked control studies, the 11G2 primary antibody was co-incubated overnight at 4°C with the HTT1a C-terminal octapeptide AEEPLHRP (ThermoFisher) in sodium phosphate buffer (pH 7.4) PB at 50 times more blocking peptide than antibody by weight (6 ug 11G2 to 300 ug AEEPLHRP), or without. The antibody/peptide solution was then diluted in 5% normal horse serum + 0.8% TritonX-100 + 0.01% sodium azide in PB before adding tissue sections for overnight incubation at room temperature or 3-day incubation at 4°C. As a control for the specificity of the 11G2 block with AEEPLHRP, a parallel study was performed, in which 11G2 primary antibody was co-incubated overnight at 4°C with the unrelated peptide YHRLLACLQNVHKVTTC (ThermoFisher).

Alternate sections were then incubated overnight at room temperature or for 3 days at 4°C with this antibody/control peptide solution diluted in 5% normal horse serum + 0.8% TritonX-100 + 0.01% sodium azide in PB. For the blocked control experiments, sections were incubated in horseradish peroxidase (HRP)-conjugated anti-rabbit IgG before visualization with DAB. All sections were mounted onto gelatin-coated slides, dried, cleared and coverslipped with Permount®. Mapping was conducted using high-resolution, uniformly illuminated digital images of the immunolabeled sections acquired using a digital scanner (3DHistech KFT Pannoramic 250 Flash scanner), supplemented by microscopic examination of sections. To compare the localization of the P90 HTT1a signal we saw in WT mice relative to the localization of wild-type HTT in WT mice, we used the D7F7 rabbit monoclonal antibody against the residues surrounding amino acid 1220 of human huntingtin (Cell Signalling), and PAP immunolabeling, to detect wild-type HTT in WT mice. Note, we have also shown detection of HTT1a by Western blot in mouse HD brain and its block by coincubation with C-terminal HTT1a octapeptide (Sapp et al., 2025).

To evaluate the cell-type specific localization of 11G2 signal, we used immunofluorescence co-labeling for markers of striatal and/or cortical neuron types. To identify neurons per se, we either used NeuroTrace 435/455 (ThermoFisher) as a general counterstain (1:100), or immunolabeling using a mouse monoclonal anti-NeuN (Chemicon) (1:2500).

Cholinergic striatal interneurons were visualized by immunolabeling (1:100) for choline acetyltransferase (ChAT) using a goat polyclonal antibody (Millipore), and parvalbuminergic interneurons were visualized by immunolabeling (1:5000 dilution) using a mouse monoclonal anti-PARV (Sigma-Aldrich). To assess the relative abundance of 11G2 signal in the two types of SPNs, we used a rat monoclonal antibody against the D1 dopamine receptor (Sigma-Aldrich) to distinguish dSPNs (D1-rich) from iSPNs (D1-poor). To assess the localization of mutant protein aggregates to dendrites in cerebral cortex, we combined 11G2 immunofluorescence with labeling using the SMI32 mouse monoclonal antibody (Abcam) (Thu et al., 2010; Nana et al., 2014). Secondaries conjugated to distinct Alexa fluorophores raised in compatible species (goat or donkey) and directed against appropriate species IgG were used to differentiate the various immunofluorescence markers, as in our prior studies (Meade et al., 2002; Reiner et al., 2007; Dragatsis et al., 2009; Deng et al., 2021). Finally, we used multiple immunofluorescence and confocal laser-scanning microscopy (CLSM) to assess the localization of 11G2, PHP1 and MW8 with respect to one another. Although both raised in mouse, PHP1 and MW8 could be selectively detected by different secondary antibodies because they differ in their IgG subtype (Table 1). A Zeiss 710 confocal laser-scanning microscope (Carl Zeiss AG, Oberkochen, Germany) was used to collect images of the immunofluorescence tissue. CLSM images shown here were adjusted for brightness, contrast and sharpness with Photoshop, always processing image groups correspondingly.

### Microscopic Analysis in Mouse – 11G2 Regional Scoring

We scored the regional abundance of 11G2 signal in basal ganglia and other forebrain structures at 2, 6, 12 and 18 months of age in Q175 mice on a 5-point scale, with 0 indicating no labeling and 4 indicating extremely pervasive labeling. We chose 11G2 for these studies since it yielded an identical pattern but more robust signal than 1B12. Two-three WT and Q175 mice at each age were analyzed and scored, for which the DAB-immunolabeled tissue was used. The scoring system is based on the regional density of 11G2+ immunolabeling, with a score of 1 denoting moderate diffuse perikaryal labeling, a score of 2 denoting more intense diffuse perikaryal labeling with some small 11G2+ aggregates, a score of 3 denoting diffuse perikaryal labeling and a moderate density of perikaryal and neuropil aggregates, and a score of 4 denoting a high density of perikaryal and neuropil aggregates with or without diffuse perikaryal labeling. For any given region, the labeling generally progressed from initially diffuse perikaryal only to aggregate labeling largely superseding the diffuse labeling with advancing age.

### Microscopic Analysis in Mouse – 11G2 Quantification in SPNs

Striatum in NeuroTrace-stained/NeuN-immunostained sections from mutant mice that had been immunolabeled for 11G2 in combination with D1 was analyzed for diffuse and aggregated mutant protein load at 6 and 18 months of age in Q175 mice, with two mice per age analyzed. For each mutant mouse, CLSM images of dorsolateral striatum were captured using a 40x objective, and the load per D1-rich versus D1-poor neuron types determined using NIH ImageJ. In our approach, the NeuN/NeuroTrace signal was used to create a mask of individual striatal perikarya (excluding those in the size range of cholinergic neurons), and the mask was subsequently directed to the D1 and 11G2 channels to measure the area occupied for each cell body by thresholded D1 signal and thresholded 11G2 signal, respectively. This approach for analyzing signal in multiple channels is similar to that used by us in Honig et al. (2021). We regarded the more D1-enriched half of the total striatal population analyzed as dSPNs, and the more D1-poor half of the total striatal population analyzed as iSPNs. For statistical comparison of the two SPN types, t-tests were performed, using Excel.

### Humans Studies - Subjects

Immunolabeling, imaging, and analysis were performed in human tissue as in our prior studies (Deng et al., 2004; Reiner et al., 2013; Reiner et al., 2012; Deng et al., 2021). Coronal tissue blocks containing caudate and putamen at a rostral level and/or at a level through globus pallidus (GP), blocks containing substantia nigra, blocks containing hippocampus, and blocks containing several different cortical gyri were obtained for 22 HD cases (male=11, female=11) that were verified by pathology, symptoms, family history, and/or CAG repeat (Table 2). All human studies were conducted on de-identified postmortem tissue, and thus the University of Tennessee Health Science Center Institutional Review Board waived the requirement for human subject approval (16-04430-NHSR). The cortical areas we focused on were Brodmann’s areas BA9 and BA46 (frontal cortex), area BA4 (primary motor cortex), BA6 (premotor cortex), BA13 (dysgranular insular cortex), area BA7 (posterior parietal cortex), and BA22 (superior temporal cortex). The tissues were obtained from the University of Tennessee Health Science Center (UTHSC), the University of Michigan Medical Center (Ann Arbor, MI), the National Neurological Resource Bank (NNRB, Los Angeles, CA), the Douglas Hospital Research Center (DHRC, Montreal, Canada), the University of Rochester (UR), Vanderbilt University (VU), and the Alabama Brain Collection of the University of Alabama at Birmingham (UAB). Brains had been immersion-fixed in formalin after autopsy. HD cases were staged according to Vonsattel et al. (1985). Note that because of limitations on tissue availability, not all cases were used for the full set of structures examined, as detailed in Supplementary Table 1.

**Table 2.** Human cases used in present study. Case identification (ID), source (see text for abbreviations), sex, HD grade, age at death, CAG, and CAP100, and postmortem delay (PMD) for the human cases used in present study. CAP100 is the shorthand for the CAG repeat length times age product, with constants included in the formula that yield a CAP100 score of 100 at the age of motor onset. In those cases in which we did not have the CAG repeat, we estimated it from the known relationship between CAG repeat and age at death, and these are shown in red (Keum et al., 2016; Schultz et al., 2020). To assess the relationship between the disease burden imposed by years of disease and repeat length, we calculated the CAP100 score for our HD cases (Warner et al., 2022), with those based on an estimated CAG shown in red. The red asterisks indicate this caveat for CAG and CAP100. Tissue source abbreviation are provided in the Methods section.

| Case ID | Source | Sex | HD Grade | Age at Death | CAG* | CAP100* | PMD |
| --- | --- | --- | --- | --- | --- | --- | --- |
| HD0-1 | VU | Male | pre | 28 | 47 | 73.3 | 6.0 |
| HD1-1 | UAB | Male | 1 | 49 | 43 | 98.2 | 18.0 |
| HD1-2 | UAB | Male | 1 | 55 | 44 | 118.6 | 21.0 |
| HD1-3 | UM | Male | 1 | 87 | 36 | 80.4 | 10.0 |
| HD1-4 | UCLA | Male | 1 | 57 | 43 | 114.2 | 15.0 |
| HD1-5 | DHRC | Male | 1 | 60 | 43 | 120.2 | 14.0 |
| HD2-1 | UAB | Female | 2 | 56 | 43 | 112.2 | 13.0 |
| HD2-2 | UR | Male | 2 | 74 | 38 | 91.2 | 16.0 |
| HD2-3 | UR | Female | 2 | 62 | 42 | 114.6 | NA |
| HD2-4 | UR | Male | 2 | 67 | 41 | 113.6 | 10.0 |
| HD2-5 | UR | Female | 2 | 70 | 40 | 107.9 | 6.0 |
| HD2-6 | UR | Female | 2 | 62 | 42 | 114.6 | 16.0 |
| HD3-1 | UCLA | Male | 3 | 72 | 40 | 110.9 | 15.0 |
| GD3-2 | UCLA | Male | 3 | 54 | 44 | 116.5 | 4.0 |
| HD3-3 | UR | Male | 3 | 76 | 40 | 117.1 | 18.0 |
| HD3-4 | DHRC | Female | 3 | 73 | 40 | 112.5 | 18.0 |
| HD3-5 | UM | Female | 3 | 60 | 43 | 120.2 | 12.0 |
| HD3-6 | UR | Female | 3 | 84 | 36 | 77.7 | 14.0 |
| HD4-1 | UR | Female | 4 | 52 | 46 | 128.2 | 14.0 |
| HD4-2 | UAB | Female | 4 | 70 | 38 | 86.3 | 17.0 |
| HD4-3 | UAB | Female | 4 | 58 | 48 | 160.9 | 43.0 |
| HD4-4 | UAB | Female | 4 | 78 | 41 | 132.2 | 24.0 |
| **Control** |  | **4F, 3M** | **Control** | **63.6** | **N/A** | **N/A** | **25.4** |
| **HD** |  | **11F, 11M** | **HD** | **63.8** | **41.7** | **110.1** | **15.4** |

### Immunohistochemical Methods in Human

Tissue blocks for HD and/or control cases were transferred to 20% sucrose/10% glycerol/0.1M sodium phosphate buffer (PB) (pH 7.4). Frozen sections were cut at 80 µm with a sliding microtome, and then processed by immunohistochemistry. To enhance immunolabeling, sections were immersed for 30 min in 1% sodium borohydride in PB, 0.1 M DL-lysine in PB, followed by 0.2% nonfat dry milk in PB. We additionally used heat-based antigen retrieval as described in our prior studies (Jiao et al., 1999; Richfield et al., 2002; Deng et al., 2004). The primary antibodies used in human were ones also used in mouse (Table 1) and immunolabeling methods were as for mouse, with 1% bovine serum albumin added (Deng et al., 2004; Reiner et al., 2013). The free-floating sliding microtome sections were incubated in microcentrifuge vials for 48–72 h at 4°C. After incubation in primary antibody, a set of sections from each case was processed by the PAP method or by using anti-rabbit IgG conjugated to HRP as secondary, followed by visualization with DAB, for mapping purposes. The immunolabeled sections were mounted onto gelatin-coated slides, dried, cleared and coverslipped with Permount®. We also used multiple immunofluorescence and CLSM to assess the colocalization of 11G2, PHP1 and MW8, using the same immunolabeling methods as in mouse, as well as to examine the colocalization of 11G2 with SMI32. Sudan Black B treatment was used to suppress autofluorescence in human brains by incubation of slide-mounted sections in 0.3% Sudan Black B for 10 minutes, followed by rinsing in TBS prior to coverslipping. For all human immunofluorescence, the sections were counterstained with NeuroTrace (1:100) for cytoarchitectonic detail. As in mouse, we scored the regional abundance of 11G2 in basal ganglia structures (somatic and limbic both) and other forebrain structures on a 5-point scale, with 0 indicating no labeling and 4 indicating extremely pervasive labeling, using the IgG-HRP/DAB-immunolabeled tissue. To assess the specificity of the P90 immunolabeling in HD brain, we performed blocked control studies using 11G2, co-incubating the 11G2 antibody with the HTT1a C-terminal octapeptide or unrelated peptide, as described above for mouse.

## Results

### P90 Labeling in Q175 and WT Mouse Basal Ganglia, Substantia Nigra and Subthalamic Nucleus

In our initial screening, we found that both the 11G2 and 1B12 antibodies detected mutant HTT1a accumulation in basal ganglia of 2-, 6-, 10-, 12-, and 18-month old Q175 mice. The regional patterns appeared identical, but the labeling was slightly more intense for 11G2. Accordingly, the 11G2 antibody was subsequently used to prepare 2-3 complete series of sections through forebrain and midbrain for mapping purposes, which are also largely used for the following description and exclusively for the illustrations. At 2 months of age, we observed diffuse nuclear immunolabeling with both 11G2 and 1B12 in the vast majority of neurons in both somatic striatum (caudate-putamen) and limbic striatum (nucleus accumbens and olfactory tubercle), and thus the vast majority of SPNs (Fig. 1F-H), as also reported by Smith et al. (2023) using the S830 antibody. In the older Q175 mice, we additionally detected neuropil and nuclear aggregates, and gradually lessened diffuse nuclear labeling in striatum. The nuclear and neuropil aggregates increased in size and abundance with age. The diffuse nuclear labeling appeared granular at higher magnification viewing, and the lessened diffuse labeling likely reflects the coalescence of the diffusely localized antigen into aggregates. Neurons in GPe, GPi, SNr and ventral pallidum were rarely labeled by the P90 antibodies. Rather, neuropil aggregates gradually increased in frequency in these SPN target areas as the mutant mice aged. Nuclear immunolabeling was seen in only a minority of subthalamic nucleus (STN) cell bodies in older Q175 mice, but fine neuropil aggregates were more numerous.

**Figure 1.**
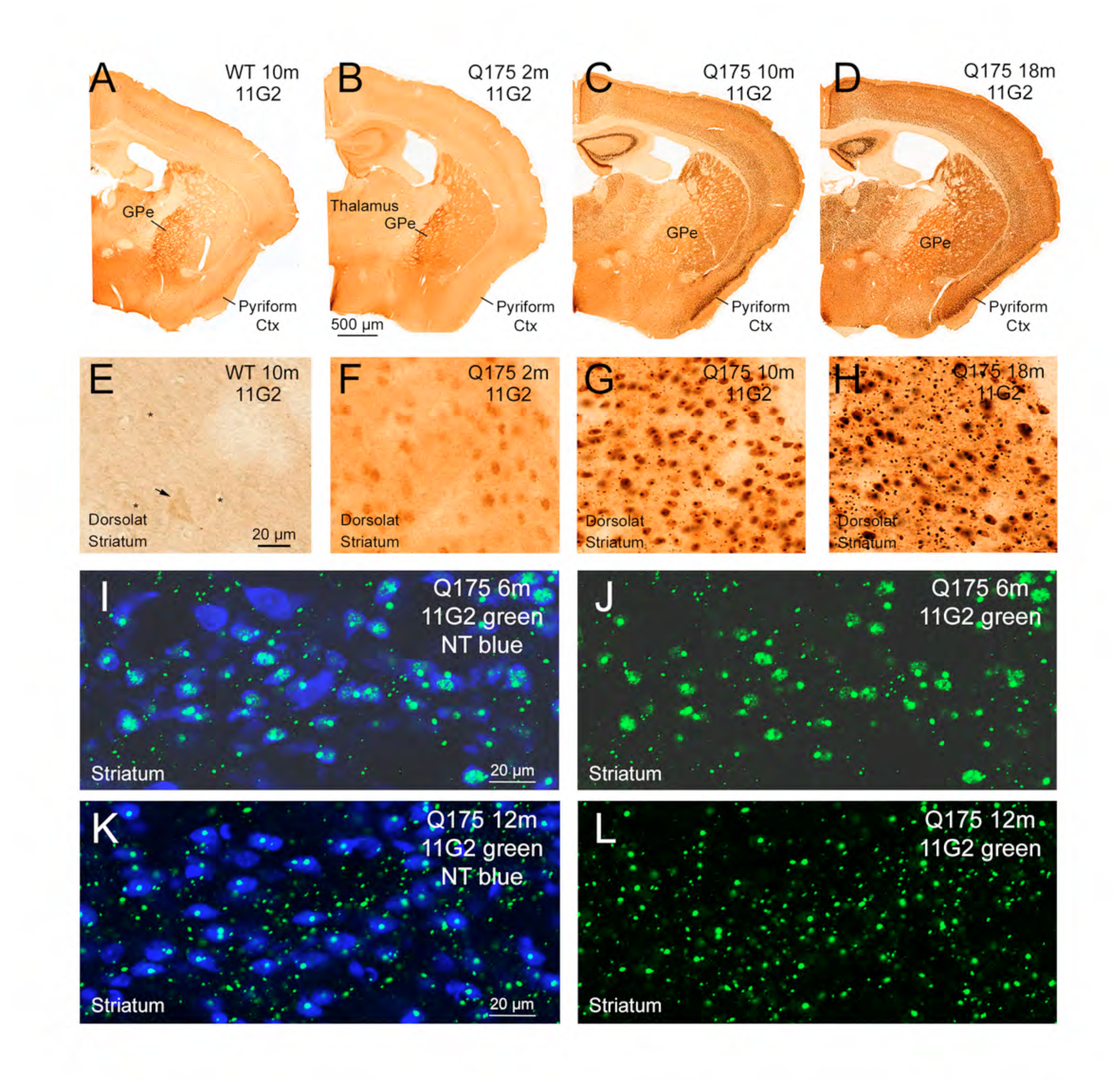
The first two rows of images show P90 immunolabeling with the 11G2 antibody in 2-month old (B, F), 10-month old (C, G), and 18-month old Q175 (D, H) mice compared to a 10-month old WT littermate control (A, E), with the first row showing an entire section through the right side of the forebrain at the level of GPe, and the second row showing a view of right dorsolateral striatum. 11G2 immunolabeling of perikarya in the Q175 mice at 2 months is diffuse and appears to be nuclear, with nuclear and neuropil aggregates forming with age progression, and lessened diffuse nuclear labeling. Images I and K show a merged CLSM image of striatum from a 6-month old Q175 mouse and a 12-month old Q175 mouse, respectively, with striatal neurons labeled blue with NeuroTrace (NT) and 11G2 immunolabeling in green. Images J and L show the 11G2 immunolabeling alone for these cases. The CLSM images show diffuse and aggregated 11G2 signal in striatal neurons at 6 months, with nuclear and neuropil aggregates more prominent at 12 months. Note, however, that image A shows that 11G2 immunolabeling is present in terminals in GPe in WT mice. The higher power view in image E of WT striatum shows that light cytoplasmic 11G2 immunolabeling is present in numerous presumptive striatal projections neurons in WT mouse (some indicated by small asterisks), and large presumptive cholinergic interneurons (arrow). Scale bar in B applies to all images in first row, scale bar in E applies to all images in second row, scale bar in I applies to both images in third row and scale bar in K applies to both images in last row.

We also saw distinct 11G2 and 1B12 immunolabeling in basal ganglia of WT littermates of Q175 mice at all ages, immunolabeling that, however, was very different than the nuclear and aggregate signal seen in the Q175 mice. In particular, focusing only on basal ganglia structures and mainly our results with the 11G2 antibody, we saw light but distinct cytoplasmic immunolabeling of the perikarya of large striatal neurons (likely cholinergic interneurons), and of presumptive SPN terminals in GPe, GPi, SNr and ventral pallidum in the WT mice (Fig. 1A, E, 2A). At higher magnification, the 11G2 signal in WT GPe could be seen to be in axons and puncta (Fig. 2C) that closely resembled the SPN axons and terminals in GPe detectible with ENK immunolabeling, and the 11G2 signal in WT GPi and SN could be seen to be in axons and puncta that closely resembled the SPN axons and terminals in GPi and SN detectible with SP immunolabeling (Deng et al., 2021). Notably, the immunolabeling in basal ganglia of WT mice resembled what we have seen with antibodies against WT huntingtin (Fusco et al., 1999; Dragatsis et al., 2018). This P90 immunolabeling in WT mice was unexpected because of the evidence that the P90 antibodies are specific for HTT1a and because prior studies have largely emphasized the production of HTT1a as an event that is associated with a CAG expansion in the gene sequence coding for HTT1a. Additionally, we saw 11G2 (Figs. 1B, 2B, 2D) and 1B12 immunolabeling of presumptive SPN terminals in GPe, GPi, SNr and ventral pallidum in 2-month old Q175 mice, but not in older mutants. P90 immunolabeling of the cytoplasm of striatal neurons was not evident in 2-month or older Q175 mice, with the diffuse nuclear labeling instead prevailing at early ages, and aggregates as well at later ages.

**Figure 2.**
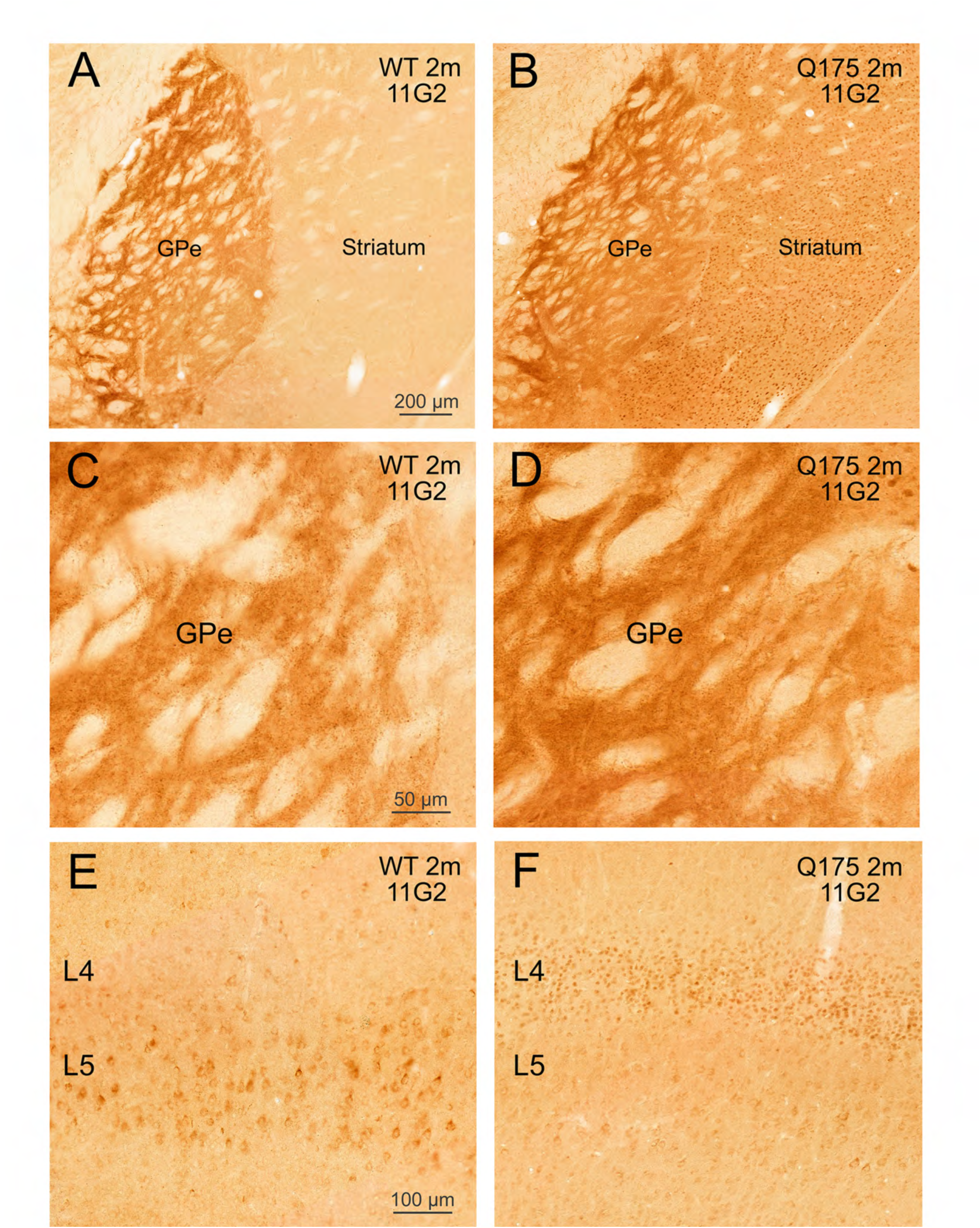
The first two rows of images show 11G2 immunolabeling in a 2-month old Q175 (B, D) mouse compared to a 2-month old WT littermate control (A, C), with higher magnification views of GPe than in Figure 1A, B. Note that in both the WT and Q175 mouse, GPe is identifiable by a dense 11G2 neuropil immunolabeling in panels A and B, with the higher magnification view in panels C and D showing that the immunolabeling represents a dense mesh of 11G2+ fibers and terminals. Images E and F show 11G2 immunolabeling of somatosensory cortex in a 2-month old Q175 mouse (F) compared to a 2-month old WT littermate control (E), focusing on layers 4 and 5, providing a higher magnification view than in Figure 1A, B. Note that WT layer 5 contains numerous perikarya with distinct 11G2+ cytoplasmic labeling, while Q175 cortex has distinct 11G2+ diffuse nuclear immunolabeling of layer 4 neurons, and some very light cytoplasmic labeling of layer 5 neurons. Scale bar in A applies to B as well, scale bar in C applies to D as well, and scale bar in E applies to F as well.

We performed multiple immunofluorescence with NeuroTrace 435/455 as a counterstain on Q175 striatum to determine the neuron-type specific localization of 11G2+ immunolabeling, as imaged using CLSM. These studies showed that 11G2+ signal at 6 months of age in Q175 mice accumulates diffusely and as aggregates in the nuclei of the vast majority of striatal neurons, which by their abundance and size are likely to be SPNs (Fig. 1I-L). By this age, nuclear aggregates are, however, far more common and evident than the diffuse nuclear 11G2+ immunolabeling, which we saw predominated at 2 months of age. The CLSM imaging also showed that a mix of small and larger 11G2+ aggregates was widespread in the neuropil of striatum at 12 months. Nuclear aggregates were typically larger than neuropil aggregates. Using interneuron markers, we observed in 10-month old Q175 mice that no cholinergic (ChAT+) interneurons possessed diffuse nuclear 11G2, 13.6% possessed cytoplasmic aggregates, and 2.3% possessed nuclear aggregates (Fig. 3A-C). By contrast, 11G2 signal was much more common in PARV+ striatal interneurons, with 48.1% possessing diffuse nuclear labeling, 36.5% possessing nuclear aggregates, and 9.6% possessing cytoplasmic aggregates (Fig. 3D-F). The nuclear aggregates were often present in PARV+ interneurons also possessing diffuse nuclear P90, and nuclear aggregate size was less than in SPNs. In some cases, multiple tiny nuclear aggregates were present in PARV+ interneurons, seemingly a precursor to their coalescence into a larger aggregate. We additionally examined GPe using multiple immunofluorescence, and found that its PARV+ orthotypic neurons were devoid of diffuse nuclear 11G2+ immunolabeling, in contrast to PARV+ striatal interneurons, although about 5% of PARV+ GPe neurons possessed either nuclear or cytoplasmic aggregates (Fig. 3G-I). PARV-negative, presumptive arkypallidal, GPe neurons were also poor in 11G2+ signal.

**Figure 3.**
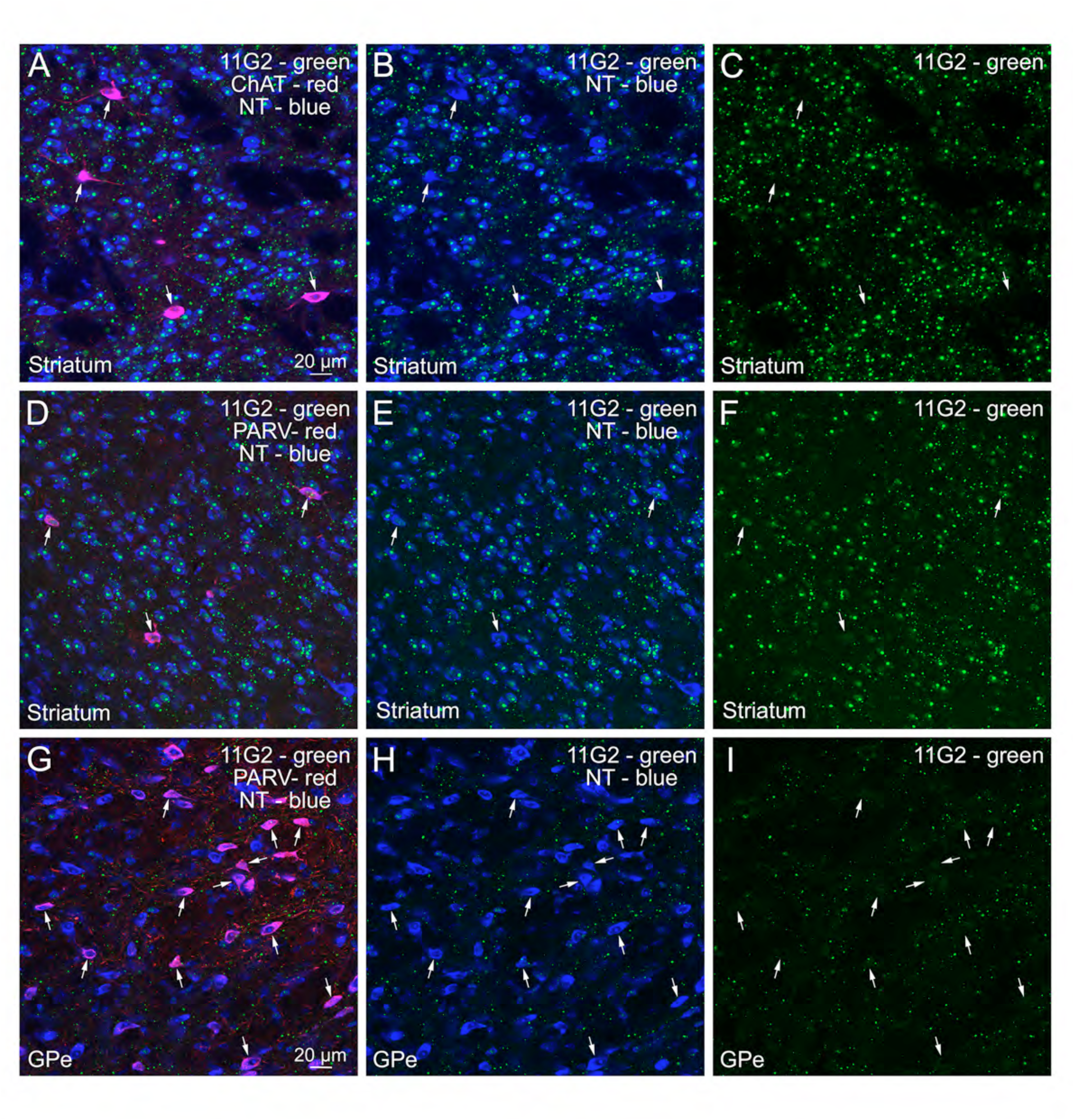
The first image in each of the first two rows shows a merged CLSM image of striatum from a 10-month old Q175 mouse with striatal neurons labeled blue with NeuroTrace, 11G2+ aggregates shown in green, and an interneuron marker (ChAT in the upper row, PARV in the second row) shown in red. The second image in each row shows the green channel and blue channel only from the merged images, and the last image in each row shows the 11G2 green channel from the merged image. The arrows in each row indicate the interneurons. Note the ChAT interneurons are largely devoid of 11G2 but the PARV interneurons commonly show diffuse and aggregated 11G2 signal in their nuclei. The last row shows a merged CLSM image of GPe from a 10-month old Q175 mouse with GPe neurons labeled blue with NeuroTrace, 11G2+ aggregates shown in green, and PARV+ orthotypic GPe neurons shown in red (some indicated by arrows). The second image in this row shows the green channel and blue channel only from the merged image, and the last image shows the 11G2 green channel from the merged image. While small 11G2+ aggregates are common in the GPe neuropil, 11G2 signal is sparse in both PARV+ orthotypic GPe neurons and PARV-negative presumptive arkypallidal GPe neurons. Scale bar in A applies to all images in first two rows, and scale bar in G applies to all images in last row.

We scored the basal ganglia 11G2 immunolabeling on a 0-4 scale at 2-, 6-, 12- and 18-months of age, with 0 indicating no labeling and 4 indicating extremely pervasive labeling, as described in the Methods section. For any given region with perikaryal signal, the labeling progressed from initially diffuse-only nuclear perikaryal localization to aggregate labeling in nuclei becoming common but not entirely superseding the diffuse nuclear labeling. Table 3 shows higher 11G2+ load for striatal structures (somatic striatum, accumbens, olfactory tubercle) than for pallidal (GPe, GPi, SNr, and ventral pallidum), which is consistent with the much higher vulnerability of striatum than pallidum in HD (Vonsattel, 2008; Reiner et al., 2011).

**Table 3.** Age-wise regional 11G2 signal in Q175 mice. Age-wise 11G2 scores for Q175 mouse for forebrain structures, scored on a scale of 0 (none) to 4 (extremely rich).

|  |  | 11G2 | 11G2 | 11G2 | 11G2 |
| --- | --- | --- | --- | --- | --- |
|  |  | Load | Load | Load | Load |
| Mouse Region | Human Equivalent | 2 mo Q175 | 6 mo Q175 | 12 mo Q175 | 18 mo Q175 |
| Rostral Medial Striatum | Rostral Caudate | 2.05 ± 0.05 (2) | 2.63 ± 0.13 (2) | 3.60 ± 0.00 (2) | 3.93 ± 0.07 (3) |
| Rostral Lateral Striatum | Rostral Putamen | 1.95 ± 0.05 (2) | 2.40 ± 0.15 (2) | 3.55 ± 0.00 (2) | 3.85 ± 0.08 (3) |
| Mid Medial Striatum | Mid Caudate | 1.95 ± 0.05 (2) | 2.55 ± 0.20 (2) | 3.50 ± 0.10 (2) | 3.93 ± 0.07 (3) |
| Mid Lateral Striatum | Mid Putamen | 1.80 ± 0.10 (2) | 2.40 ± 0.20 (2) | 3.45 ± 0.10 (2) | 3.90 ± 0.03 (3) |
| Caudal Medial Striatum | Caudal Caudate | 2.08 ± 0.03 (2) | 2.45 ± 0.20 (2) | 3.55 ± 0.05 (2) | 3.87 ± 0.09 (3) |
| Caudal Lateral Striatum | Caudal Putamen | 1.98 ± 0.08 (2) | 2.38 ± 0.28 (2) | 3.50 ± 0.05 (2) | 3.85 ± 0.05 (3) |
| All Striatum | Caudo-putamen | 1.97 ± 0.01 (2) | 2.47 ± 0.19 (2) | 3.53 ± 0.03 (2) | 3.89 ± 0.06 (3) |
| All Caudate | Caudate | 2.03 ± 0.02 (2) | 2.54 ± 0.18 (2) | 3.55 ± 0.02 (2) | 3.91 ± 0.07 (3) |
| All Putamen | Putamen | 1.91 ± 0.01 (2) | 2.39 ± 0.21 (2) | 3.50 ± 0.05 (2) | 3.87 ± 0.04 (3) |
| Globus Pallidus Externus | Globus Pallidus Externus | 0.00 ± 0.00 (2) | 0.18 ± 0.03 (2) | 0.38 ± 0.13 (2) | 1.13 ± 0.13 (2) |
| Globus Pallidus Internus | Globus Pallidus Internus | 0.00 ± 0.00 (2) | 0.25 ± 0.00 (2) | 0.45 ± 0.05 (2) | 0.88 ± 0.13 (2) |
| SN pars Reticulata | SN pars Reticulata | 0.00 ± 0.00 (2) | 0.43 ± 0.08 (2) | 0.88 ± 0.13 (2) | 1.43 ± 0.08 (2) |
| Nucleus Accumbens | Nucleus Accumbens | 1.83 ± 0.33 (2) | 2.50 ± 0.00 (2) | 3.40 ± 0.00 (2) | 3.72 ± 0.12 (2) |
| Olfactory Tubercle | Olfactory Tubercle | 1.80 ± 0.30 (2) | 2.38 ± 0.13 (2) | 3.50 ± 0.00 (2) | 3.85 ± 0.10 (3) |
| Ventral Pallidum | Ventral Pallidum | 0.00 ± 0.00 (2) | 0.33 ± 0.08 (2) | 0.75 ± 0.00 (2) | 0.83 ± 0.08 (3) |
| CA1 | CA1 | 2.13 ± 0.13 (2) | 2.70 ± 0.05 (2) | 4.00 ± 0.00 (2) | 4.00 ± 0.00 (2) |
| CA2 | CA2 | 0.04 ± 0.01 (2) | 0.45 ± 0.05 (2) | 3.00 ± 0.00 (2) | 3.50 ± 0.50 (2) |
| CA3 | CA3 | 1.00 ± 0.00 (2) | 2.33 ± 0.07 (2) | 3.90 ± 0.10 (2) | 3.95 ± 0.05 (2) |
| CA4 | CA4 | 1.00 ± 0.00 (2) | 2.20 ± 0.20 (2) | 3.40 ± 0.10 (2) | 3.83 ± 0.07 (2) |
| Dentate Granule | Dentate Granule | 0.15 ± 0.05 (2) | 2.18 ± 0.07 (2) | 4.00 ± 0.00 (2) | 4.00 ± 0.00 (2) |
| Dentate Hilus | Dentate Hilus | 0.00 ± 0.00 (2) | 0.88 ± 0.13 (2) | 3.55 ± 0.05 (2) | 3.93 ± 0.07 (2) |
| Subiculum | Subiculum | 0.00 ± 0.00 (2) | 0.88 ± 0.13 (2) | 3.13 ± 0.63 (2) | 3.28 ± 0.53 (2) |
| Hippocampal Complex | Hippocampal Complex | 0.69 ± 0.01 (2) | 1.66 ± 0.02 (2) | 3.57 ± 0.31 (2) | 3.78 ± 0.13 (2) |
| Entorhinal Ctx | Entorhinal Ctx | 0.00 ± 0.00 (2) | 1.70 ± 0.05 (2) | 3.00 ± 0.25 (2) | 3.40 ± 0.00 (2) |
| Pyriform Ctx | Pyriform Ctx | 0.00 ± 0.00 (2) | 2.63 ± 0.13 (2) | 3.50 ± 0.00 (2) | 4.00 ± 0.00 (3) |
| Basolateral Amygdala | Basolateral Amygdala | 0.00 ± 0.00 (2) | 1.38 ± 0.38 (2) | 1.83 ± 0.08 (2) | 2.50 ± 0.00 (2) |
| Basomedial Amygdala | Basomedial Amygdala | 0.00 ± 0.00 (2) | 0.55 ± 0.20 (2) | 1.78 ± 0.03 (2) | 1.50 ± 0.50 (2) |
| Cortical Amygdala | Cortical Amygdala | 0.00 ± 0.05 (2) | 0.55 ± 0.20 (2) | 1.88 ± 0.13 (2) | 1.50 ± 0.50 (2) |
| Central Amygdala | Central Amygdala | 0.00 ± 0.00 (2) | 1.25 ± 1.0 (2) | 2.75 ± 0.25 (2) | 2.75 ± 0.25 (2) |
| Medial Amygdala | Medial Amygdala | 0.00 ± 0.00 (2) | 0.30 ± 0.05 (2) | 1.13 ± 0.13 (2) | 1.50 ± 0.25 (2) |
| Amygdalar Complex | Amygdalar Complex | 0.00 ± 0.00 (2) | 0.81 ± 0.21 (2) | 1.87 ± 0.07 (2) | 1.95 ± 0.20 (2) |
| All Frontal Cortex | All Frontal Cortex | 0.00 ± 0.00 (2) | 1.76 ± 0.01 (2) | 3.19 ± 0.06 (2) | 3.93 ± 0.07 (2) |
| PRL Frontal Ctx | DL Med Frontal Ctx (A9) | 0.00 ± 0.00 (2) | 1.88 ± 0.07 (2) | 3.25 ± 0.00 (2) | 3.93 ± 0.07 (2) |
| IL/MO Frontal Ctx | Middle Frontal Ctx (A46) | 0.00 ± 0.00 (2) | 1.64 ± 0.04 (2) | 3.13 ± 0.12 (2) | 3.93 ± 0.07 (2) |
| **2° Motor Ctx** | **2° Motor Ctx (A6)** | 0.00 ± 0.00 (2) | 2.23 ± 0.10 (2) | 2.72 ± 0.05 (2) | 3.29 ± 0.09 (3) |
| **Primary Motor Ctx** | **Primary Motor Ctx (A4)** | 0.00 ± 0.00 (2) | 2.23 ± 0.10 (2) | 2.52 ± 0.10 (2) | 3.21 ± 0.13 (3) |
| **1° Sensory Ctx** | **1° Sensory Ctx (A3)** | 0.50 ± 0.00 (2) | 1.95 ± 0.00 (2) | 2.50 ± 0.00 (2) | 2.87 ± 0.00 (3) |
| **Posterior Parietal Ctx** | **Post Parietal Ctx (A7)** | 0.00 ± 0.00 (2) | 1.88 ± 0.14 (2) | 2.50 ± 0.13 (2) | 3.20 ± 0.06 (2) |
| **1° Auditory Ctx** | **1° Auditory Ctx (A41)** | 0.00 ± 0.00 (2) | 2.17 ± 0.14 (2) | 2.42 ± 0.08 (2) | 2.96 ± 0.02 (2) |
| **Temporal Assoc Ctx** | **Sup Temporal Ctx (A22)** | 0.00 ± 0.00 (2) | 2.11 ± 0.11 (2) | 2.54 ± 0.06 (2) | 2.81 ± 0.13 (2) |
| **1° Visual Cortex** | **1° Visual Ctx (A17)** | 0.00 ± 0.00 (2) | 1.84 ± 0.11 (2) | 2.55 ± 0.02 (2) | 2.66 ± 0.09 (2) |
| **Insular Ctx** | **Insular Ctx (A13-16)** | 0.00 ± 0.00 (2) | 0.72 ± 0.11 (2) | 2.16 ± 0.00 (2) | 2.56 ± 0.04 (3) |
| **Cingulate Cortex** | **Cingulate Cortex (A24)** | 0.00 ± 0.00 (2) | 1.50 ± 0.00 (2) | 2.77 ± 0.10 (2) | 3.48 ± 0.16 (3) |
| **All Cerebral Cortex** | **All Cerebral Cortex** | 0.05 ± 0.00 (2) | 1.83 ± 0.08 (2) | 2.64 ± 0.01 (2) | 3.17 ± 0.06 (3) |
| **PRL Frontal Ctx L2/3** | **A9 Layers 2/3** | 0.00 ± 0.00 (2) | 2.80 ± 0.05 (2) | 3.63 ± 0.13 (2) | 3.95 ± 0.05 (2) |
| **PRL Frontal Ctx L4** | **A9 Layer 4** | - | - | - | - |
| **PRL Frontal Ctx L5** | **A9 Layer 5** | 0.00 ± 0.00 (2) | 1.43 ± 0.08 (2) | 3.00 ± 0.00 (2) | 3.90 ± 0.10 (2) |
| **PRL Frontal Ctx L6** | **A9 Layer 6** | 0.00 ± 0.00 (2) | 1.43 ± 0.08 (2) | 3.13 ± 0.13 (2) | 3.95 ± 0.05 (2) |
| **IL/MO Frontal Ctx L2/3** | **A46 Layers 2/3** | 0.00 ± 0.00 (2) | 2.63 ± 0.13 (2) | 3.53 ± 0.23 (2) | 3.95 ± 0.05 (2) |
| **IL/MO Frontal Ctx L4** | **A46 Layer 4** | - | - | - | - |
| **IL/MO Frontal Ctx L5** | **A46 Layer 5** | 0.00 ± 0.00 (2) | 1.23 ± 0.08 (2) | 2.88 ± 0.13 (2) | 3.90 ± 0.10 (2) |
| **IL/MO Frontal Ctx L6** | **A46 Layer s6** | 0.00 ± 0.00 (2) | 1.08 ± 0.08 (2) | 3.00 ± 0.00 (2) | 3.95 ± 0.05 (2) |
| **2° Motor Ctx L2/3** | **A6 Layers 2/3** | 0.00 ± 0.00 (2) | 2.63 ± 0.13 (2) | 3.30 ± 0.04 (2) | 3.70 ± 0.10 (3) |
| **2° Motor Ctx L4** | **A6 Layer 4** | - | - | - | - |
| **2° Motor Ctx L5** | **A6 Layer 5** | 0.00 ± 0.00 (2) | 1.50 ± 0.00 (2) | 1.55 ± 0.05 (2) | 2.55 ± 0.10 (3) |
| **2° Motor Ctx L6** | **A6 Layer 6** | 0.00 ± 0.00 (2) | 2.58 ± 0.18 (2) | 3.30 ± 0.05 (2) | 3.62 ± 0.07 (3) |
| **1° Motor Ctx L2/3** | **A4 Layers 2/3** | 0.00 ± 0.00 (2) | 2.63 ± 0.13 (2) | 3.05 ± 0.05 (2) | 3.57 ± 0.15 (3) |
| **1° Motor Ctx L4** | **A4 Layer 4** | - | - | - | - |
| **1° Motor Ctx L5** | **A4 Layer 5** | 0.00 ± 0.00 (2) | 1.50 ± 0.00 (2) | 1.38 ± 0.13 (2) | 2.50 ± 0.14 (3) |
| **1° Motor Ctx L6** | **A4 Layer 6** | 0.00 ± 0.00 (2) | 2.58 ± 0.18 (2) | 3.13 ± 0.13 (2) | 3.57 ± 0.12 (3) |
| **1° Sensory Ctx L2/3** | **A1-3 Layers 2/3** | 0.00 ± 0.00 (2) | 2.63 ± 0.13 (2) | 3.13 ± 0.13 (2) | 3.42 ± 0.22 (3) |
| **1° Sensory Ctx L4** | **A1-3 Layer 4** | 2.00 ± 0.00 (2) | 1.68 ± 0.08 (2) | 2.38 ± 0.13 (2) | 2.65 ± 0.24 (3) |
| **1° Sensory Ctx L5** | **A1-3 Layer 5** | 0.00 ± 0.00 (2) | 0.88 ± 0.13 (2) | 1.38 ± 0.13 (2) | 1.98 ± 0.16 (3) |
| **1° Sensory Ctx L6** | **A1-3 Layer 6** | 0.00 ± 0.00 (2) | 2.63 ± 0.13 (2) | 3.13 ± 0.13 (2) | 3.42 ± 0.46 (3) |
| **Post Parietal Ctx L2/3** | **A7 Layers 2/3** | 0.00 ± 0.00 (2) | 2.55 ± 0.05 (2) | 3.13 ± 0.13 (2) | 3.68 ± 0.03 (2) |
| **Post Parietal Ctx L4** | **A7 Layer 4** | 0.00 ± 0.00 (2) | 1.48 ± 0.27 (2) | 2.38 ± 0.13 (2) | 2.85 ± 0.05 (2) |
| **Post Parietal Ctx L5** | **A7 Layer 5** | 0.00 ± 0.00 (2) | 0.88 ± 0.13 (2) | 1.38 ± 0.13 (2) | 2.50 ± 0.25 (2) |
| **Post Parietal Ctx L6** | **A7 Layer 6** | 0.00 ± 0.00 (2) | 2.63 ± 0.13 (2) | 3.13 ± 0.13 (2) | 3.78 ± 0.02 (2) |
| **1° Auditory Ctx L2/3** | **A41 Layers 2/3** | 0.00 ± 0.00 (2) | 2.90 ± 0.10 (2) | 3.13 ± 0.13 (2) | 3.58 ± 0.03 (2) |
| **1° Auditory Ctx L4** | **A41 Layer 4** | 0.00 ± 0.00 (2) | 1.75 ± 0.25 (2) | 2.13 ± 0.13 (2) | 2.45 ± 0.05 (2) |
| **1° Auditory Ctx L5** | **A41 Layer 5** | 0.00 ± 0.00 (2) | 1.13 ± 0.13 (2) | 1.13 ± 0.13 (2) | 2.03 ± 0.18 (2) |
| **1° Auditory Ctx L6** | **A41 Layer L6** | 0.00 ± 0.00 (2) | 2.90 ± 0.10 (2) | 3.30 ± 0.05 (2) | 3.78 ± 0.07 (2) |
| **Temporal Assoc Ctx L2/3** | **A22 Layers 2/3** | 0.00 ± 0.00 (2) | 2.63 ± 0.13 (2) | 3.18 ± 0.18 (2) | 3.38 ± 0.18 (2) |
| **Temporal Assoc Ctx L4** | **A22 Layer 4** | 0.00 ± 0.00 (2) | 1.68 ± 0.08 (2) | 2.35 ± 0.00 (2) | 2.35 ± 0.15 (2) |
| **Temporal Assoc Ctx L5** | **A22 Layer 5** | 0.00 ± 0.00 (2) | 1.13 ± 0.13 (2) | 1.30 ± 0.05 (2) | 1.83 ± 0.03 (2) |
| **Temporal Assoc Ctx L6** | **A22 Layer 6** | 0.00 ± 0.00 (2) | 3.03 ± 0.13 (2) | 3.35 ± 0.00 (2) | 3.68 ± 0.18 (2) |
| **1° Visual Ctx** | **A17 Layers 2/3** | 0.00 ± 0.00 (2) | 2.70 ± 0.10 (2) | 3.00 ± 0.00 (2) | 3.10 ± 0.10 (2) |
| **1° Visual Ctx** | **A17 Layer 4** | 0.00 ± 0.00 (2) | 1.70 ± 0.10 (2) | 1.80 ± 0.00 (2) | 1.80 ± 0.10 (2) |
| **1° Visual Ctx** | **A17 Layer 5** | 0.00 ± 0.00 (2) | 0.45 ± 0.15 (2) | 2.40 ± 0.00 (2) | 2.55 ± 0.05 (2) |
| **1° Visual Ctx** | **A17 Layer 6** | 0.00 ± 0.00 (2) | 2.50 ± 0.10 (2) | 3.00 ± 0.10 (2) | 3.20 ± 0.10 (2) |
| **Insular Ctx L2/3** | **A13-16 Layers 2/3** | 0.38 ± 0.00 (2) | 0.63 ± 0.15 (2) | 2.13 ± 0.00 (2) | 2.75 ± 0.03 (3) |
| **Insular Ctx L4** | **A13-16 Layer 4** | 0.19 ± 0.00 (2) | 1.33 ± 0.06 (2) | 2.63 ± 0.00 (2) | 3.25 ± 0.03 (3) |
| **Insular Ctx L5** | **A13-16 Layer 5** | 0.13 ± 0.00 (2) | 0.38 ± 0.06 (2) | 0.85 ± 0.00 (2) | 1.00 ± 0.03 (3) |
| **Insular Ctx L6** | **A13-16 Layer 6** | 0.75 ± 0.00 (2) | 1.25 ± 0.18 (2) | 1.88 ± 0.00 (2) | 2.38 ± 0.12 (3) |
| **All Cortex** | **Layers 2/3** | 0.00 ± 0.00 (2) | 2.44 ± 0.09 (2) | 3.09 ± 0.04 (2) | 3.45 ± 0.05 (2) |
| **All Cortex** | **Layer 4** | 0.33 ± 0.00 (2) | 1.43 ± 0.14 (2) | 2.15 ± 0.02 (2) | 2.50 ± 0.12 (2) |
| **All Cortex** | **Layer 5** | 0.00 ± 0.00 (2) | 1.12 ± 0.07 (2) | 1.87 ± 0.01 (2) | 2.59 ± 0.03 (2) |
| **All Cortex** | **Layer 6** | 0.00 ± 0.00 (2) | 2.20 ± 0.10 (2) | 3.06 ± 0.05 (2) | 3.50 ± 0.07 (2) |
| **Thalamus** | **Thalamus** | 0.00 ± 0.00 (2) | 0.63 ± 0.13 (2) | 2.13 ± 0.63 (2) | 2.75 ± 0.25 (2) |
| **PFN of Thalamus** | **PFN of Thalamus** | 0.00 ± 0.00 (2) | 1.33 ± 0.68 (2) | 2.63 ± 0.13 (2) | 3.25 ± 0.00 (2) |
| **Hypothalamus** | **Hypothalamus** | 0.00 ± 0.00 (2) | 0.38 ± 0.13 (2) | 0.85 ± 0.05 (2) | 1.00 ± 0.50 (2) |
| **PVN of Hypothalamus** | **PVN of Hypothalamus** | 0.00 ± 0.00 (2) | 1.25 ± 0.25 (2) | 1.88 ± 0.13 (2) | 2.38 ± 0.38 (2) |
| **Subthalamic Nucleus** | **Subthalamic Nucleus** | 0.00 ± 0.00 (2) | 0.21 ± 0.15 (2) | 0.63 ± 0.13 (2) | 1.88 ± 0.63 (2) |

### 11G2 immunolabeling of dSPNs versus iSPNs in Q175 Mouse

We used D1 immunolabeling to distinguish dSPNs (D1-rich) from iSPNs (D1-poor) (Lei e al., 2004; Deng et al., 2006), in conjunction with NeuN immunolabeling and NeuroTrace 435/455 staining to define the boundaries of neuronal nuclei and perikarya, and 11G2 immunolabeling. Surprisingly, we found that at 6 months of age, the presumptive dSPNs contained a 47.1% greater load of diffuse 11G2 by percent of cell area occupied than did the presumptive iSPNs, a difference that was significant for the 228 iSPNs and 228 dSPNs analyzed (Table 4). There was, however, no significant difference between presumptive dSPNs and presumptive iSPNs in the 11G2+ aggregate load as a percent of cell area at 6 months. The aggregate area was, nonetheless, slightly greater (12.4%) in D1-rich presumptive dSPN perikarya. By contrast, at 18 months of age, when aggregates have become more prominent, the presumptive dSPNs and presumptive iSPNs did not differ significantly in their diffuse 11G2 load (Table 4). In the case of aggregated 11G2 signal, the load at 18 months in D1-poor (iSPN) perikarya exceeded that in D1-rich (dSPN) perikarya by 11.4% (Table 4), but this difference was again not statistically significant. Thus, although a slightly higher 11G2 aggregate load may occur in presumptive iSPNs at this older age, at the younger age dSPNs show much greater enrichment in diffuse 11G2 signal.

**Table 4.** Age-wise HTT1A accumulation in dSPNs and iSPNs of Q175 mice relative to vulnerability. The table shows 11G2 mutant HTT1a load for dSPNs versus iSPNs at 6 and 18 months of age in Q175 mice, 2 mice per age group, in comparison to the loss of the dSPN marker PPT versus the iSPN marker PPE by ISHH in Q175 mice at the same ages as assessed for 11G2 load, and compared to the reported loss of dSPNs versus iSPNs at grade 1 HD in humans. ISHH data from Deng et al., 2021, and HD data from Deng et al. (2004).

|  |  | Q175 | Q175 | Q175 Loss |  | HD Gr1 Loss |
| --- | --- | --- | --- | --- | --- | --- |
| SPN Type | Q175 Age | Agg Area | Diffuse Area | % < Cntrl | SPN Type | % < Cntrl |
| dSPN | 6 mo | 2.78% ± 0.14% | 7.64% ± 0.35% | 5.0% | dSPN | 29.1% |
| iSPN | 6 mo | 2.47% ± 0.12% | 5.19% ± 0.28% | 15.3% | iSPN | 66.4% |
| dSPN | 18 mo | 2.79% ± 0.15% | 1.72% ± 0.12% | 7.0% | dSPN | 29.1% |
| iSPN | 18 mo | 3.11% ± 0.15% | 1.69% ± 0.10% | 25.5% | iSPN | 66.4% |

### P90 Labeling in Q175 and WT Mouse Cerebral Cortex, Hippocampus, Amygdala and Diencephalon

In our initial screening, we also found that both the 11G2 and 1B12 antibodies detected mutant HTT1a accumulation in pallial structures (which include cerebral cortex, hippocampus and much of amygdala) and diencephalic structures of 2-, 6-, 10-, 12-, and 18-month old Q175 mice. The regional patterns appeared identical, but the labeling was slightly more intense for 11G2. Accordingly, the 11G2 antibody was used to prepare 2-3 complete series of sections through forebrain and midbrain for mapping purposes, which are also largely used for the following description and exclusively used for the illustrations. In cerebral cortex of mutants, we observed prominent diffuse nuclear immunolabeling in layer 4 neurons of somatosensory cortex at 2 months of age, as also reported by Smith et al. (2023) using the S830 antibody, and by 6 months we saw widespread diffuse nuclear immunolabeling of neurons across the cortical layers (Fig. 4B-C). The immunolabeling, however, was less prevalent in layer 5 than the other layers, and less common in layer 5B than layer 5A. Nuclear aggregates and neuropil aggregates became yet more abundant stepwise with further aging of Q175 mice out to 18 months (Fig. 4D). We performed multiple immunofluorescence with NeuroTrace 435/455 to determine the cellular localization of 11G2+ immunolabeling. We found that 11G2 signal at 6-12 months of age in Q175 mice accumulates in cortex largely as aggregates in the nuclei of cortical neurons in layers 2-4 and 6, but largely in the neuropil of layer 5 (Fig. 4E-H).

**Figure 4.**
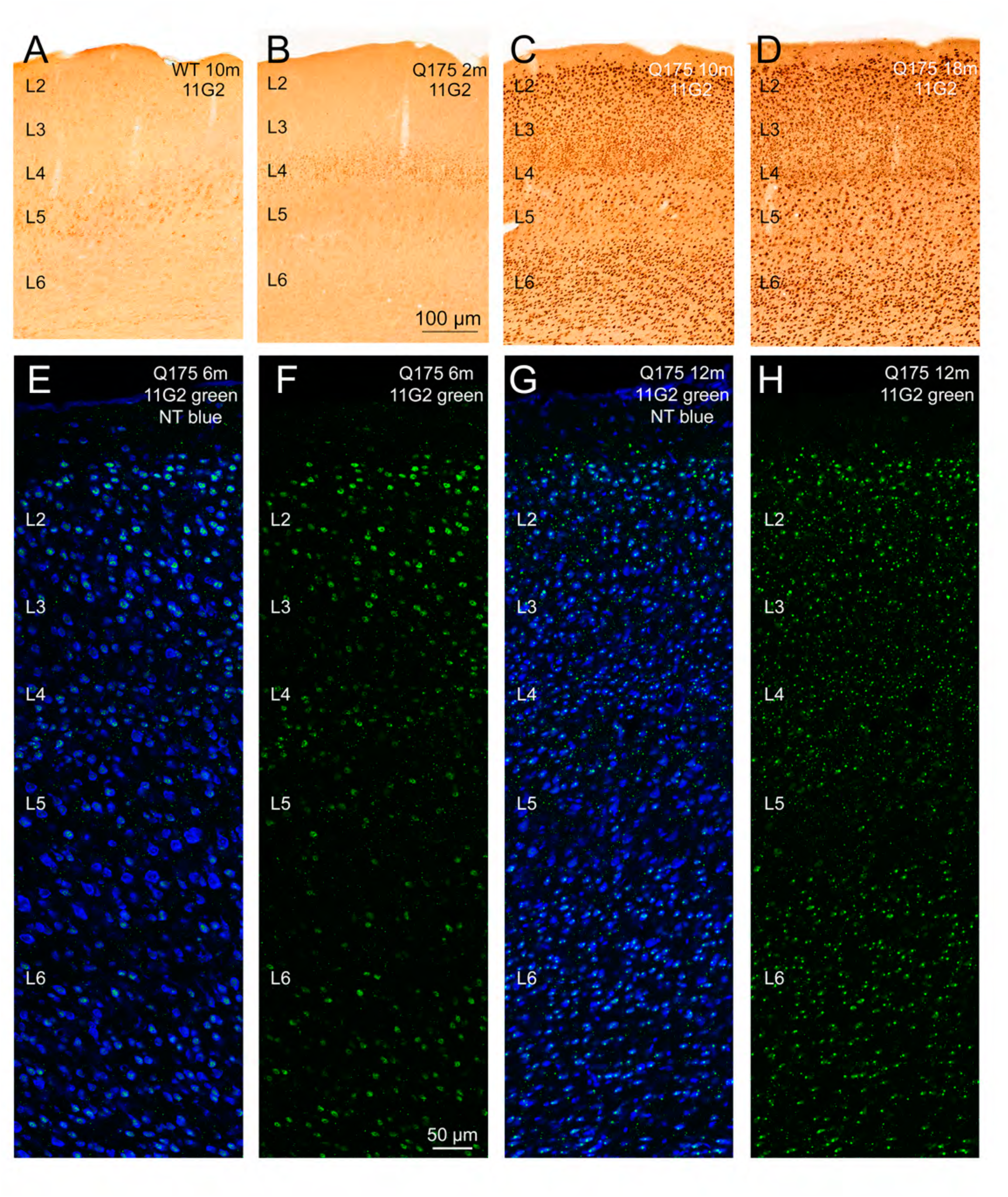
The first row of images shows P90 immunolabeling with the 11G2 antibody in 2-month old (B), 10-month old (C), and 18-month old Q175 (D) mice compared to a 10-month old WT littermate control (A) throughout the depth of somatosensory cortex, using DAB labeling. 11G2 immunolabeling of perikarya in the Q175 mice at 2 months manifests mainly as diffuse nuclear labeling of layer 4 neurons. Subsequently, diffuse nuclear labeling appears in neurons of most layers followed by nuclear and neuropil aggregate formation. Images E and G show a merged CLSM image through the depth of somatosensory cortex from a 6-month old Q175 mouse and a 12-month old Q175 mouse, respectively, with striatal neurons labeled blue with NeuroTrace (NT) and 11G2 immunolabeling in green. Images F and H show the 11G2 immunolabeling alone for these fields of view. The CLSM images show diffuse and aggregated 11G2 signal in some cortical neurons at 6 months, with nuclear and neuropil aggregates more prominent at 12 months. Note, however, that image A shows that 11G2 immunolabeling is present in the cytoplasm of many layer 5 cortical neurons in WT mice. Scale bar in B applies to images in first row, and scale bar in F to images in second row.

As in the case of basal ganglia structures, in our study of WT cortex, we saw P90+ immunolabeling that resembled what we have seen with antibodies against WT huntingtin in WT mice (i.e., Q175 WT littermates) – in this case evident immunolabeling of the cytoplasm of layer 5 cortical neurons (Figs. 1A, 2E, 4A) (Fusco et al., 1999; Dragatsis et al., 2018). Neither nuclear nor aggregate labeling was, however, seen in WT cortex, hippocampus, amygdala or diencephalon, unlike in mutant mice. Additionally, we saw similar but fainter cytoplasmic P90 immunolabeling of layer 5 cortical neurons in 2-month old Q175 mice, as well as the aforementioned nuclear labeling of layer 5 neurons of somatosensory cortex (Figs. 1B, 2F, 4B). Labeling of the cytoplasm of layer 5 neurons was not evident in older Q175 mice, and instead the nuclear labeling occurred (Fig. 4C, D).

In the Q175 hippocampal complex, diffuse nuclear P90+ immunolabeling was seen in CA1 pyramidal neurons at 2 months, and dentate gyrus granule cells had slightly above WT background levels of diffuse nuclear signal, but neuropil labeling was absent (Fig. 1B). In the amygdalar complex, P90+ signal was largely absent at 2 months, except for neuropil labeling in the central amygdala (CEA) of both WT and Q175 mice (not shown). A few neurons with cytoplasmic P90+ signal were also present in CEA of both WT and Q175 mice. Immunolabeling in the majority of neurons in the thalamus and hypothalamus was largely at background levels at 2 months in Q175 mice (Fig. 1B). Although light diffuse perikaryal labeling was evident in the paraventricular nucleus of the hypothalamus (PVN) of mutants (Fig. 5E), similar labeling was seen in 2-month old WT mice (Fig. 5A). By 6 months of age, hippocampal pyramidal neuron labeling in the cornu ammonis (CA) field of Q175 mice had become much more prominent, with neurons of CA1, CA3 and CA4 but not CA2 highly enriched in 11G2+ signal (not shown). The labeling was largely nuclear, with diffuse labeling and aggregates present in seemingly every CA1 neuron cell body, but only diffuse nuclear labeling evident in CA3 and CA4. Labeling of dentate gyrus granule cells in Q175 mice was more prominent than at 2 months, consisting of intense diffuse nuclear labeling, and punctate neuropil labeling in the dentate hilus was also prominent. In the amygdalar complex at 6 months (Fig. 6A), immunolabeling in WT mice resembled that at 2 months, while prominent diffuse nuclear immunolabeling was now evident in neurons of the CEA and basolateral amygdala (BLA) of Q175 mice, more so the CEA (not shown). Very light nuclear immunolabeling was seen in the cortical, basomedial, and medial amygdala of Q175 mice (not shown). Immunolabeling in thalamus of Q175 mice was also advanced beyond that at 2 months, with diffuse nuclear immunolabeling evident in neurons of the posterior, lateral posterior, midline thalamic nuclei, and in the parafascicular nucleus (PFN) (not shown). P90 immunolabeling of hypothalamus of Q175 mice in general remained sparse, but the diffuse cytoplasmic PVN labeling seen at 2 months of age remained evident in WT mice (Fig. 5B), and in Q175 mice diffuse nuclear labeling of PVN neurons was now evident (Fig. 5F). By 10-12 months, intense diffuse nuclear labeling and intensely labeled nuclear aggregates were seen in all neurons of the dentate gyrus (granule cell layer and hilus both), CA1, CA3, and CA4 of Q175 mice (Figs. 1C, 7). Labeling in CA2, by contrast, was mainly diffusely nuclear (Fig. 7C,F). In the amygdala, immunolabeling in WT mice at 10-12 months was unchanged from 6 months, but in Q175 mice, diffuse nuclear immunolabeling, nuclear aggregates, and neuropil aggregates were commonplace, most notably in CEA, BLA, and basomedial amygdala (BMA) (Fig. 6B). The thalamus of Q175 mice by 10-12 months showed widespread diffuse labeling of neuronal nuclei, again mainly in the posterior, lateral posterior, midline thalamic nuclei, and PFN (Fig. 1C). At this age, the PVN of Q175 hypothalamus was densely packed with neurons with diffuse nuclear signal in mutants (Fig. 5G), while in WT PVN the immunolabeling was as it was at 2 and 6 months (Fig. 5C). The overall regional labeling in hippocampus and amygdala was similar at 18 months in Q175 mice as at 12 months, but with nuclear aggregates larger where they were present at 10-12 months and now accompanied by neuropil aggregates as well, and regions previously showing mainly diffuse nuclear signal now also showing nuclear aggregates (Fig. 1D). Thalamic labeling in Q175 mice at 18 months was pervasive, with all thalamic cell groups now showing at least intense diffuse nuclear labeling (Fig. 1D), and PFN among the more enriched, in keeping with its high huntingtin expression (Allen Mouse Brain Atlas, 2004).

**Figure 5.**
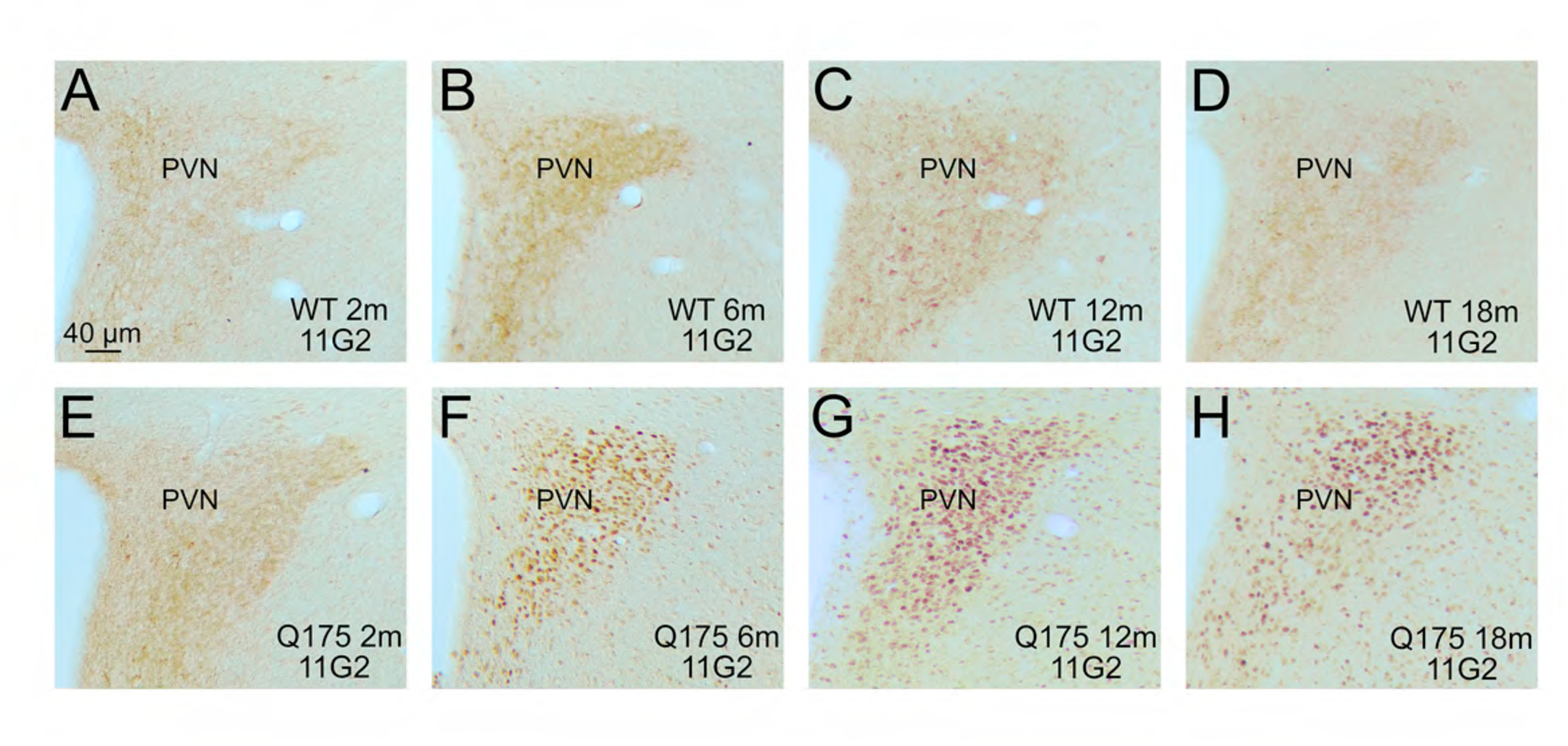
The first row of images shows P90 immunolabeling in the right paraventricular nucleus of the hypothalamus (PVN) with the 11G2 antibody in 2-month old (A), 6-month old (B), 12-month old Q175 (C), and 18-month old WT (D) mice, while the second row shows the PVN immunolabeling for 11G2 in 2-month old (E), 6-month old (F), 12-month old (G), and 18-month old Q175 (H) mice. Diffuse cytoplasmic immunolabeling and some neuropil immunolabeling makes the PVN stand out from the surrounding hypothalamus in WT mice. Similar immunolabeling occurs in 2-month Q175 mice (E), but by 6 months and thereafter PVN neurons in Q175 mice show distinct diffuse nuclear 11G2 (F-G). Scale bar in A applies to all images.

**Figure 6.**
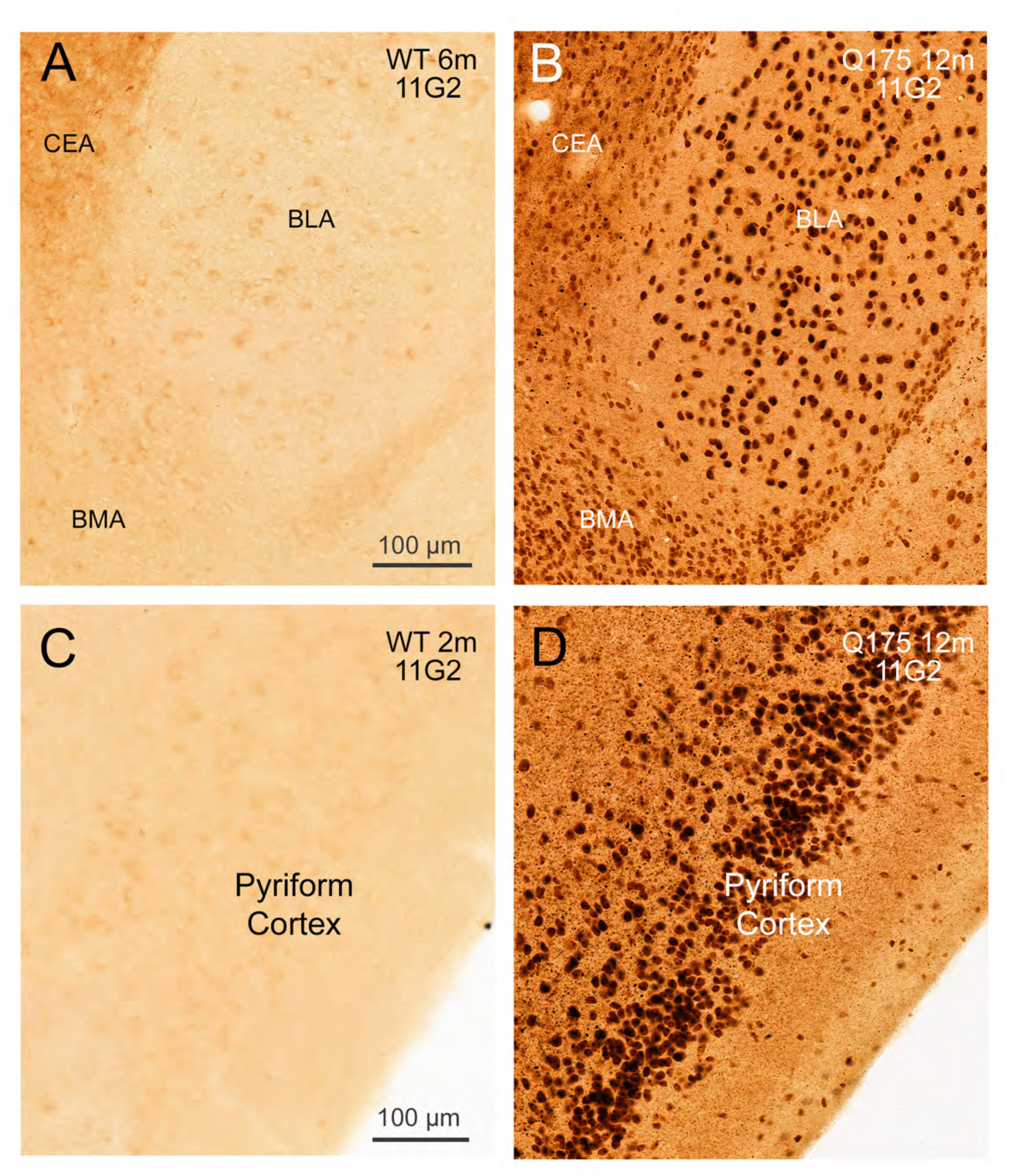
The first row of images shows P90 immunolabeling with the 11G2 antibody in a 12-month old Q175 (B) mouse compared to a 6-month old WT littermate control (A), in the basolateral (BLA), basomedial (BMA), and central (CEA) amygdaloid nuclei of the right side of the brain. Note that in the WT mouse, light cytoplasmic 11G2 immunolabeling of some perikarya in BLA and BMA is evident, while distinct neuropil immunolabeling is evident in CEA. By contrast, in the 12-month old Q175 mouse, dense nuclear 11G2+ signal is evident in many neurons of these three amygdaloid cell groups. The second row of images shows 11G2 immunolabeling of pyriform cortex in a 12-month old Q175 (D) mouse compared to a 2-month old WT littermate control (C). Note that WT pyriform cortex is largely devoid of distinct 11G2+ signal, with some perikarya perhaps with very light cytoplasmic immunolabeling bordering on background level. By contrast, in the 12-month old Q175 mouse, dense nuclear 11G2+ signal is evident in seemingly all pyriform cortex neurons, with neuropil aggregates evident as well. Scale bar in A applies to B as well, and scale bar in C applies to D as well.

**Figure 7.**
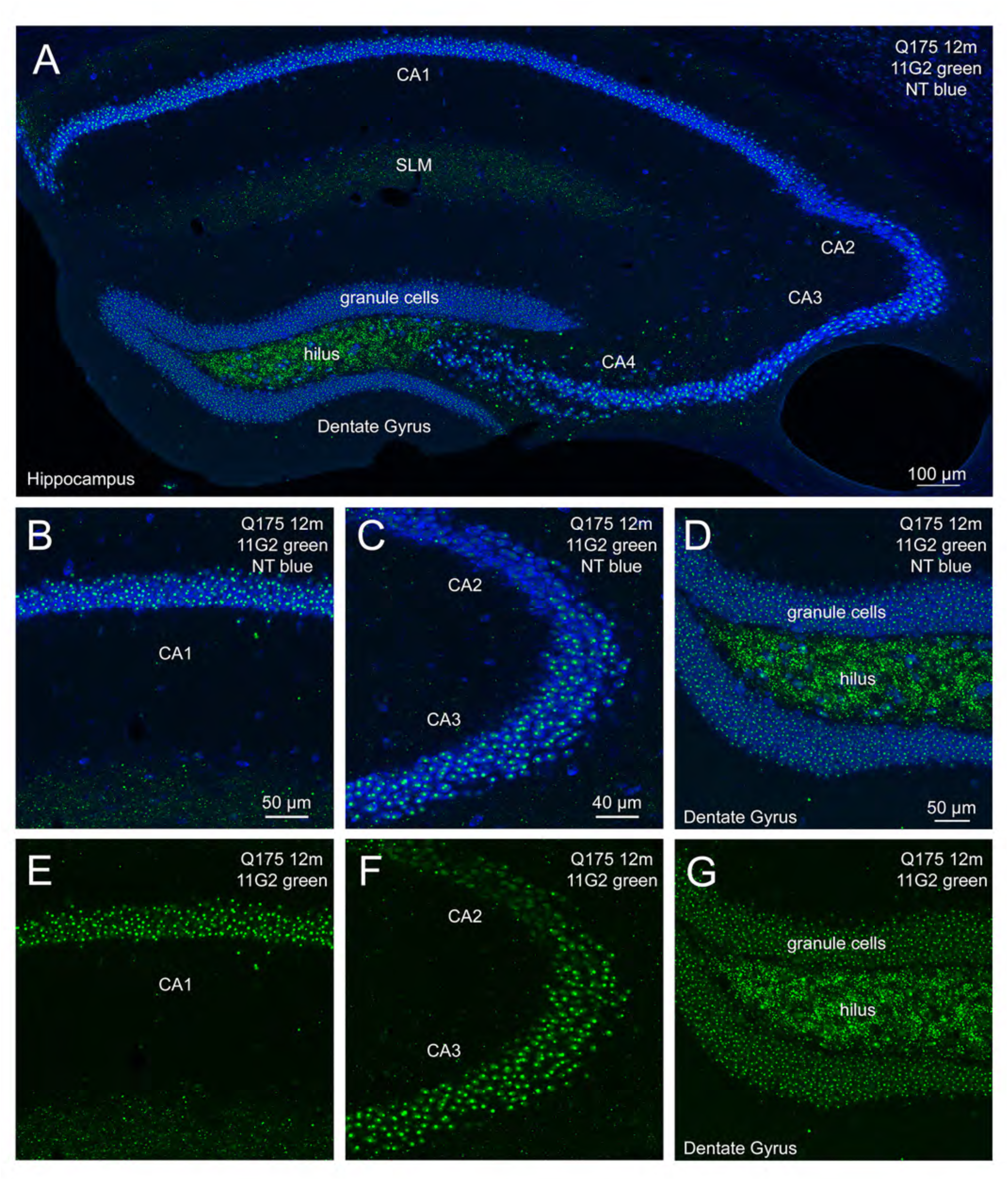
The images show P90 immunolabeling with the 11G2 antibody in the right-side hippocampus of a 12-month old Q175 mouse. Image A shows hippocampus with neurons labeled blue with NeuroTrace (NT) and 11G2 immunolabeling in green. Images B, C, and D show merged CLSM images of CA1, CA2/CA3, and dentate gyrus, respectively, with neurons labeled blue with NT and green for 11G2. Images E, F and G show the 11G2 immunolabeling alone for these same fields of view, respectively. The CLSM images show diffuse and aggregated nuclear 11G2 signal in CA1, CA3, CA4, and dentate granule cell layer neurons, but mainly diffuse nuclear signal in CA2. The hilus is neuron sparse, and its labeling is mainly in neuropil. Scale bar in B apples also to E, scale bar in C applies also to F, and scale bar in D applies also to G.

Hypothalamus in Q175 mice was poor in P90+ signal at 18 months (Fig. 1D), in keeping with its low huntingtin expression (Allen Mouse Brain Atlas, 2004), with PVN being an exception, as its neurons possessed diffuse nuclear immunolabeling in mutants (Fig. 5H). Notable from 6 months on was the intense labeling of pyriform cortex neurons in Q175 mice, beginning as mainly diffuse nuclear signal at 6 months and progressing to the presence of large aggregates in seemingly all pyriform cortex neurons, as well as in neuropil (Fig. 1C, D, 6D). By contrast, in WT mice immunolabeling of pyriform cortex appeared to be at background levels (Fig. 6C). We scored the 11G2 immunolabeling in Q175 mice on a 0-4 scale at 2-, 6-, 12- and 18-months of age as shown in Table 3. In multiple immunofluorescence studies with NeuroTrace 435/455 counterstain, we observed that both 11G2+ aggregates and diffuse labeling were sparse in PARV+ cortical interneurons in Q175 mice (Fig. 8). Finally, we found that the 11G2+ neuropil aggregates that predominate in layer 5 of cortex at 10-12 months of age in Q175 mice did not align with SMI32+ processes in cortex (Fig. 9).

**Figure 8.**
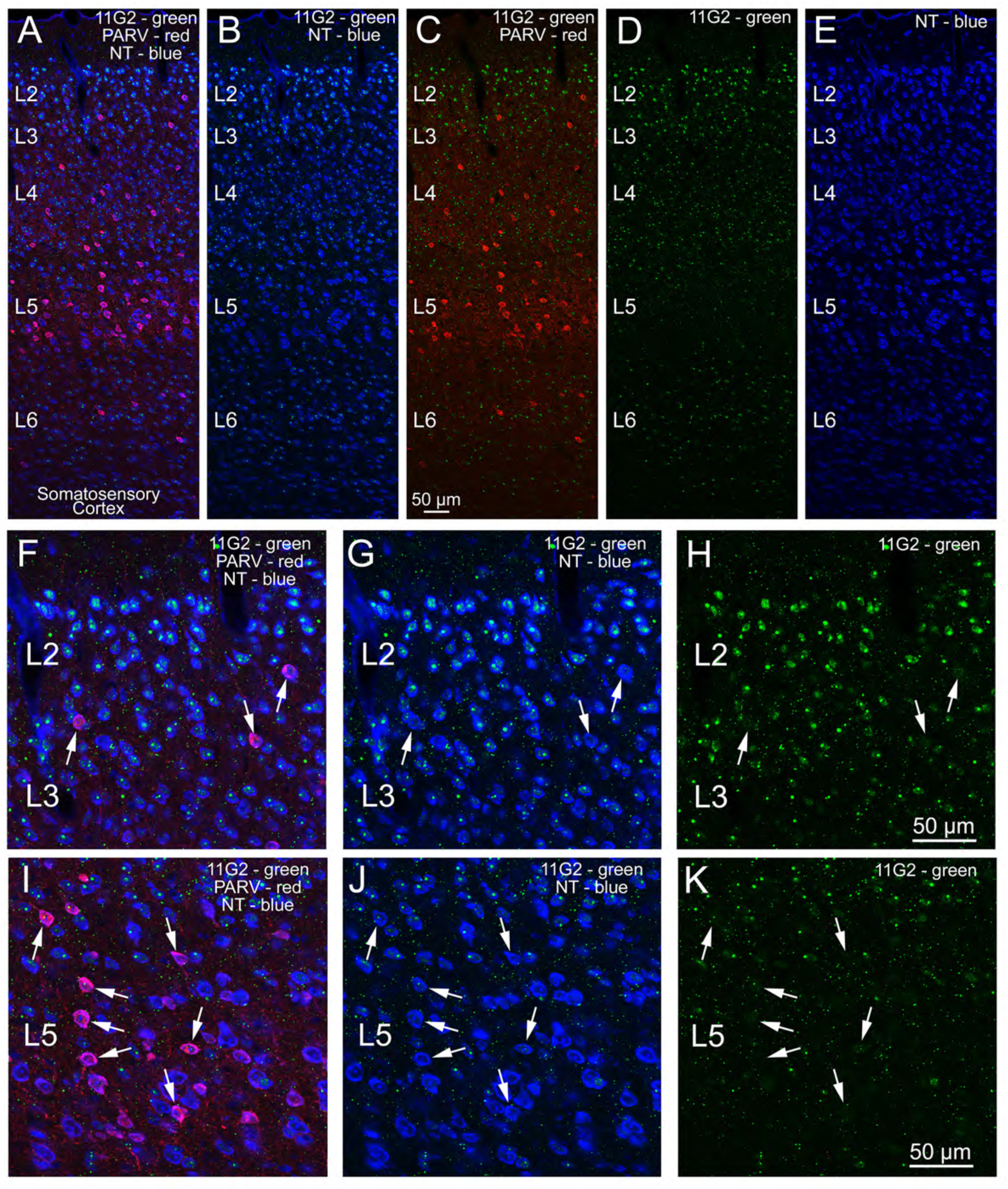
The first row shows a single CLSM field of view of the entire depth of somatosensory cortex from a 10-month old Q175 mouse with cortical neurons labeled blue with NeuroTrace (NT), 11G2+ aggregates shown in green, and PARV+ interneurons shown in red, with the panels A-C each showing the labeling in different combinations, and D and E showing the 11G2 and NT signal alone, respectively. The second and third rows shows higher power images of parts of the same field of view as in the first row. The first column of the second and third rows shows the merged 11G2, PARV, and NT signals for layer 3 (F) and layer 5 (I), respectively, with arrows in each row indicating some of the PARV+ interneurons. The subsequent columns show the 11G2 and NT signals (G, J), and the 11G2-only signal (H, K), with arrows indicating the same interneurons as in the first column. In general, 11G2+ signal is sparse in PARV+ cortical interneurons. Scale bar in C applies to images in first row, scale bar in H applies to images in second row, and scale bar in K applies to images in third row.

**Figure 9.**
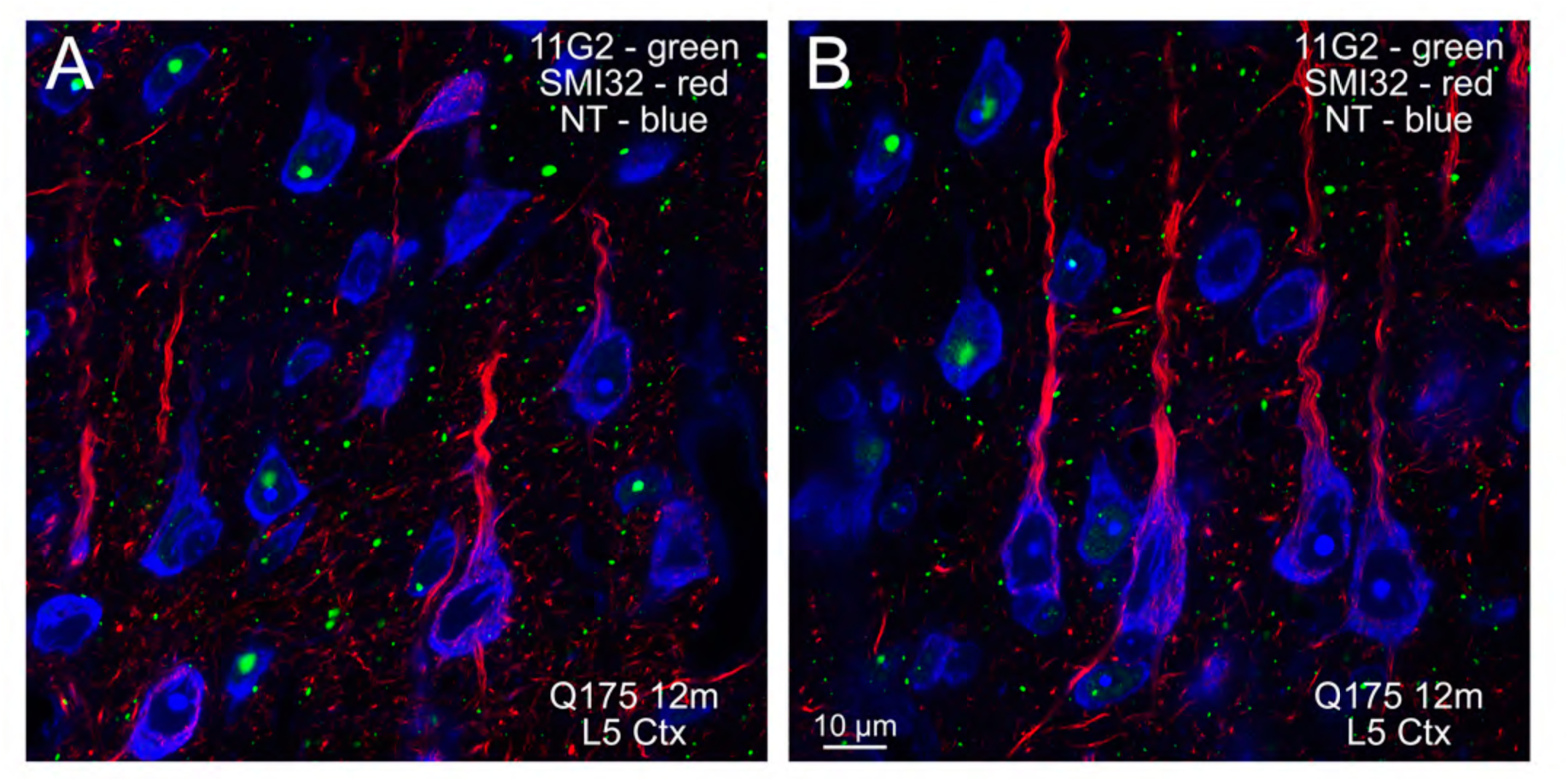
The images show a merged CLSM image of layer 5 (L5) of somatosensory cortex from a 12-month old Q175 mouse with cortical neurons labeled blue with NeuroTrace (NT), 11G2+ aggregates shown in green, and the pyramidal neuron dendrite marker SMI32 shown in red. Although 11G2+ aggregates not localized to neuronal nuclei are present (i.e., neuropil aggregates), none are localized to the SMI32+ dendrites. Thus, neuropil aggregates appear to be localized to terminals or SMI32-negative dendrites in Q175 cortex. Scale bar in B applies to both images.

### P90 Immunolabeling in R6/2 Mouse Basal Ganglia, Cerebral Cortex, Hippocampus, Amygdala and Diencephalon

We examined the R6/2 mouse because its transgene specifically produces mutant human HTT1a. We found that the 11G2 and 1B12 P90 antibodies both detected mutant huntingtin accumulation in the five 9-12 week-old R6/2 mice examined, with the 11G2 signal slightly more intense. In either case, the immunolabeling in R6/2 mice was prominent and pervasive, with all or nearly all neurons throughout brain, including cortex, hippocampus, amygdala, somatic striatum, limbic striatum, thalamus and hypothalamus showing dense nuclear labeling (Fig. 10B, C, E, F). Many neurons in GPe, GPi and SNr also were intensely labeled. Nuclear aggregates were not obvious in the DAB-labeled material, but small neuropil aggregates were. Immunofluorescence labeling for 11G2 in combination with NeuroTrace 435/455 counterstain, however, revealed that nuclear aggregates are present in striatal neurons in R6/2 mice, but apparently obscured in the DAB-labeled material by the dense nuclear labeling (Fig. 10G, H). Small neuropil aggregates were evident in both the DAB-labeled and immunofluorescence tissue. In the DAB-labeled tissue, we additionally saw moderate diffuse immunolabeling of small cell bodies in white matter that we assume are oligodendrocytes. Note that although earlier studies have not commonly detected mutant huntingtin aggregates in R6/2 glia (Davies et al., 1997), more recent studies have detected them in R6/2, Q175, and HdhQ150 mice (Jansen et al., 2017). As noted above, the immunolabeling in cortex, hippocampus, amygdala, thalamus and hypothalamus was also highly intense in R6/2 mice, with neurons in these regions showing dense labeling throughout their nucleus (Fig. 10B, C). Any nuclear aggregates that might be present were again obscured in the DAB-labeled tissue by the dense nuclear labeling. Immunofluorescence labeling for 11G2 in combination with NeuroTrace 435/455 staining, however, again revealed that nuclear aggregates are present in cortical neurons in R6/2 mice, with small neuropil aggregates present in cortex as well (Fig. 11).

**Figure 10.**
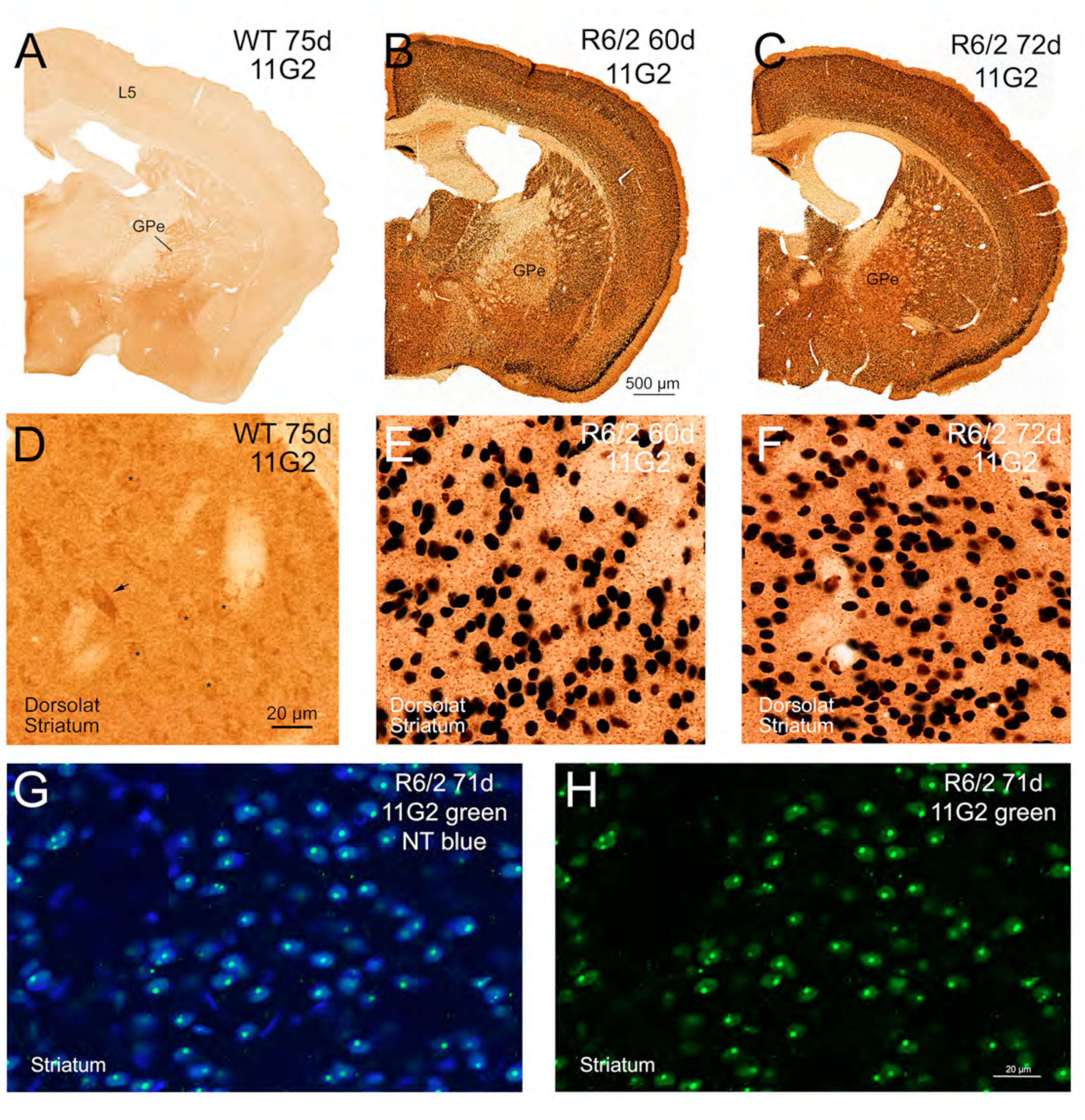
The first two rows of images show P90 immunolabeling with the 11G2 antibody in 60-day old (B, E) and 72-day old R6/2 (C, F) mice compared to a 75-day old WT littermate control (A, D), with the first row showing the right side of the forebrain at the level of GPe, and the second row showing a view of dorsolateral striatum. 11G2 immunolabeling of perikarya in the R6/2 mice at both ages is intense and widespread, and appears to be nuclear. Images G and H show a merged CLSM image of striatum from a 71-day old R6/2 mouse with striatal neurons labeled blue with NeuroTrace and 11G2 immunolabeling in green, and an image of the 11G2 immunolabeling alone, respectively. Note that these images show that the dense immunolabeling of seemingly all individual striatal neurons in E and F reflects intense diffuse nuclear labeling plus a large nuclear aggregate obscured in E and F by the intense nuclear labeling. Note, however, that image A shows that 11G2 immunolabeling is present in terminals in GPe and in the cytoplasm of perikarya in cortical layer 5 in WT mice. The higher power view in image D of WT striatum shows that light cytoplasmic 11G2 immunolabeling is present in numerous presumptive striatal projections neurons in WT mouse (some indicated by small asterisks), with intense 11G2 signal present in scattered large neurons the size of cholinergic interneurons (one indicated by an arrow). Scale bar in B applies to images in first row, scale bar in D applies to images in second row and scale bar in H applies to images in third row.

**Figure 11.**
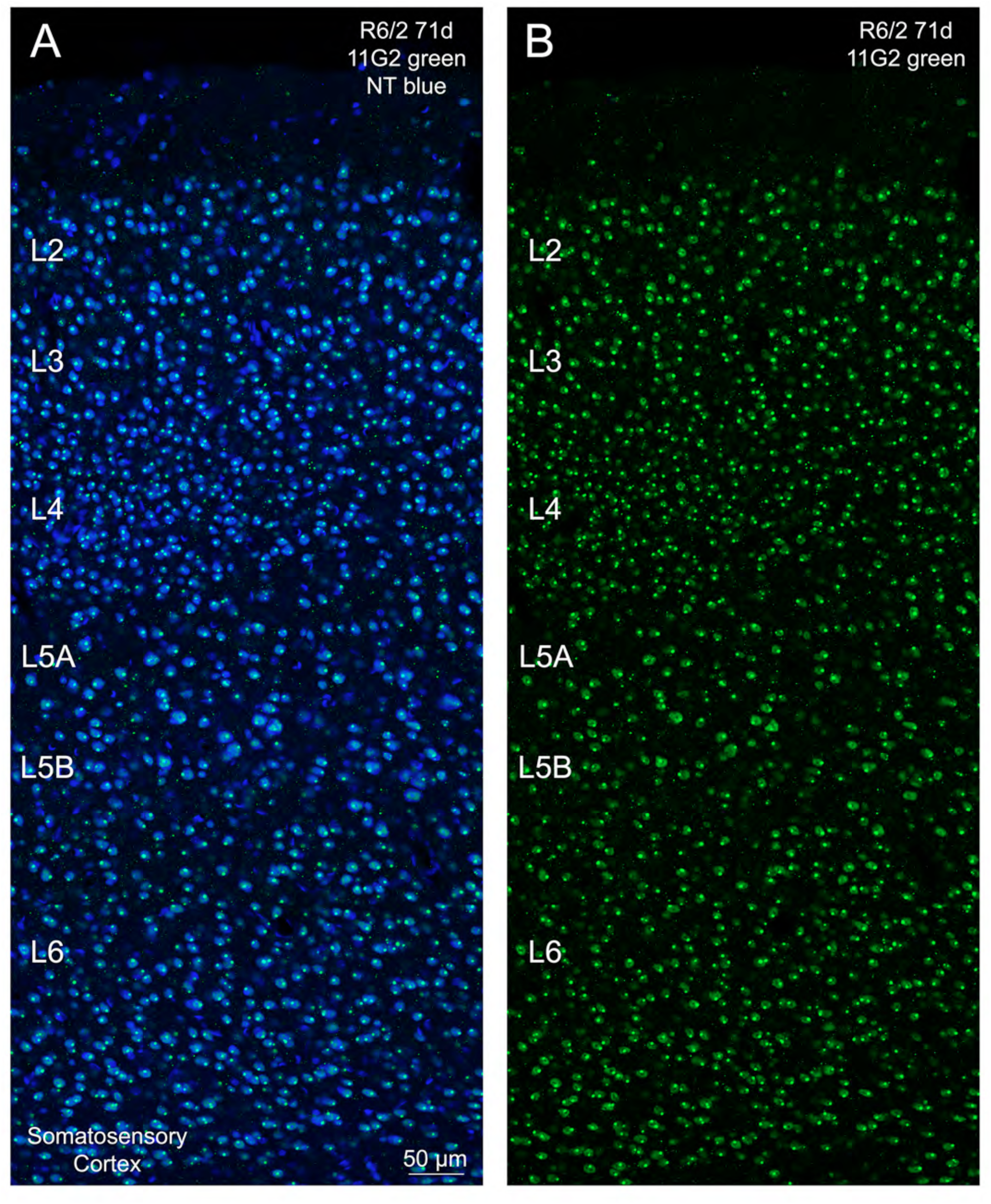
Images A and B show a merged CLSM image of somatosensory cortex from a 71-day old R6/2 mouse with cortical neurons labeled blue with NeuroTrace and 11G2 immunolabeling in green, and an image of the 11G2 immunolabeling alone, respectively. Note that these images show that the dense immunolabeling of seemingly all individual cortical neurons in R6/2 mice reflects intense unaggregated nuclear labeling plus a large nuclear aggregate obscured in the DAB-labeled tissue by the intense nuclear labeling. Scale bar in A applies to both images.

As in the case of the WT littermates of Q175 mice, we saw immunolabeling in the four WT littermates of R6/2 mice examined in the cytoplasm of layer 5 cortical neurons and presumptive cholinergic striatal interneurons, and in likely SPN terminals in GPe, GPi, SNr, and ventral pallidum (Figs. 10A, D, 12A). Very light cytoplasmic P90 immunolabeling was also seen in WT mouse SPNs (Fig. 10D). Like in WT mice from Q175 litters, we also saw neuropil immunolabeling in CEA and PVN, but only background immunolabeling of pyriform cortex perikarya (Fig. 12C-E). Such labeling was not seen in R6/2 mice (Figs. 10B, C, E, F, 12B).

**Figure 12.**
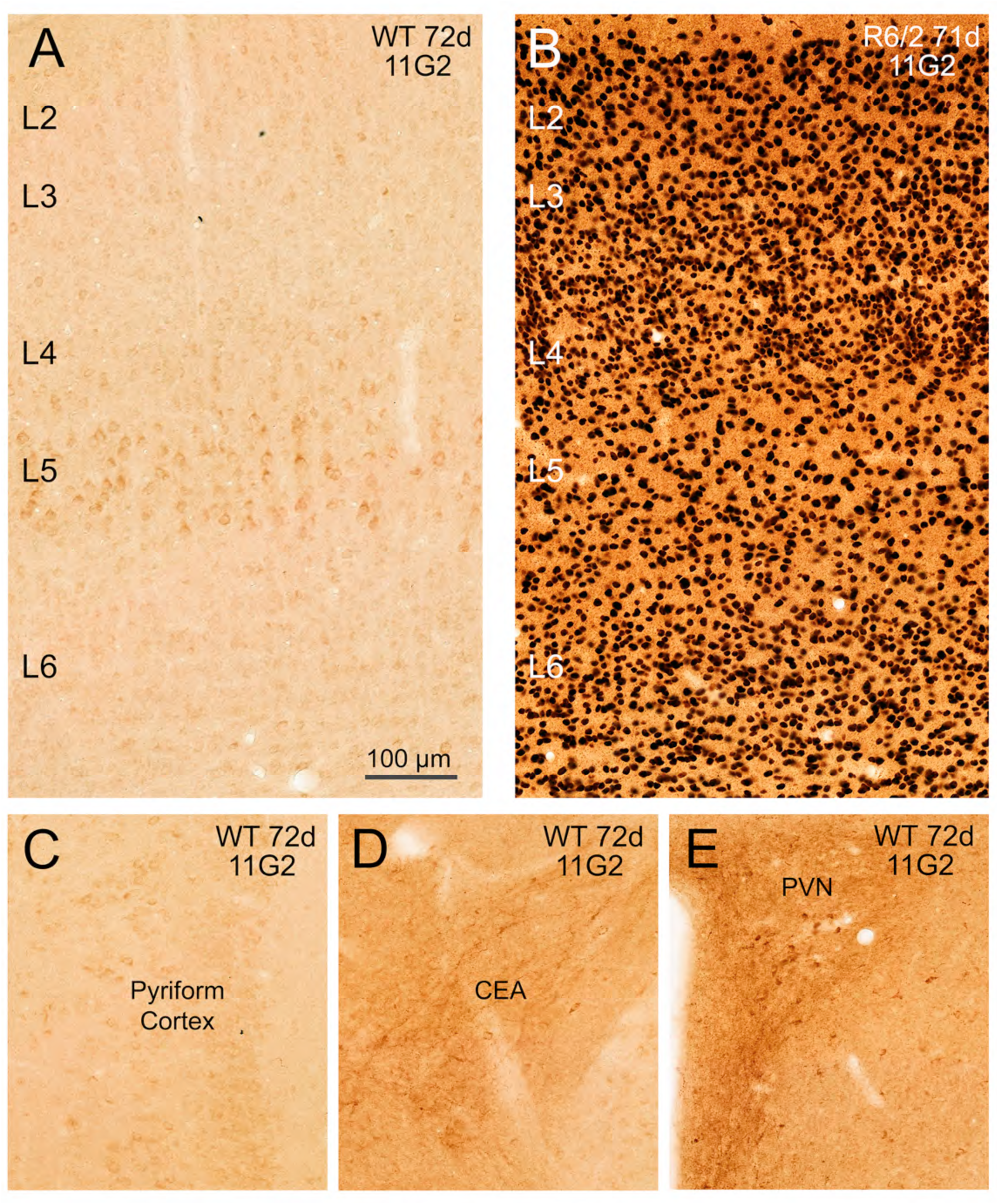
The first row of images shows 11G2 immunolabeling in the full depth of somatosensory cortex in a 71-day old R6/2 (B) mouse compared to a 72-day old WT littermate control (A). Note that WT layer 5 contains numerous perikarya with distinct 11G2+ cytoplasmic immunolabeling, while R6/2 cortex at about the same age has distinct 11G2+ nuclear immunolabeling of seemingly all neurons throughout the depth of cortex. Figure 11 shows that the R6/2 nuclear immunolabeling seen in DAB-labeled material comprises both diffuse nuclear signal and a large aggregate. The second row of images shows 11G2 immunolabeling in pyriform cortex (C), CEA (D), and PVN (E) on the right side of the brain in a 72-day old WT mouse. Note that WT pyriform cortex is largely devoid of distinct 11G2+ signal, with some perikarya perhaps with very light cytoplasmic immunolabeling. By contrast, neuropil immunolabeling is evident in CEA and PVN. The scale bar in A applies to all images.

These results again raise two possibilities – either WT mice produce a detectible level of HTT1a in some neurons from their endogenous mouse huntingtin (which is identical to human HTT1a for the last 8 amino acids at the C-terminus), or these neural structures possess some other antigen to which the P90 antibodies bind, either an extended form of HTT1a to which the P90 antibodies unexpectedly bind or a protein with antigenic determinants similar to the C-terminal octapeptide of HTT1a and that has the same distribution as huntingtin.

### Specificity of P90 Antibodies in Mice

We immunolabeled sections from N171-Q82 mice at 4 months of age with the P90 antibodies, when they are known to form mutant protein aggregates (Schilling et al., 1999). Since N171-Q82 mice do not produce mutant HTT1a, the N171-Q82 mice serve as negative controls for the specificity of the antibodies for the P90 neo-epitope in the context of a polyglutamine expansion to HD-causing length. Our immunolabeling confirmed that there was no diffuse nuclear or aggregated P90 signal in N171-Q82 brain, while robust PHP1 immunolabeling of aggregates was observed in these same mice, representing detection of the aggregated N171 mutant protein (Fig. 13). This result suggests that the diffuse nuclear immunolabeling, and the nuclear and neuropil aggregate immunolabeling observed with the P90 antibodies in Q175 and R6/2 mice represents detection of mutant human HTT1a, but not a mutant huntingtin form with amino acids beyond the final C-terminal proline of HTT1a. We saw similar results with the MW8 antibody as with the P90 antibodies, confirming the findings of Smith et al. (2023) and Landles et al. (2021), which indicate that MW8 too does not appear to detect a form of the huntingtin protein that extends beyond the HTT1a amino acid sequence.

**Figure 13.**
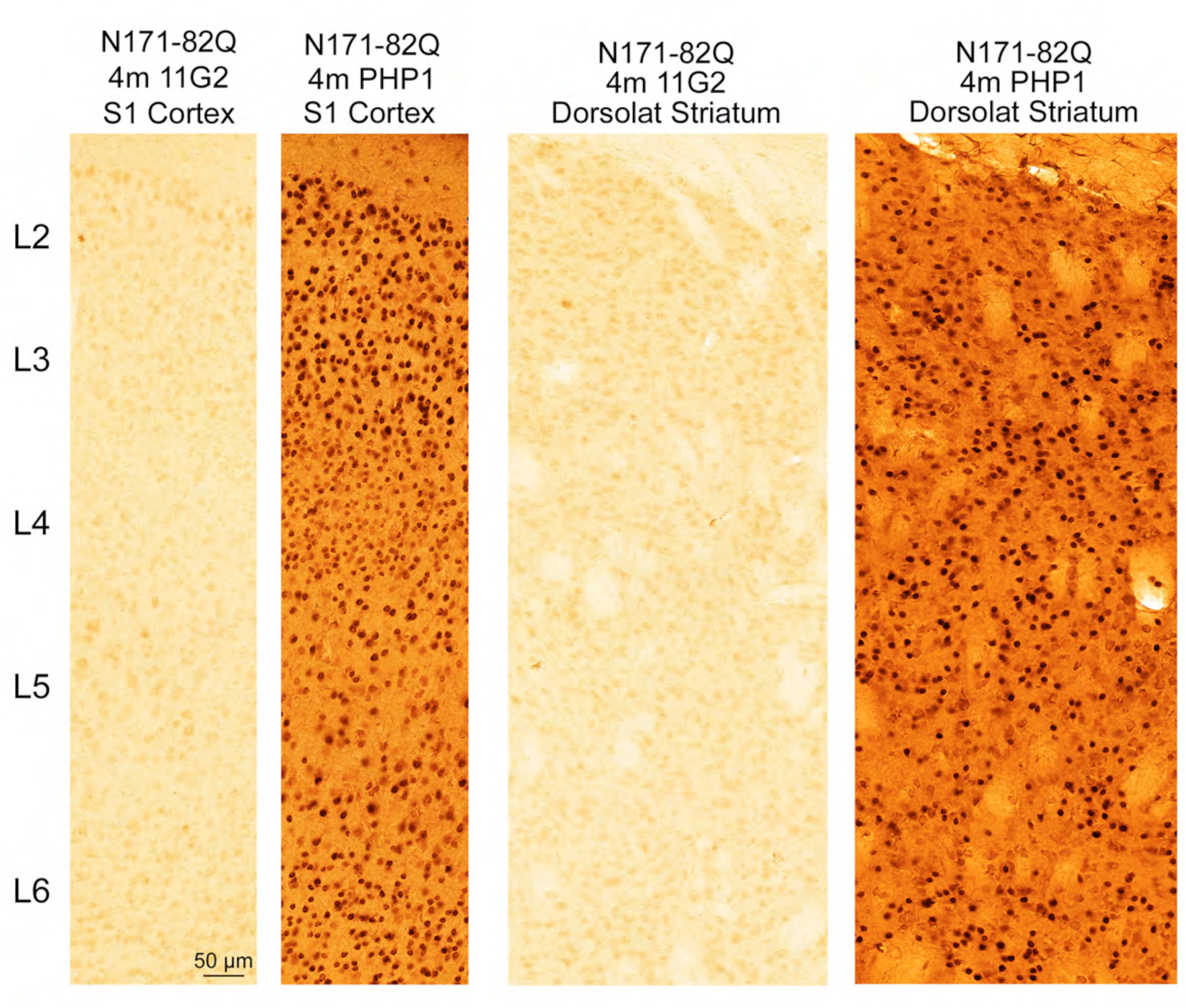
The images compare 11G2 immunolabeling and PHP1 immunolabeling in cortex and striatum of a 4-month old N171-82Q mouse. No 11G2 immunolabeling of aggregates was present in the cortex, striatum (A, C), or any other brain area in the N171-82Q mouse. By contrast, widespread immunolabeling of aggregates is seen in cortex and striatum with PHP1 (B, D). Scale bar in A applies to all images.

Note that N171-82Q mice also produce WT huntingtin from their endogeneous mouse huntingtin genes, and thus might be expected to yield P90 immunolabeling like we saw in WT littermates of Q175 and R6/2 mice. We did, in fact, see P90 immunolabeling in CEA in N171-82Q mice resembling what we saw in WT littermates of Q175 and R6/2 mice (Supplementary Fig. 1), but we did not see clearcut P90 immunolabeling in GPe fibers, large striatal neurons, or layer 5 cortical perikarya like we had in the WT littermates of Q175 and R6/2 mice. As the N171-82Q mice had been perfusion fixed at JHU and shipped to us, we thought it possible that these factors had diminished the antigenicity of the tissue and thus contributed to the absence of clearcut P90 immunolabeling in GPe fibers, large striatal neurons, or layer 5 cortical perikarya. We, thus, subjected the N171-82Q tissue to our standard antigen retrieval protocol involving heating in basic Tris-EDTA buffer (Jiao et al., 1999; Deng et al., 2021). Following this, we saw clear 1B12 and 11G2 immunolabeling like in WT mice in the cytoplasm of layer 5 cortical neurons, fiber immunolabeling in GPe, and cytoplasmic immunolabeling in large striatal neurons, as well as intensified immunolabeling in CEA (Supplementary Fig. 1). Thus, N171-Q82 mice too produce an antigen detected by 11G2 and 1B12 with the same regional distribution as WT huntingtin.

To further assess the specificity of the P90 antibodies, we performed immunolabeling of tissue from 2- and 12-month old WT and Q175 mice, and 10-week old WT and R6/2 mice, with 11G2 having been pre-incubated for 24 hours with a many fold excess of the P90 target peptide AEEPLHRP. We found that the immunolabeling we otherwise saw in WT mice and 2-month Q175 mice in the cytoplasm of layer 5 neuronal perikarya, in the cytoplasm of large striatal neurons, in presumptive SPN terminals in striatal target areas (Fig. 14A-D; Supplementary Fig. 2), and in the neuropil of PVN and CEA was prevented. Moreover, nuclear and aggregate immunolabeling in Q175 mice was prevented or markedly reduced by blocking 11G2 binding in tissue by preadsorption with exogeneous AEEPLHRP (Fig. 14E-L; Supplementary Figs. 3-5).

**Figure 14.**
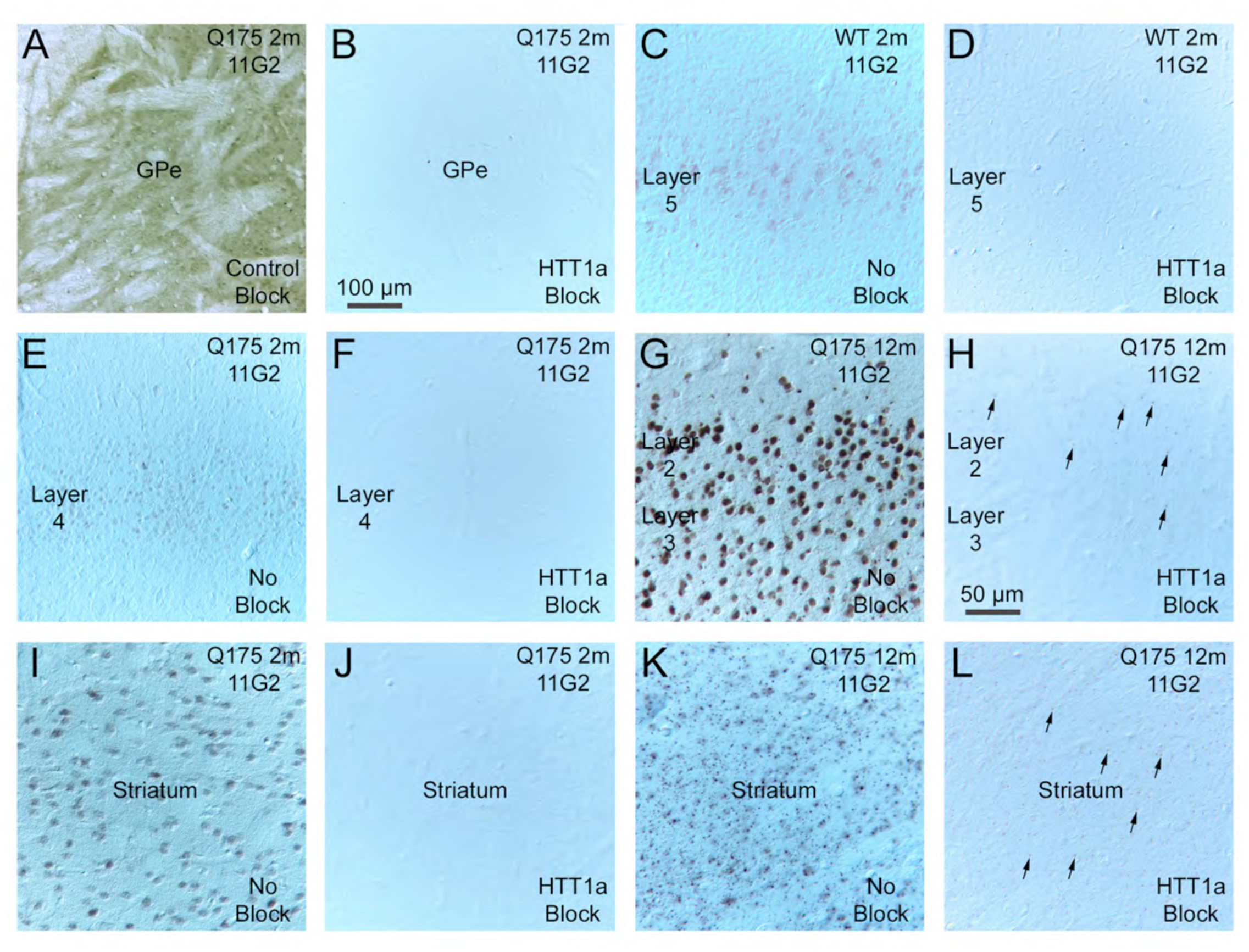
Images showing that 11G2 immunostaining in WT and Q175 mouse brain sections is blocked by HTT1a target octapeptide (AEEPLHRP). Images A and B show the robust immunostaining of GPe fibers in 2-month old Q175 mouse with 11G2 (A) co-incubated with non-target peptide (YHRLLACLQNVHKVTTC) and the absence of immunostaining when 11G2 (B) is co-incubated with HTT1a target peptide (AEEPLHRP). Medial is to the left. Images C and D show the cytoplasmic immunostaining of cortical layer 5 neuronal perikarya in 2-month old WT mouse (Q175 littermate) with 11G2 (C) and the absence of layer 5 neuronal immunostaining when 11G2 (D) is co-incubated with HTT1a target peptide. Images E and F show the nuclear immunostaining of cortical layer 4 neurons in 2-month old Q175 mouse with 11G2 (E) and the absence of immunostaining when 11G2 (F) is co-incubated with HTT1a target peptide. Images G and H show the widespread diffuse immunostaining of cortical neuron nuclei and of nuclear and neuropil aggregates in layers 2 and 3 of 12-month old Q175 mouse with 11G2 (G) and the near absence of immunostaining when 11G2 (H) is co-incubated with HTT1a target peptide, with only scattered small aggregates remaining present (some indicated by arrows). Images I and J show the diffuse nuclear immunostaining of striatal neuronal perikarya in 2-month old Q175 mouse with 11G2 (I) and the absence of striatal immunostaining when 11G2 (D) is co-incubated with HTT1a target peptide. Images K and L show the widespread diffuse immunostaining of neuronal nuclei and of nuclear and neuropil aggregates in striatum of 12-month old Q175 mouse with 11G2 (G) and the near absence of immunostaining when 11G2 (H) is co-incubated with HTT1a target peptide, with only scattered small aggregates remaining present (some indicated by arrows). Images were taken using differential interference Nomarski optics to enhance labeling contrast. Scale bar in B applies to A-F, and that in H applies to G-L.

By contrast, blocking with control peptide (YHRLLACLQNVHKVTTC) did not prevent the 11G2 immunolabeling at any age, although background labeling was increased (Fig. 14A; Supplementary Figs. 2-5). Blocking reduced but did not eliminate P90 immunolabeling in R6/2 mice (Supplementary Figs. 6-7). We attribute this outcome to the extreme enrichment of R6/2 brain in HTT1a, since antibody binding is a competitive reaction and the abundant endogeneous HTT1a may have thus outcompeted the exogeneous AEEPLHRP. Blocking did, however, eliminate the P90 immunolabeling in the WT littermates of R6/2 mice.

Because blocking 11G2 with its preferred target, the C-terminal octapeptide of HTT1a, might also prevent off-target binding to similar antigens, the blocked control does not necessarily rule out the possibility that P90 signal in WT mice occurred due to binding to an antigen other than HTT1a that also possesses the AEEPLHRP sequence (or a highly similar one), despite the evidence for the HTT1a selectivity of 11G2 (Missineo et al., 2024). To assess this, we examined tissue from mice with tamoxifen-driven huntingtin deletion, and thus deletion of any possible WT HTT1a, and control conditional mice without deletion (Bragg et al., 2024). In mice with a 90% huntingtin deletion, as demonstrated in prior studies (Bragg et al., 2024) and our own immunolabeling with the D7F7 antibody against full-length huntingtin, we saw substantially diminished 11G2 immunolabeling of layer 5 cortical neurons, striatal neurons, and SPN terminals in striatal target areas. The remaining 11G2 signal may stem from residual neurons without huntingtin deletion, due to incompleteness and mosaicism in the HTT knockdown (Fig. 15), or to a contribution of an undiminished HTT1a mimic to the labeling. The issue of HTT1a production from WT alleles will be considered further in the Discussion.

**Figure 15.**
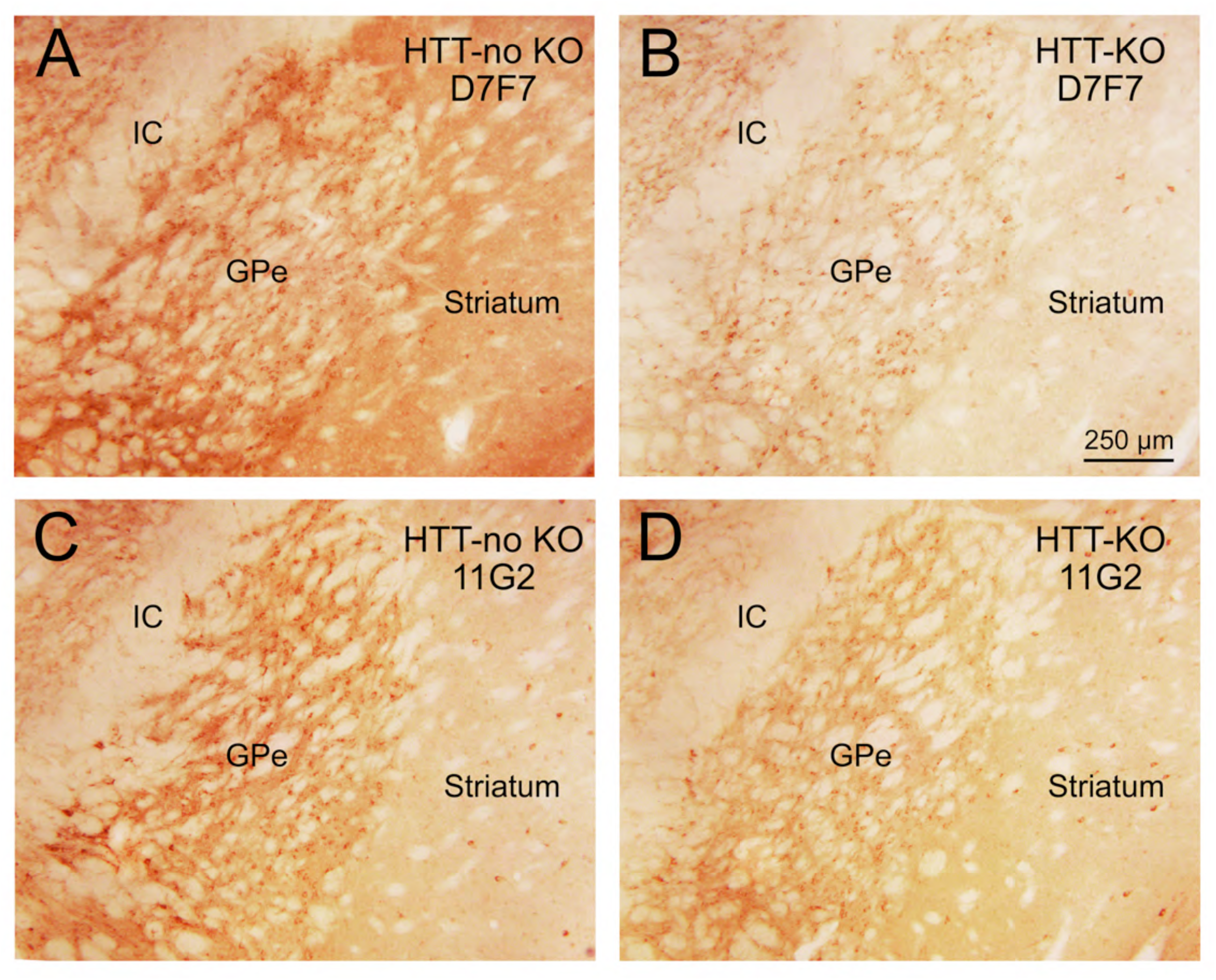
Immunolabeling in striatum at the level of right GPe for D7F7 (A, B) and 11G2 (C, D) in huntingtin*-*flox mice with a tamoxifen inducible Cre recombinase, i.e. Httflox/flox;CAG-CreERT (Bragg et al., 2024), without tamoxifen treatment (A, C) and with tamoxifen treatment (B, D). The mice were 3 months old and one month post vehicle or tamoxifen treatment. In the mice with vehicle treatment, and thus with normal levels of WT huntingtin, both the D7F7 antibody against full-length huntingtin (A) and 11G2 (C) yielded immunolabeling of terminals in GPe, presumably of SPN origin. In the tamoxifen-treated mouse, the D7F7 immunolabeling of GPe fibers was substantially reduced but not entirely eliminated, consistent with prior evidence that WT huntingtin levels are only 90% reduced by tamoxifen treatment (Bragg et al., 2024). In the same mouse, 11G2 immunolabeling of GPe fibers was also substantially reduced but not completely eliminated (D), suggesting that 11G2 in WT mouse detects some level of production of HTT1a from WT huntingtin. Scale bar in B applies to all images. Abbreviation: internal capsule – IC.

### P90 Signal Overlap with MW8 and PHP1 in Q175 Mice

To gain insight into the relationship of P90 signal to the mutant huntingtin signal with other antibodies, we used multiple immunofluorescence to compare 11G2, MW8 and PHP1 in 6-month old, 12-month old, and 18-month old Q175 mice. We found that at six months the MW8+ signal occupies only the center of 11G2+ aggregates in striatum, and diffuse nuclear labeling was present but not as salient as for 11G2 (Figure 16A-F). The aggregate area occupied by PHP1 signal at six months was even less, and diffuse nuclear labeling was again less salient than for 11G2. By 12 months, both MW8 signal and PHP1 signal occupied more of the central 11G2+ area of each aggregate, but still not its entirety (Fig. 16G-L). Diffuse nuclear labeling was much reduced for 11G2 but still present, but again much less salient for MW8 and PHP1. These trends continued out to 18 months, with MW8 and PHP1 signal filling more of the 11G2+ area of each aggregate, but never its entirety. Similar results were obtained for cerebral cortex at 12 and 18 months for the three antibodies, when aggregate immunolabeling for MW8 and PHP1 first becomes noteworthy. As for striatum, the MW8+ and PHP1+ area of any given aggregate was less than for 11G2 and occupied the center of the 11G2+ aggregate (Fig. 17).

**Figure 16.**
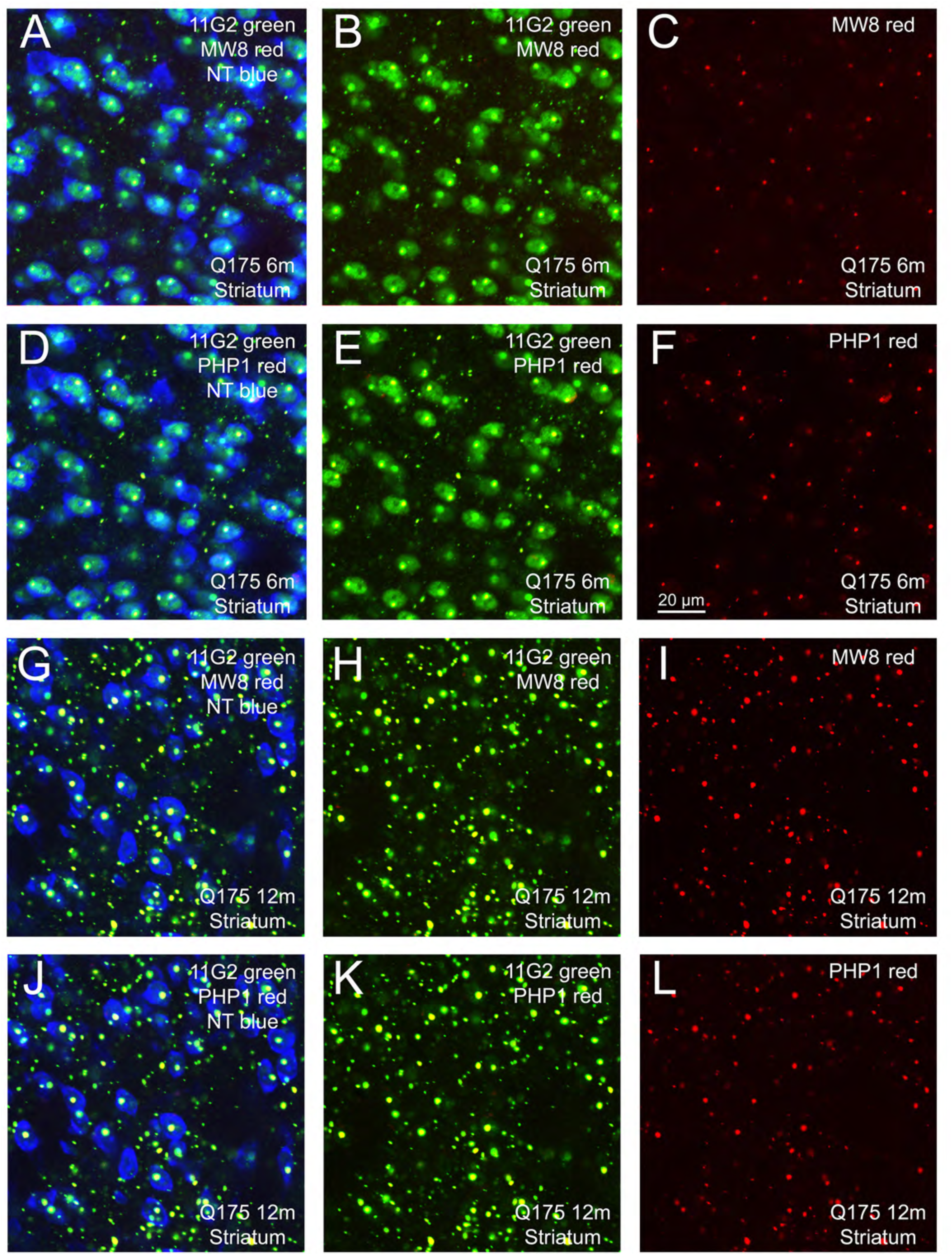
The first two rows show a CLSM image of a single field of view of striatum from a 6-month Q175 mouse from tissue that had been simultaneously immunolabeled for 11G2, MW8 and PHP1, while the last two rows show a CLSM image of a single field of view of striatum from a 12-month Q175 mouse from tissue that had been simultaneously immunolabeled for 11G2, MW8 and PHP1. The first and third row show in succession the merged CLSM image for 11G2 (green), MW8 (red) and NeuroTrace (blue) in the first column, the merged 11G2 and MW8 image in the second column, and the MW8-only image in the third column. The second and fourth row show in succession the merged CLSM image for 11G2 (green), PHP1 (red) and NeuroTrace (blue) in the first column, the merged 11G2 and PHP1 image in the second column, and the PHP1-only image in the third column. Note that at 6 months the MW8+ signal occupies only the center of 11G2+ aggregates, and diffuse nuclear labeling is not prominent for MW8. The aggregate area occupied by PHP1 signal at six months is slightly greater, but diffuse nuclear labeling is again meager. By 12 months, both the MW8 signal and the PHP1 signal occupy more of the 11G2+ area of each aggregate, but still not its entirety. Scale bar in F applies to all images.

**Figure 17.**
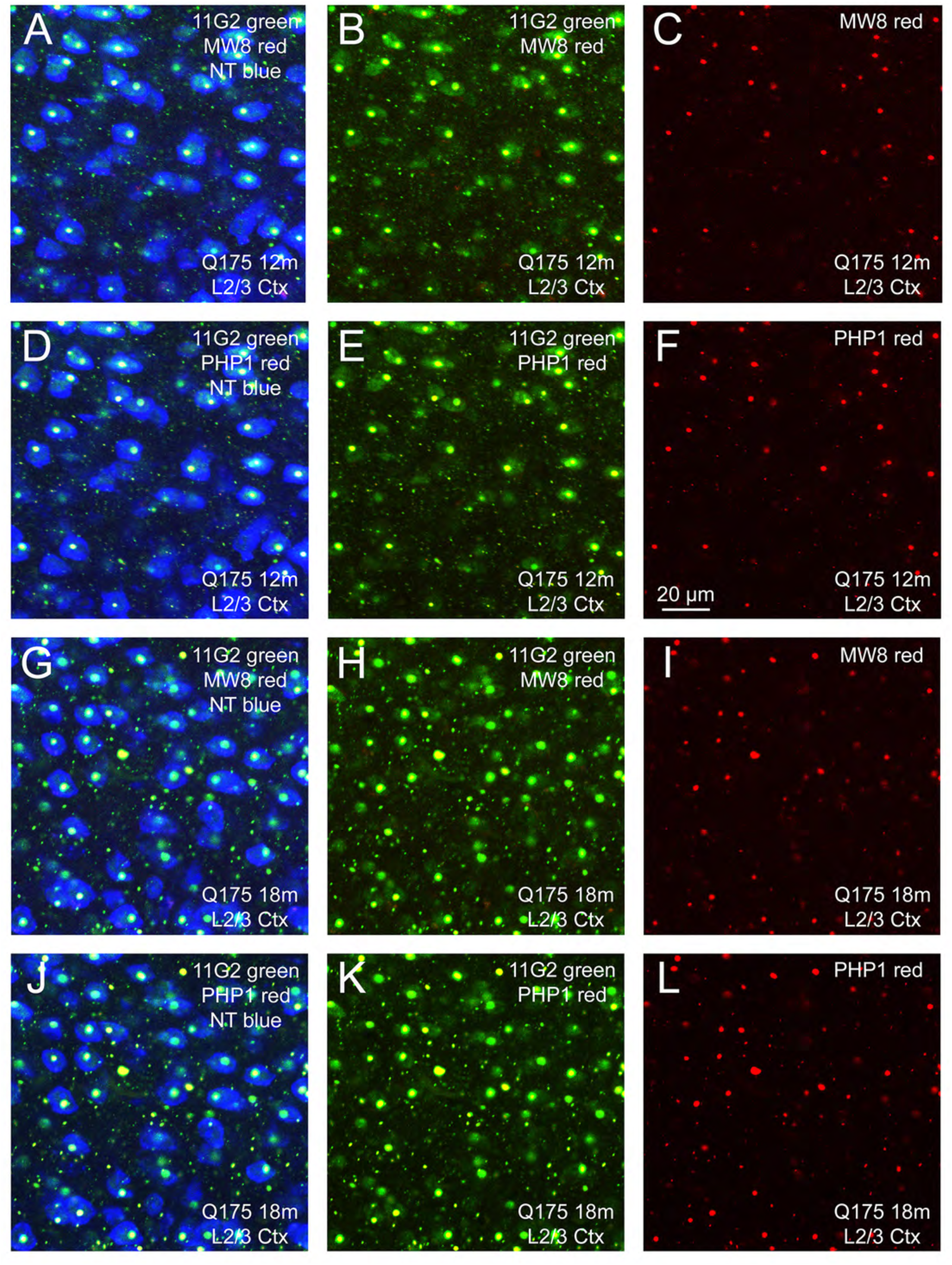
The first two rows show a CLSM image of a single field of view of layers 2 and 3 of somatosensory cortex from a 12-month Q175 mouse from tissue that had been simultaneously immunolabeled for 11G2, MW8 and PHP1, while the last two rows show a CLSM image of a single field of view of striatum from an 18-month Q175 mouse from tissue that had been simultaneously immunolabeled for 11G2, MW8 and PHP1. The first and third row show in succession the merged CLSM image for 11G2 (green), MW8 (red) and NeuroTrace (blue) in the first column, the merged 11G2 and MW8 image in the second column, and the MW8-only image in the third column. The second and fourth row show in succession the merged CLSM image for 11G2 (green), PHP1 (red) and NeuroTrace (blue) in the first column, the merged 11G2 and PHP1 image in the second column, and the PHP1-only image in the third column. Note that at 12 months the MW8+ signal occupies only the center of 11G2+ aggregates, and diffuse nuclear labeling is not prominent for MW8. The aggregate area occupied by PHP1 signal at 12 months is slightly greater, but diffuse nuclear labeling is again meager. By 18 months, both MW8 signal and PHP1 signal occupy more of the 11G2+ area of each aggregate, but still not its entirety. Scale bar in F applies to all images.

### P90 Labeling in Human Basal Ganglia

We found that both the 11G2 and 1B12 antibodies detected aggregates in HD human basal ganglia and cortex. Control brain did not label for these, or for PHP1 or MW8. Because the 11G2 yielded more robust labeling than the 1B12, we used immunofluorescence with it in combination with NeuroTrace to assess the localization of 11G2 in caudate and putamen. The 11G2 immunolabeling was mainly observed in neuropil aggregates in grade 1 through grade 4 HD striatum, with aggregates less common at grade 4 (Fig. 18). Nucleus accumbens was slightly richer than caudate and putamen. In our DAB-labeled tissue, we observed 11G2+ aggregates in the neuropil of GPe, GPi, SNr, and VP, but fewer than in striatum. Blocking 11G2 with an excess of exogeneous AEEPLHRP (the target of 11G2 within HTT1a) prevented aggregate immunolabeling in HD basal ganglia, while blocking with control peptide did not (Supplementary Fig. 8). The regional 11G2 localization scores for basal ganglia are shown in Table 5. We also assessed 11G2 immunolabeling in caudate and putamen in combination with immunofluorescence for PHP1 and MW8 (Figs. 19, 20). 11G2 tended to detect many but not all of the same aggregates as PHP1. Moreover, for any given aggregate detected by both, the aggregate area was less with 11G2 and more centrally located. In the case of MW8, the overlap with 11G2 was more complete, which is not surprising since both are directed at the C-terminus of HTT1a, while PHP1 is directed at its proline-rich domain. MW8+ aggregates that were 11G2-negative were seen, however. Aggregates that only labeled with 11G2 were rare.

**Figure 18.**
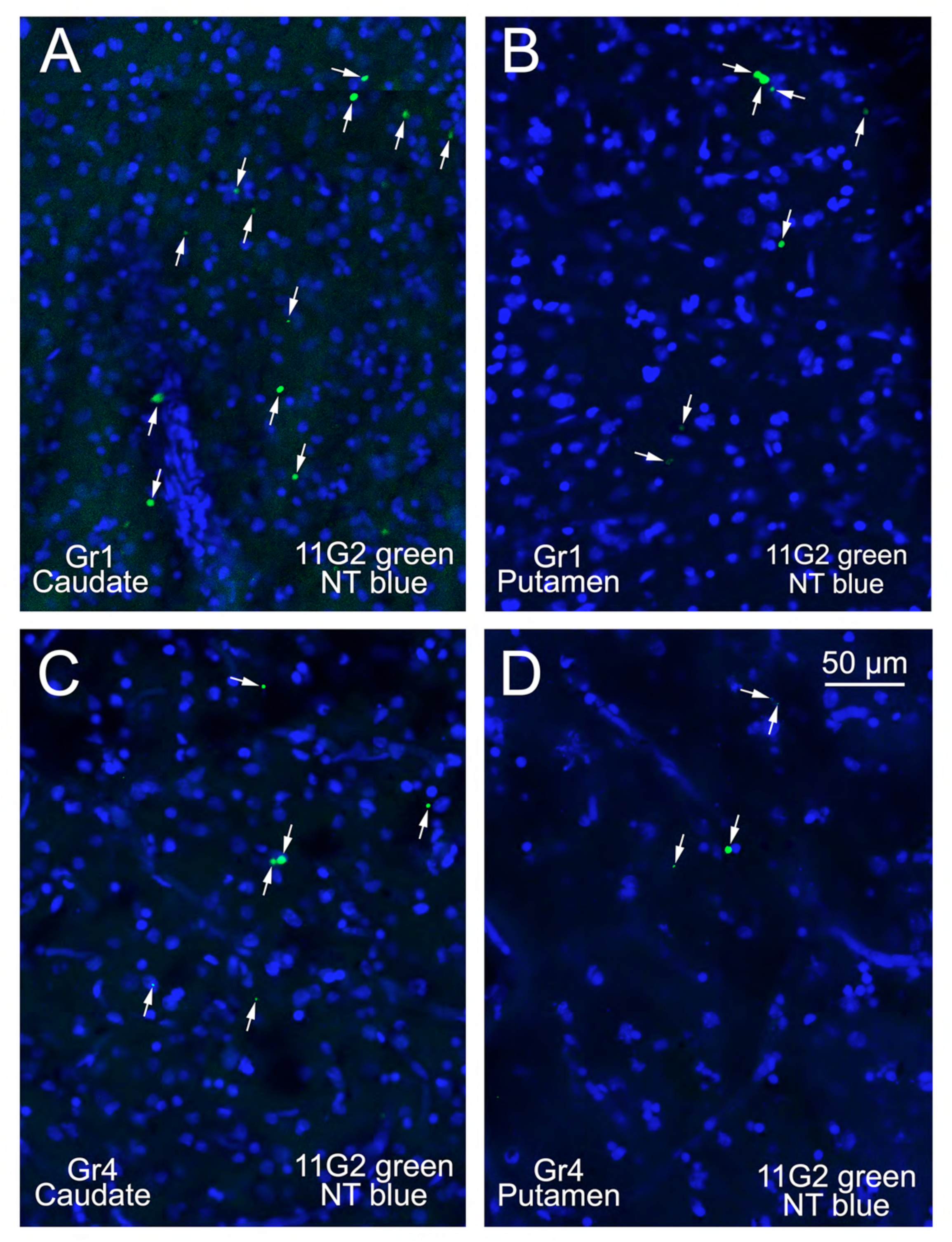
CLSM images of a single field of view through caudate and putamen, respectively, from a grade 1 HD case (A, B) and a grade 4 HD case (C, D) immunolabeled green for 11G2 with NeuroTrace (blue). 11G2+ aggregates are sparse and localized to neuropil mainly. Scale bar in D applies to all images.

**Figure 19.**
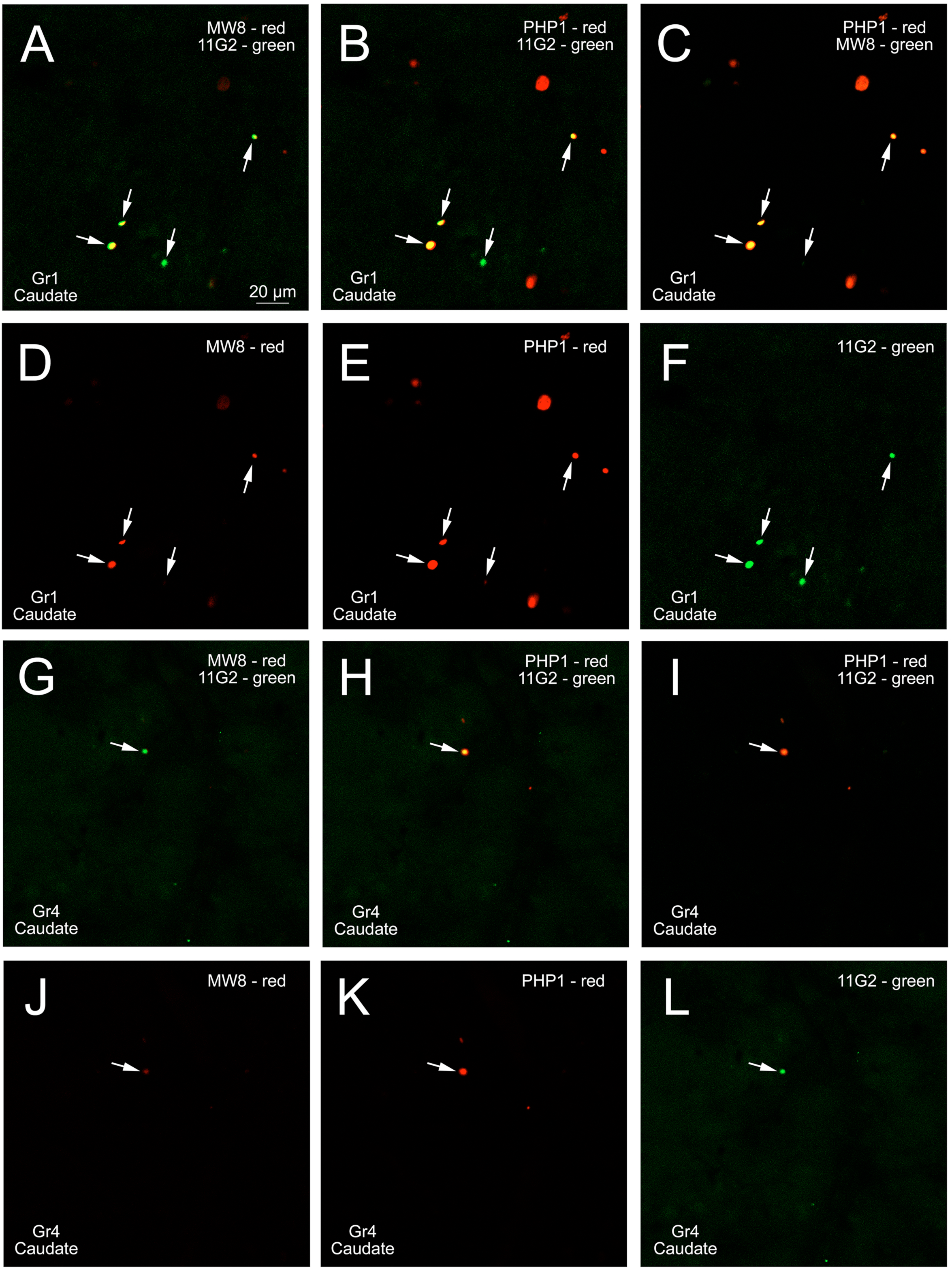
The first two rows present a single CLSM field of view through caudate from a grade 1 HD case (A-F), and the last two from a grade 4 HD case (G-L) showing the immunolabeling for 11G2, MW8, and PHP1 in each pairing (first row and third rows), and each alone (second and last rows). Arrows indicate the aggregates with 11G2 localization in the fields of view, which also tend to possess MW8 or PHP1 signal. PHP1 detects the largest aggregate area, while the more centrally located 11G2 and MW8 signals in each PHP1+ aggregate tend to be similar in areal extent. Scale bar in A applies to all images.

**Table 5.**
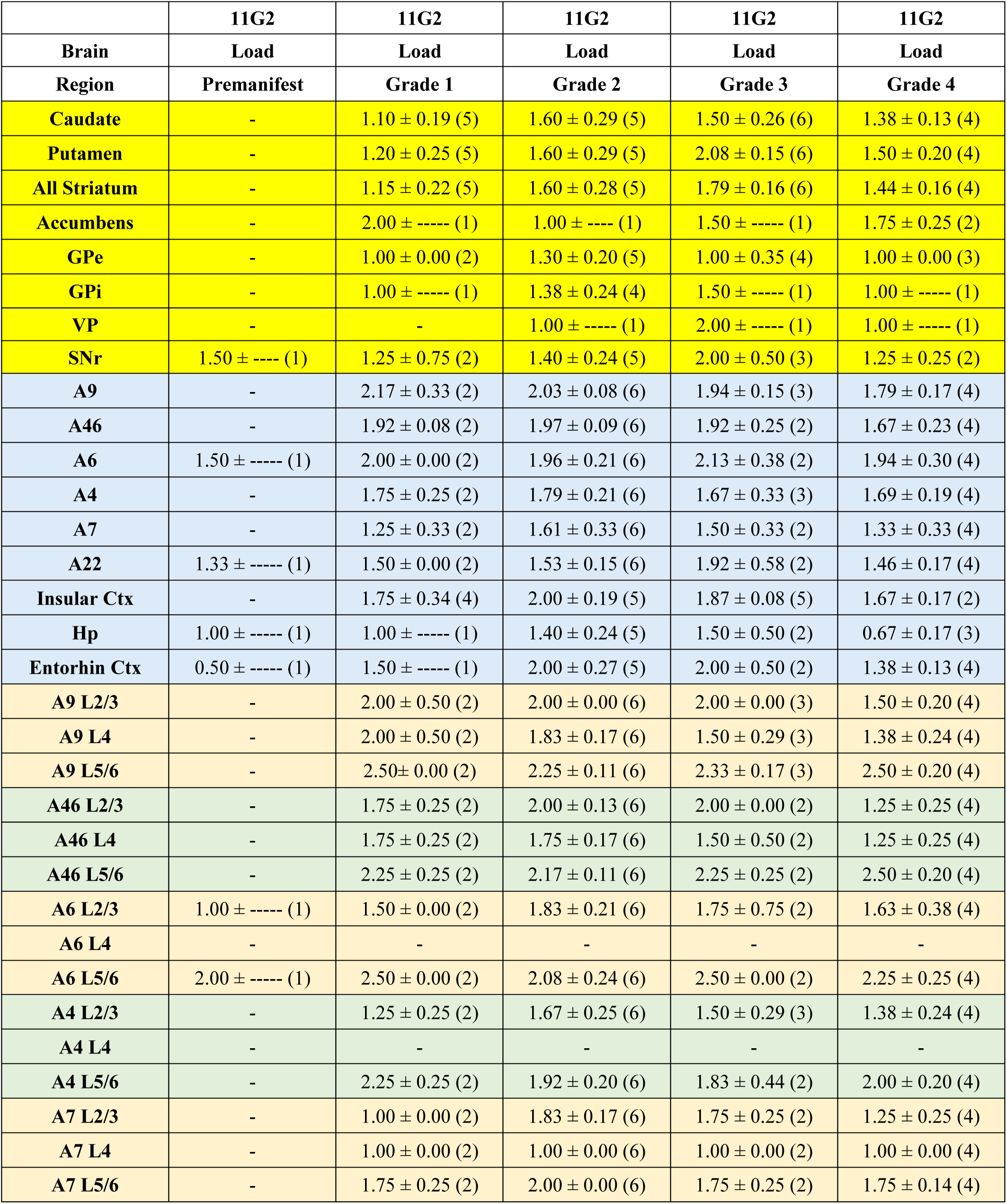
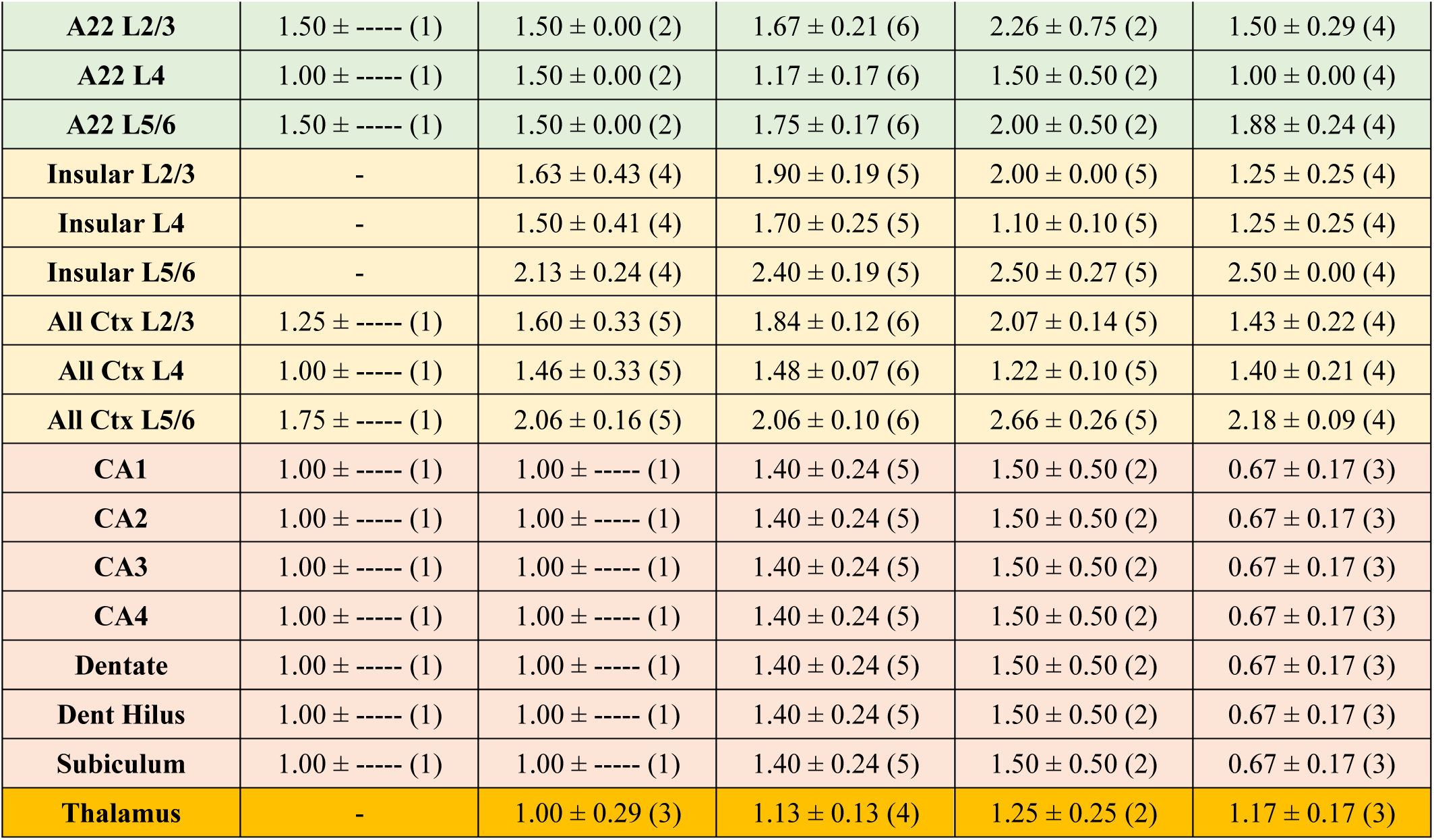
Grade-wise regional 11G2 signal in HD cases. Grade-wise 11G2 scores for human HD cases for forebrain structures, scored on a scale of 0 (none) to 4 (extremely rich).

|  | 11G2 | 11G2 | 11G2 | 11G2 | 11G2 |
| --- | --- | --- | --- | --- | --- |
| Brain | Load | Load | Load | Load | Load |
| Region | Premanifest | Grade 1 | Grade 2 | Grade 3 | Grade 4 |
| Caudate | - | 1.10 ± 0.19 (5) | 1.60 ± 0.29 (5) | 1.50 ± 0.26 (6) | 1.38 ± 0.13 (4) |
| Putamen | - | 1.20 ± 0.25 (5) | 1.60 ± 0.29 (5) | 2.08 ± 0.15 (6) | 1.50 ± 0.20 (4) |
| All Striatum | - | 1.15 ± 0.22 (5) | 1.60 ± 0.28 (5) | 1.79 ± 0.16 (6) | 1.44 ± 0.16 (4) |
| Accumbens | - | 2.00 ± ----- (1) | 1.00 ± ---- (1) | 1.50 ± ----- (1) | 1.75 ± 0.25 (2) |
| GPe | - | 1.00 ± 0.00 (2) | 1.30 ± 0.20 (5) | 1.00 ± 0.35 (4) | 1.00 ± 0.00 (3) |
| GPI | - | 1.00 ± ----- (1) | 1.38 ± 0.24 (4) | 1.50 ± ----- (1) | 1.00 ± ----- (1) |
| VP | - | - | 1.00 ± ----- (1) | 2.00 ± ----- (1) | 1.00 ± ----- (1) |
| SNr | 1.50 ± ---- (1) | 1.25 ± 0.75 (2) | 1.40 ± 0.24 (5) | 2.00 ± 0.50 (3) | 1.25 ± 0.25 (2) |
| A9 | - | 2.17 ± 0.33 (2) | 2.03 ± 0.08 (6) | 1.94 ± 0.15 (3) | 1.79 ± 0.17 (4) |
| A46 | - | 1.92 ± 0.08 (2) | 1.97 ± 0.09 (6) | 1.92 ± 0.25 (2) | 1.67 ± 0.23 (4) |
| A6 | 1.50 ± ----- (1) | 2.00 ± 0.00 (2) | 1.96 ± 0.21 (6) | 2.13 ± 0.38 (2) | 1.94 ± 0.30 (4) |
| A4 | - | 1.75 ± 0.25 (2) | 1.79 ± 0.21 (6) | 1.67 ± 0.33 (3) | 1.69 ± 0.19 (4) |
| A7 | - | 1.25 ± 0.33 (2) | 1.61 ± 0.33 (6) | 1.50 ± 0.33 (2) | 1.33 ± 0.33 (4) |
| A22 | 1.33 ± ----- (1) | 1.50 ± 0.00 (2) | 1.53 ± 0.15 (6) | 1.92 ± 0.58 (2) | 1.46 ± 0.17 (4) |
| Insular Ctx | - | 1.75 ± 0.34 (4) | 2.00 ± 0.19 (5) | 1.87 ± 0.08 (5) | 1.67 ± 0.17 (2) |
| Hp | 1.00 ± ----- (1) | 1.00 ± ----- (1) | 1.40 ± 0.24 (5) | 1.50 ± 0.50 (2) | 0.67 ± 0.17 (3) |
| Entorhin Ctx | 0.50 ± ----- (1) | 1.50 ± ----- (1) | 2.00 ± 0.27 (5) | 2.00 ± 0.50 (2) | 1.38 ± 0.13 (4) |
| A9 L2/3 | - | 2.00 ± 0.50 (2) | 2.00 ± 0.00 (6) | 2.00 ± 0.00 (3) | 1.50 ± 0.20 (4) |
| A9 L4 | - | 2.00 ± 0.50 (2) | 1.83 ± 0.17 (6) | 1.50 ± 0.29 (3) | 1.38 ± 0.24 (4) |
| A9 L5/6 | - | 2.50 ± 0.00 (2) | 2.25 ± 0.11 (6) | 2.33 ± 0.17 (3) | 2.50 ± 0.20 (4) |
| A46 L2/3 | - | 1.75 ± 0.25 (2) | 2.00 ± 0.13 (6) | 2.00 ± 0.00 (2) | 1.25 ± 0.25 (4) |
| A46 L4 | - | 1.75 ± 0.25 (2) | 1.75 ± 0.17 (6) | 1.50 ± 0.50 (2) | 1.25 ± 0.25 (4) |
| A46 L5/6 | - | 2.25 ± 0.25 (2) | 2.17 ± 0.11 (6) | 2.25 ± 0.25 (2) | 2.50 ± 0.20 (4) |
| A6 L2/3 | 1.00 ± ----- (1) | 1.50 ± 0.00 (2) | 1.83 ± 0.21 (6) | 1.75 ± 0.75 (2) | 1.63 ± 0.38 (4) |
| A6 L4 | - | - | - | - | - |
| A6 L5/6 | 2.00 ± ----- (1) | 2.50 ± 0.00 (2) | 2.08 ± 0.24 (6) | 2.50 ± 0.00 (2) | 2.25 ± 0.25 (4) |
| A4 L2/3 | - | 1.25 ± 0.25 (2) | 1.67 ± 0.25 (6) | 1.50 ± 0.29 (3) | 1.38 ± 0.24 (4) |
| A4 L4 | - | - | - | - | - |
| A4 L5/6 | - | 2.25 ± 0.25 (2) | 1.92 ± 0.20 (6) | 1.83 ± 0.44 (2) | 2.00 ± 0.20 (4) |
| A7 L2/3 | - | 1.00 ± 0.00 (2) | 1.83 ± 0.17 (6) | 1.75 ± 0.25 (2) | 1.25 ± 0.25 (4) |
| A7 L4 | - | 1.00 ± 0.00 (2) | 1.00 ± 0.00 (6) | 1.00 ± 0.00 (2) | 1.00 ± 0.00 (4) |
| A7 L5/6 | - | 1.75 ± 0.25 (2) | 2.00 ± 0.00 (6) | 1.75 ± 0.25 (2) | 1.75 ± 0.14 (4) |
| **A22 L2/3** | 1.50 ± ----- (1) | 1.50 ± 0.00 (2) | 1.67 ± 0.21 (6) | 2.26 ± 0.75 (2) | 1.50 ± 0.29 (4) |
| **A22 L4** | 1.00 ± ----- (1) | 1.50 ± 0.00 (2) | 1.17 ± 0.17 (6) | 1.50 ± 0.50 (2) | 1.00 ± 0.00 (4) |
| **A22 L5/6** | 1.50 ± ----- (1) | 1.50 ± 0.00 (2) | 1.75 ± 0.17 (6) | 2.00 ± 0.50 (2) | 1.88 ± 0.24 (4) |
| **Insular L2/3** | - | 1.63 ± 0.43 (4) | 1.90 ± 0.19 (5) | 2.00 ± 0.00 (5) | 1.25 ± 0.25 (4) |
| **Insular L4** | - | 1.50 ± 0.41 (4) | 1.70 ± 0.25 (5) | 1.10 ± 0.10 (5) | 1.25 ± 0.25 (4) |
| **Insular L5/6** | - | 2.13 ± 0.24 (4) | 2.40 ± 0.19 (5) | 2.50 ± 0.27 (5) | 2.50 ± 0.00 (4) |
| **All Ctx L2/3** | 1.25 ± ----- (1) | 1.60 ± 0.33 (5) | 1.84 ± 0.12 (6) | 2.07 ± 0.14 (5) | 1.43 ± 0.22 (4) |
| **All Ctx L4** | 1.00 ± ----- (1) | 1.46 ± 0.33 (5) | 1.48 ± 0.07 (6) | 1.22 ± 0.10 (5) | 1.40 ± 0.21 (4) |
| **All Ctx L5/6** | 1.75 ± ----- (1) | 2.06 ± 0.16 (5) | 2.06 ± 0.10 (6) | 2.66 ± 0.26 (5) | 2.18 ± 0.09 (4) |
| **CA1** | 1.00 ± ----- (1) | 1.00 ± ----- (1) | 1.40 ± 0.24 (5) | 1.50 ± 0.50 (2) | 0.67 ± 0.17 (3) |
| **CA2** | 1.00 ± ----- (1) | 1.00 ± ----- (1) | 1.40 ± 0.24 (5) | 1.50 ± 0.50 (2) | 0.67 ± 0.17 (3) |
| **CA3** | 1.00 ± ----- (1) | 1.00 ± ----- (1) | 1.40 ± 0.24 (5) | 1.50 ± 0.50 (2) | 0.67 ± 0.17 (3) |
| **CA4** | 1.00 ± ----- (1) | 1.00 ± ----- (1) | 1.40 ± 0.24 (5) | 1.50 ± 0.50 (2) | 0.67 ± 0.17 (3) |
| **Dentate** | 1.00 ± ----- (1) | 1.00 ± ----- (1) | 1.40 ± 0.24 (5) | 1.50 ± 0.50 (2) | 0.67 ± 0.17 (3) |
| **Dent Hilus** | 1.00 ± ----- (1) | 1.00 ± ----- (1) | 1.40 ± 0.24 (5) | 1.50 ± 0.50 (2) | 0.67 ± 0.17 (3) |
| **Subiculum** | 1.00 ± ----- (1) | 1.00 ± ----- (1) | 1.40 ± 0.24 (5) | 1.50 ± 0.50 (2) | 0.67 ± 0.17 (3) |
| **Thalamus** | - | 1.00 ± 0.29 (3) | 1.13 ± 0.13 (4) | 1.25 ± 0.25 (2) | 1.17 ± 0.17 (3) |

### P90 Antibody Studies in Human Cortex

We found that both the 11G2 and 1B12 antibodies detect aggregated HTT1a in HD human cortex. Blocking 11G2 with exogeneous AEEPLHRP prevented aggregate immunolabeling in HD cortex, while blocking with control peptide did not (Fig. 21; Supplementary Fig. 9). No aggregate immunolabeling with 11G2 was seen in control cortex. As for basal ganglia, we examined 11G2 localization in comparison to PHP1 and MW8 localization, in this instance in BA9 cortex in grade 1 and grade 4 cases (Figs. 22, 23). Aggregates at both grades were located mainly in the neuropil and most concentrated in layer 5. MW8 and 11G2 generally detected the same aggregates, which were a subset of those detected by PHP1. When detecting the same aggregates, MW8 and 11G2 labeled the center of the PHP1+ area. We assessed if the neuropil 11G2+ aggregates that predominate in HD cortex are localized to dendrites. To this end, we triple labeled tissue for 11G2, SMI32 and NeuroTrace (Fig. 24). We found that many large 11G2+ aggregates were contained within SMI32+ processes in cortex. Additionally, some smaller 11G2+ aggregates aligned with what appeared likely to be dendritic spines. The regional 11G2 localization scores for pallial and diencephalic structures are shown in Table 5.

**Figure 20.**
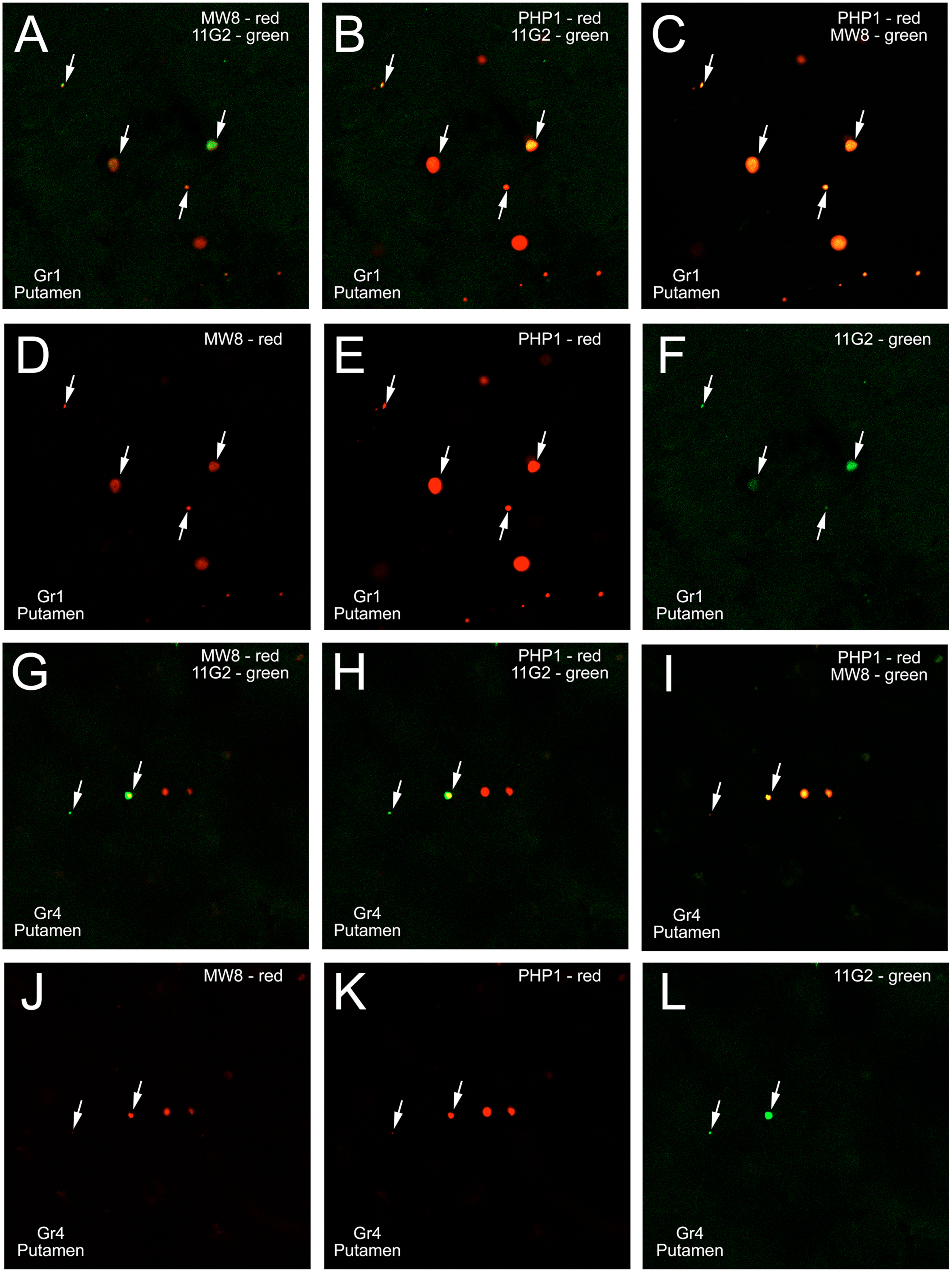
The first two rows present a single CLSM field of view through putamen from a grade 1 HD case (A-F), and the last two from a grade 4 HD case (G-L) showing the immunolabeling for 11G2, MW8, and PHP1 in each pairing (first row and third rows), and each alone (second and last rows). Arrows indicate the aggregates with 11G2 localization in the fields of view, which also tend to possess MW8 or PHP1 signal. PHP1 detects the largest aggregate area, while the more centrally located 11G2 and MW8 signals in each PHP1+ aggregate tend to be similar in areal extent. Scale bar in A applies to all images.

**Figure 21.**
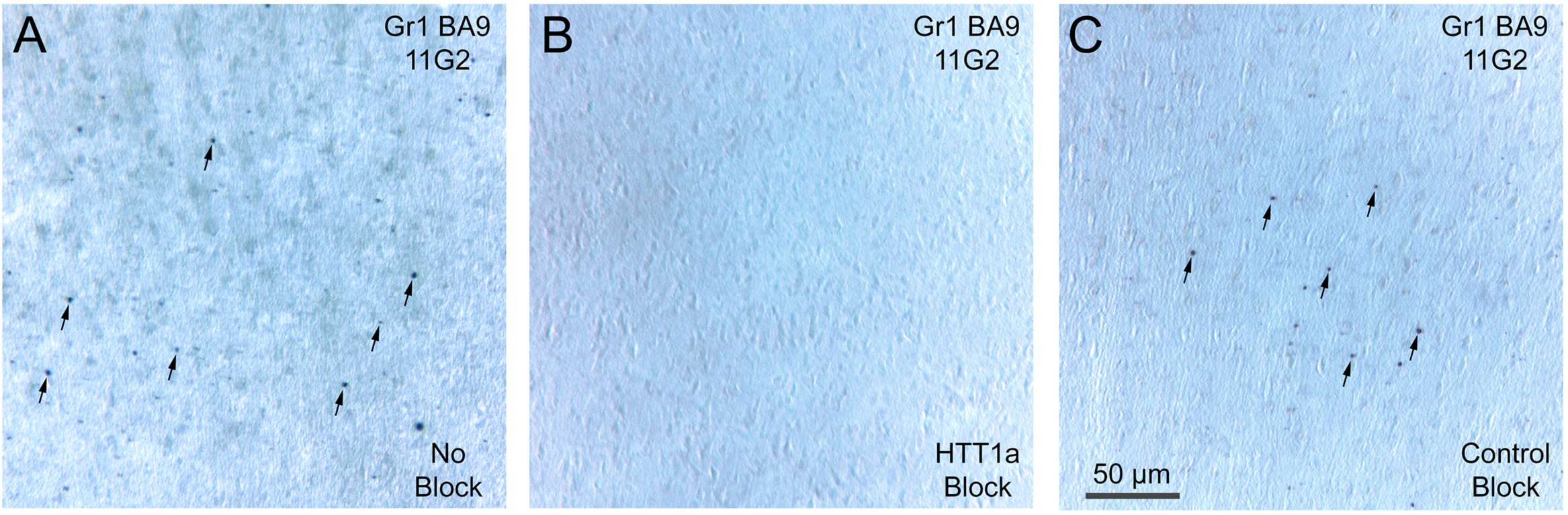
Images showing that 11G2 immunostaining in layer 5 of cortical area BA9 in a grade 1 HD case (A) is blocked when 11G2 (B) is co-incubated with HTT1a target octapeptide (AEEPLHRP), but not when 11G2 (C) co-incubated with non-target peptide (YHRLLACLQNVHKVTTC). Some of the aggregates in A and C are indicated by arrows. The scale bar applies to all images.

**Figure 22.**
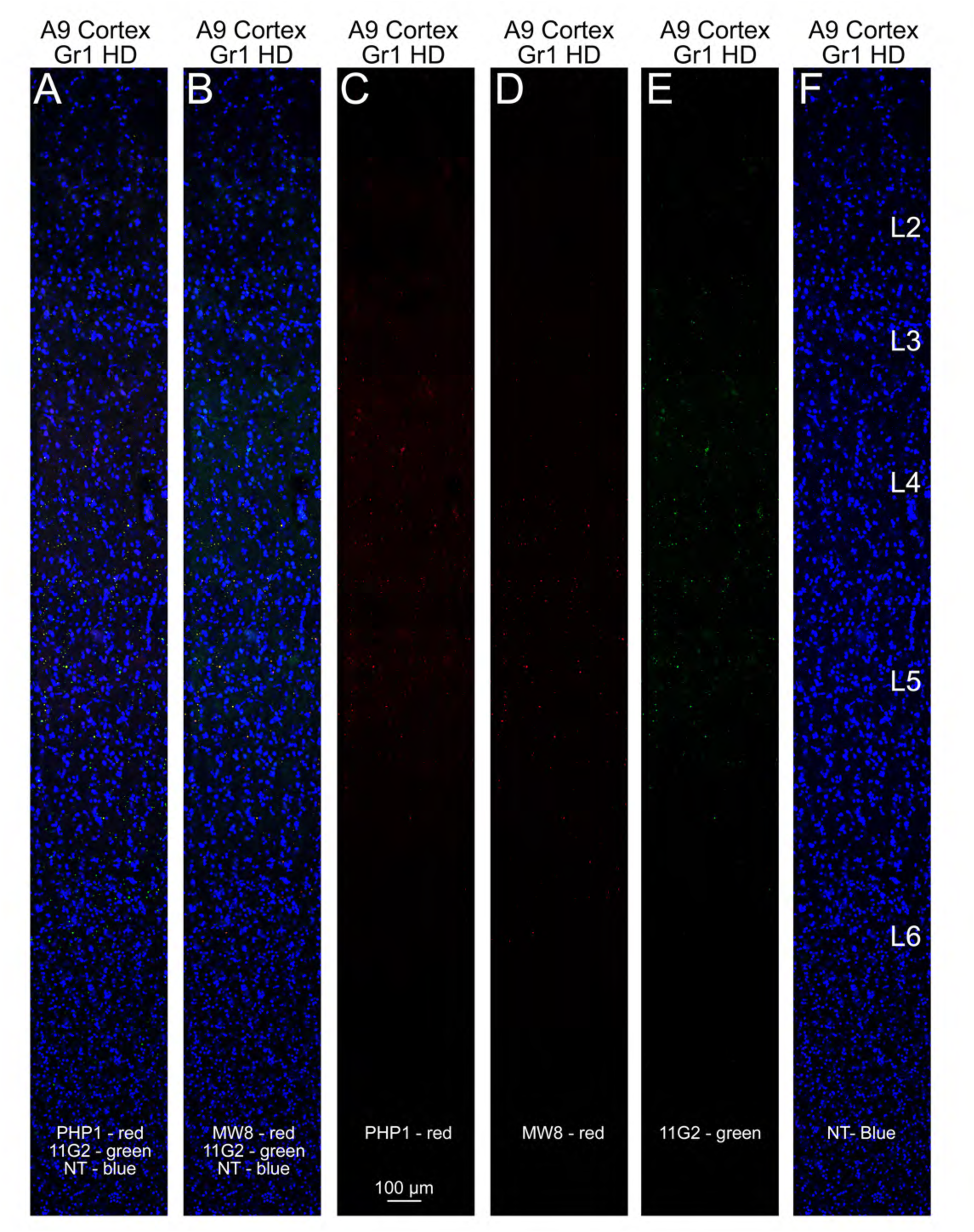
Immunofluorescence triple-label comparing localization of 11G2 (E) to that of MW8 (D) and PHP1 (C) in a single slab of BA9 cortex of a grade 1 HD case. The first panel (A) shows the merged image for PHP1 and 11G2 with a NeuroTrace counterstain, and the second (B) shows a merged image for MW8 and 11G2 with a NeuroTrace counterstain. The last panel (F) shows NeuroTrace counterstain alone. The three antibodies largely detect the same aggregates, with PHP1 detecting more aggregates and more of any given aggregate area. Scale bar in C applies to all images. Note that in this view of the full depth of BA9, the aggregates appear as tiny speckles that can be better seen by zooming in on the image in the PDF version of this paper, or by examining the higher magnification views of layers 3 and 5 in Figure 23 for this same slab.

**Figure 23.**
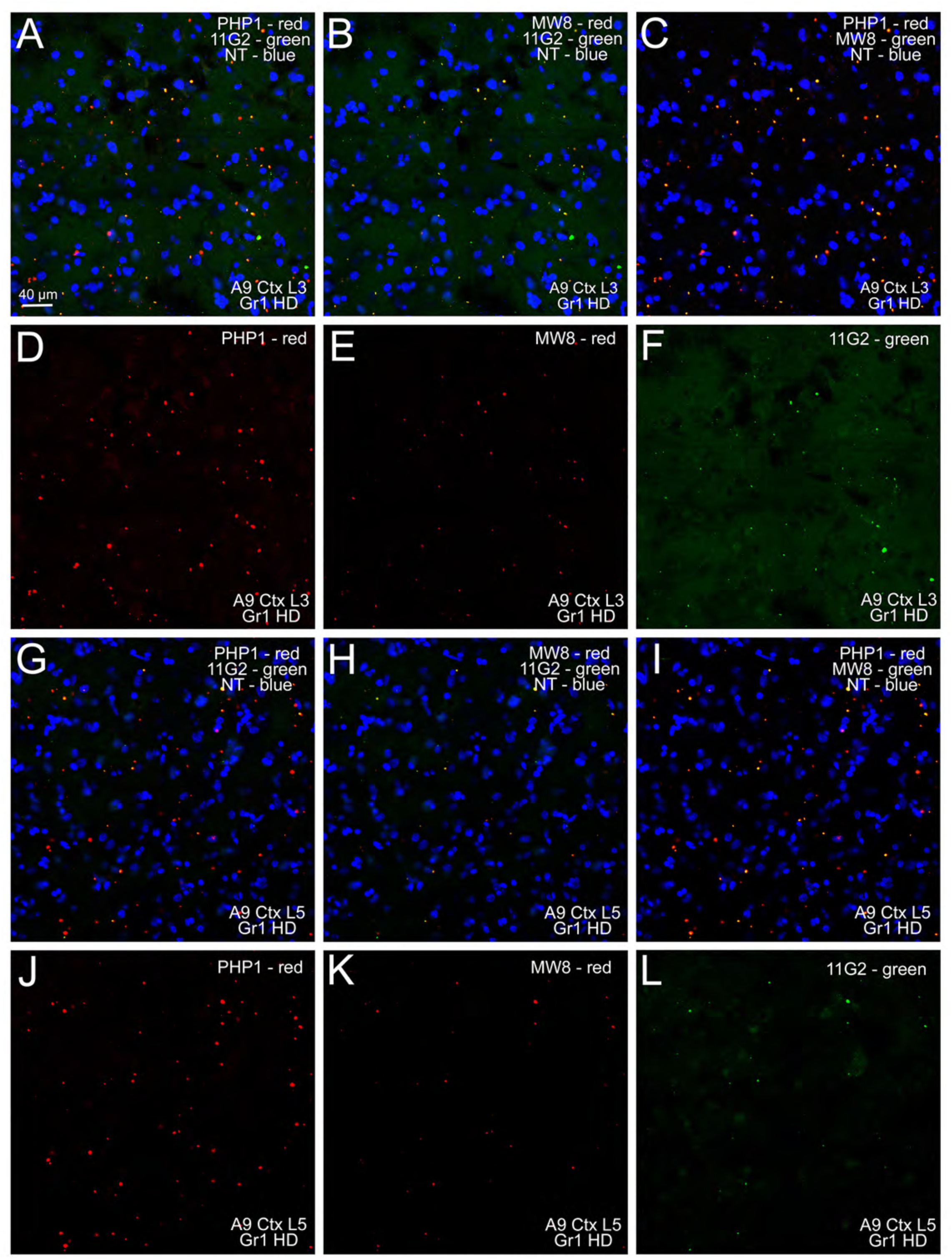
Immunofluorescence triple-label comparing localization of 11G2, MW8, and PHP1 in single fields of view of layer 3 (rows 1 and 2) and layer 5 (rows 3 and 4) of BA9 cortex from a grade 1 HD case, with a NeuroTrace counterstain. The first and third rows show each pairing for 11G2, MW8, and PHP1, and the second and fourth rows show each separately. The three antibodies largely detect the same aggregates, with PHP1 detecting more aggregates and more of any given aggregate. Scale bar in A applies to all images. Note that the aggregates appear as speckles that can be better seen by zooming in on the image in the PDF version of this paper.

**Figure 24.**
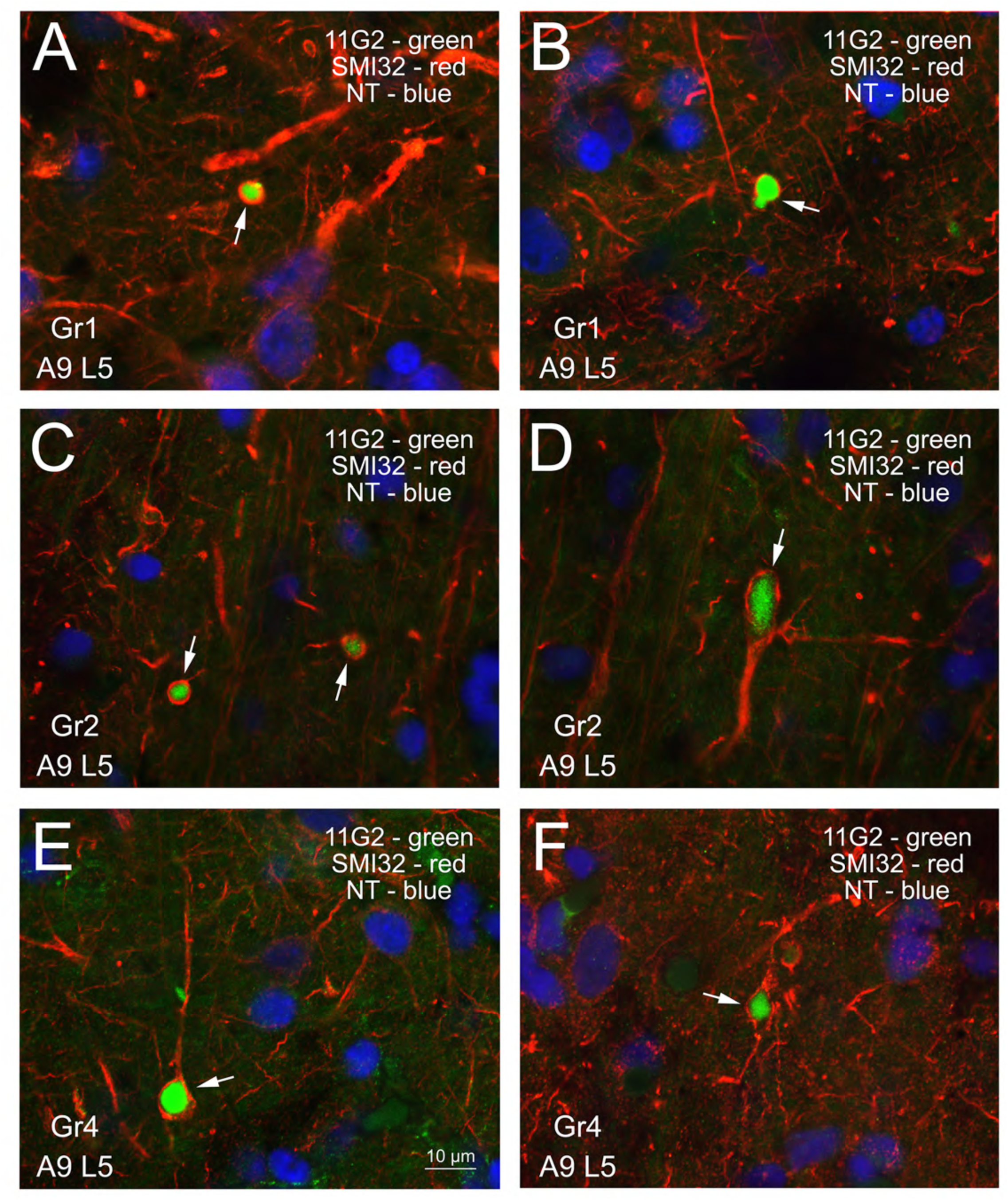
Panels showing merged CLSM images of layer 5 of BA9 cortex from a grade 1 HD case (A, B), a grade 2 case (C, D) and a grade 4 case (E, F), with cortical neurons labeled blue with NeuroTrace, 11G2+ aggregates shown in green, and the pyramidal neuron dendrite marker SMI32 shown in red. Many large 11G2+ aggregates are encapsulated by SMI32+ processes. Scale bar in E applies to all images.

## Discussion

In the present study, we show that the mutant HTT1a does accumulate and aggregate in Q175 mouse brain, R6/2 mouse brain, and human HD brain, as expected from the work of others (Sathasivam et al., 2013; Neueder et al., 2017; Smith et al., 2023). Moreover, the magnitude of its differential regional accumulation in forebrain is in many regards consistent with the differential vulnerability of those regions in HD. Nonetheless, accumulation for some regions and some neuron types was inconsistent with a determinative role of mutant HTT1a abundance in the vulnerability differences for those regions and those neuron types. Moreover, the differences between mutant mice and HD humans in the role of somatic expansion of the CAG repeat in the disease process need to be taken into account in comparing the current Q175 mouse and human HD results, as discussed in more detail below.

### HTT1a Accumulation – HD mice and humans

In Q175 mice and R6/2 mice but not N171-82Q mice, the P90 antibodies yielded nuclear and aggregate immunolabeling in many neurons throughout brain, showing that neurons in Q175 mice and R6/2 mice but not N171-82Q mice produce mutant HTT1a. Consistent with this, Sapp et al. (2025) detected HTT1a in Q175 and R6/2 mouse brain by western blot. This supports the selectivity of the P90 antibodies for mutant HTT1a, as the transgene in R6/2 mice specifically produces mutant HTT1a and the chimeric mutant huntingtin gene in Q175 mice produces HTT1a due to the same splicing anomaly that occurs in human HD (Sathasivam et al., 2013; Neueder et al., 2017). By contrast, the transgene in N171-82Q mice spans exons 1 and 2, and is thus not subject to the same splicing anomaly that leads to mutant HTT1a build-up in Q175 mice and human HD (Schilling et al., 1999; Smith et al., 2023). Moreover, nuclear and aggregated immunolabeling in Q175 mice with 11G2 was blocked by pre-incubating it with exogeneous HTT1a C-terminal octapeptide, and greatly attenuated in R6/2 mice.

Although the P90 antibodies do not distinguish between mutant and WT HTT1a, as their epitope is the eight C-terminal residues of HTT1a (AEEPLHRP) shared in mouse and human, they potently detect mutant HTT1a because of its propensity to accumulate as oligomers and aggregates. Nonetheless, we did see P90 immunolabeling in WT mice, but it was unlike the nuclear and aggregate immunolabeling seen in Q175 and R6/2 mice. Specifically, in WT mice we saw relatively light cytoplasmic signal in some neuronal populations that also immunolabel with antibodies against full-length huntingtin, notably in the perikarya of layer 5 cortical neurons, in the perikarya of scattered large striatal neurons, and in presumptive SPN terminals in GPe, GPi and SNr (Fusco et al., 1999; Dragatsis et al., 2018). Similar cortical and SPN target immunolabeling was seen in 2-month old Q175 mice, and in N171-82Q mice. This immunolabeling was eliminated by pre-incubating 11G2 with exogeneous AEEPLHRP, and reduced in conditional deletion mice with 90% reduction in WT huntingtin. Our P90 immunolabeling results in WT mice have either of three explanations: 1) WT mice produce some amount of HTT1a from their endogeneous WT huntingtin alleles; 2) WT mice produce some form of HTT longer than HTT1a that is detected by the P90 antibodies despite their apparent specificity for HTT1a; and 3) WT mice produce a protein other than HTT1a that contains antigenic determinants similar to AEEPLHRP, is detected by the P90 antibodies, and has the same distribution as HTT. With regard to the first possibility, some prior studies show data suggesting some low level of production of *HTT1a* message (Hoschek et al., 2024) and HTT1a protein (Neueder et al., 2017; Landles et al., 2024; Sogorb-Gonzalez et al., 2024), and Hoschek et al. (2024) appear to suggest that a low level of HTT1A production from the WT huntingtin alleles is to be anticipated. Although possibilities 2 and 3 cannot be ruled out, they seem less likely given the demonstrated specificity of the P90 antibodies, which are monoclonal antibodies that all evidence shows to be specific for the last 8 amino acids at the C-terminus of HTT1a.

Nonetheless, it should be noted that Sapp et al., were unable to detect HTT1a by western blot in WT mouse or control human brain with either 11G2 or 1B12. In any case, the possibility suggested by our data of a low level of HTT1a production from the WT huntingtin alleles that is detectible by the P90 antibodies in those neuron types and projections systems most enriched in HTT would benefit from further study.

In R/2 mice, intense nuclear P90 signal was observed in all or nearly all neurons throughout brain by 9 weeks of age, with the signal consisting of intense diffuse labeling of the entire nucleus, and a nuclear aggregate evident by CLSM but obscured in DAB-labeled material by pervasive nuclear labeling. Consistent with its slower pace of exon 1 protein production, P90 signal was largely localized to SPNs at this same age in Q175 heterozygotes, with it manifesting as light diffuse nuclear labeling. A similar result was reported by Smith et al., (2023) using the S830 antibody against mutant HTT1a. The early preferential accumulation of HTT1a in the nuclei of SPNs indicates that even when all neurons in the brain possess a huntingtin gene with the same expanded pathogenic repeat, SPNs are still nonetheless prone to accumulating more of this protein – consistent with their greater vulnerability in HD. The early occurrence at 2 months of diffuse nuclear localization of mutant HTT1a in SPNs is associated with a slight reduction in preproenkephalin expression by iSPNs (Deng et al., 2021). With age progression, mutant HTT1a accumulation gradually becomes yet more prominent in striatum, and more widespread in the brain, with nuclear aggregates and/or neuropil aggregates present in many brain regions, and neurochemical and behavioral deficits more evident (Deng et al., 2021; Menalled et al., 2012). Even by 18 months, however it is not nearly as intense and widespread in Q175 heterozygotes as it is in 9-week old R6/2 mice. The transition from diffuse labeling to increasing aggregation aligns with the change in solubility of HTT1a seen by SDS-PAGE and western blot assays in cortex lysates from HD knock-in mice (Sapp et al., 2025). With increasing age, the migration of soluble HTT1a slowed and diminished in intensity and was replaced by a high molecular mass smear. Even small differences in the number of CAG repeats between Q111 mice altered HTT1a migration and solubility.

We probed the composition of the aggregates, comparing labeling with 11G2 with labeling obtained using the PHP1 and MW8 antibodies in striatum and cortex of 6-18 month old Q175 mice. At six months, the MW8+ signal occupied only the center of 11G2+ aggregates in Q175 striatum. The aggregate area occupied by PHP1 signal at six months was even less. By 12 months, both MW8 signal and PHP1 signal occupied more of the 11G2+ aggregate area, but still not its entirety. Thus, our results show that the PHP1 and MW8 target epitopes are only antibody-accessible in the core of the 11G2+ aggregates in these mice, and that the epitope targeted by MW8 is very similar but not the same as that for 11G2, as also observed in western blot assays by Sapp et al. (2025). Additionally, at both ages, diffuse nuclear signal was more evident with 11G2 than PHP1 and MW8, indicating that the C-terminus of HTT1a is more accessible to the P90 antibodies in its diffuse, presumably monomeric or oligomeric, form in Q175 mice, than the polyproline region of HTT1a detected by PHP1. Our results thus show that the presumably toxic huntingtin mutant HTT1a is more pervasive in Q175 mouse than revealed by antibodies such as PHP1 and MW8. Similarly, mutant HTT1a is more pervasive in R6/2 mice than suggested by antibodies such as EM48 or those directed at ubiquitin (DiFiglia et al., 1997; Meade et al., 2002; Bayram-Weston et al., 2016).

In our human HD cases as well, the P90 antibodies yielded clear immunolabeling of aggregates, albeit with far less regional density than in Q175 mouse. As the mutant *HTT1a* transcript and the mutant HTT1a protein have been shown to be produced in human HD brain (Sathasivam et al., 2013; Neueder et al., 2017), our human findings with the P90 antibodies show that mutant HTT1a accumulates and forms aggregates in HD brain. Although P90+ nuclear and neuropil aggregates were both common in Q175, especially older ones, and R6/2 mice, the P90 antibodies in human HD cases mainly labeled neuropil aggregates in striatum and cortex.

Similarly, prior studies using other antibodies have also found mutant protein aggregates in only a small percent of human striatal neurons of HD patients (DiFiglia et al., 1997; Maat-Schieman et al., 1999; Gutekunst et al., 1999; Kuemmerle et al., 1999), despite their prominence in striatal neurons in mouse models of HD (Meade et al., 2002; Brooks and Dunnett, 2015). Part of the explanation for this difference for striatum at least may stem from the asynchronous nature of the CAG repeat somatic expansion required for mutant protein production and accumulation in the human disease (Ma tlik et al., 2024; Handsaker et al., 2024). It seems likely that, at any moment in time, only a small number of SPNs will have progressed to a repeat expansion sufficient to yield mutant HTT1a accumulation, and that neurons die relatively rapidly after reaching this point. This window of time is uncertain, but Handsaker et al. (2025) have suggested it is on the order of months rather than years. Although this consideration may explain why aggregates per se are scarce in human striatum, it does not explain why neuropil aggregates predominate. As SPN vulnerability to mutant protein in mouse models seems to prominently involve a pathogenic effect in the nucleus (Benn et al., 2005; Dragatsis et al., 2009; Gu et al., 2015), the predominantly non-nuclear localization in human HD striatum raises questions about the pathogenic mechanism of mutant huntingtin in SPNs. It is possible, however, that the suboptimal fixation of human tissue hinders mutant HTT1a localization. Aggregates immunolabeled for 11G2 were somewhat more common in cerebral cortex in HD brain than in striatum, but in cortex as well they were largely restricted to the neuropil, in many cases to dendrites. Other investigators have reported them in axons as well (Sapp et al., 1999). This raises questions about the mechanism by which mutant HTT1a accumulation injures cortical neurons. Nuclear accumulation clearly can disrupt transcriptional activity, but dendritic or axonal localization could also adversely and deleteriously impair cell function and viability (Gutekunst et al., 1999; Zuccato and Cattaneo, 2014; Vitet et al., 2020).

### Differential Regional and Cellular Vulnerability – HD mice and humans

For Q175 mice and human HD brains, the overall regional distribution of 11G2+ signal in forebrain, as well as specifically in basal ganglia structures, appeared consistent with the HD vulnerability of those regions. For example, 11G2 content was higher in more vulnerable basal ganglia structures than less vulnerable pallial structures in Q175 mouse, and greater in the more vulnerable striatal than the less vulnerable pallidal structures in Q175 mouse and human HD. In relating mouse to human, however, it needs to be noted that a key factor in mutant HTT1a accumulation in humans is the regional differences in somatic expansion of the CAG repeat (Mätlik et al., 2024; Pressl et al., 2024; Handsaker et al., 2025). This phenomenon seems, in particular, to explain the greater vulnerability of SPNs than other cell types in HD (Mätlik et al., 2024; Pressl et al., 2024; Handsaker et al., 2025). The greater vulnerability of iSPNs relative to dSPNs, despite the inherently greater propensity for mutant protein accumulation we saw in dSPNs relative to iSPNs in Q175 mice, is perhaps explainable by a slightly greater propensity of iSPNs for CAG expansion (Handsaker et al., 2025). Without expansion to the 150 repeats suggested by Handsaker et al. (2025) as needed for pathogenesis, cell types that produce copious HTT1a in Q175 mouse, might make much less in human HD brain. This may explain why structures such as CA1 and pyriform cortex, which show an extreme accumulation of HTT1a in all neuronal nuclei in Q175 mice, have not been found to show notable pathology in HD (Spargo et al., 1993; Reiner et al., 2011). Some regions such as hippocampus may also have intrinsically greater resistance to the effects of mutant HTT1a accumulation, because even in Q175 mice, in which hippocampal HTT1a accumulation rivals that in striatum, pathology seems demonstrably less in hippocampus than striatum (Solem et al. 2025; Häggkvist et al., 2017.).

On the other hand, some neuron types, such as striatal cholinergic interneurons, are known to show somatic expansion in human HD (Mätlik et al., 2024; Handsaker et al., 2025) and produce large amounts of huntingtin (Fusco et al., 1999), but nonetheless were found by us to show little mutant HTT1a aggregate formation. The meager accumulation of mutant HTT1a in cholinergic interneurons in Q175 mice, while consistent with their resistance in HD (Massouh et al., 2008), is not predictable from their expected large production of mutant HTT1a. Possibly, cholinergic interneurons may have more active degradative mechanisms (Waelter et al., 2001; Zhao et al., 2024). On the other hand, PARV+ striatal interneurons show little somatic expansion in human HD (Mätlik et al., 2024; Handsaker et al., 2025), yet their HD vulnerability is comparable to that of iSPNs (Reiner et al., 2013). Our results for Q175 mice show that with a CAG repeat length of 190, PARV+ interneurons accumulate more mutant HTT1a than do cholinergic interneurons. With their limited somatic expansion in human HD, however, PARV+ interneurons would not be expected to accumulate pathogenic levels of HTT1a, raising questions then about the basis of their loss in human HD. These results suggest that selective neuronal vulnerability in HD is not tightly correlated with HTT1a aggregate load and distribution.

In line with considerations above, the vulnerability of layer 5A cortical neurons in HD (Reiner et al., 2011) relative to their low levels of HTT1a accumulation is of interest as well. Neurons of cortical layer 5A project only within telencephalon, inclusive of other cortical areas and striatum, and are thus termed intratelencephalically projecting cortical neurons (IT-type) (Reiner et al., 2003). The neurons of layer 5B also project to striatum, but to brainstem and spinal cord as well via the pyramidal tract, and are thus called pyramidal-tract type (PT-type) (Reiner et al., 2003). The IT neuron type or a subset of it has been found to show somatic expansion in human HD, while the PT-type or a subset of it does not (Pressl et al., 2024).

Although neuropathological studies have not reported differential pathology of layer 5A versus 5B neurons in HD (Reiner et al., 2011), Pressl et al. (2024) report greater loss of 5A than 5B neurons in HD based on single-nuclei analysis, and similarly Deng et al. (2014) reported greater loss of IT-type corticostriatal terminals (i.e., 5A) than PT-type corticostriatal (i.e., 5B) terminals in Q140 mouse. Surprisingly, we observed lesser HTT1a accumulation in layer 5A neurons than 5B neurons in Q175 mice. Layer 5A and 5B neurons are, however, not just singular types and each possesses a diversity of subtypes (Sorensen et al., 2015; Bakken et al., 2021; Hausmann et al., 2022). It would be useful to relate that diversity to somatic expansion in human HD, their vulnerability in HD, and mutant HTT1a accumulation in mouse HD models and in human HD.

### Aggregate Composition

The relative localization of the epitopes detected by the 11G2, MW8 and PHP1 antibodies differed between Q175 mouse and human HD. In mouse, MW8 and PHP1 signal occupied the core of aggregates defined by 11G2 signal, and showed less diffuse nuclear signal. Since all three antibodies target HTT1a, the results suggest that in the oligomerization and aggregation process, the 11G2 epitope remains accessible throughout, while the MW8 and PHP1 epitopes are cryptic or largely so, during the oligomer stage and parts of the aggregation stage in mouse. By contrast, in human HD, MW8 and 11G2 generally detected the same aggregates, which were a subset of those detected by PHP1. When detecting the same aggregates as PHP1, the MW8 and 11G2 antibodies labeled the center of the PHP1+ area. These results seem to suggest that the aggregated mutant HTT1a targeted by MW8 and 11G2 might be the seed around which the aggregate enlarges in HD brain, with the aggregate periphery potentially including larger forms of the mutant protein, as well as WT huntingtin protein. The interpretation of these results needs to take into consideration the suboptimal fixation of human tissue compared to Q175 tissues, and the antigen retrieval used in human but not mouse – both of which affect antigen accessibility (Meade et al., 2002; Bayram-Weston and Weston, 2016; Smith et al., 2023). Nonetheless, we know from our own studies that antigen retrieval does not notably affect PHP1 and MW8 localization in Q175 mouse, and so it seems unlikely that antigen retrieval alone accounts for the human – mouse difference we see in the relative localization of 11G2, MW8 and PHP1 signal. Whether fixation conditions account for the mouse – human difference in aggregate epitope localization, and which in any case provides the truer insight into aggregate composition and formation in human disease, is uncertain.

## Acknowledgements

We thank Dr. Jeff Carroll of the University of Washington, Dr. Deanna Marchionini of the CHDI, and Dr. Wenzhen Duan of Johns Hopkins University for providing tissues used in this study. We thank the CHDI for providing the P90 antibodies, and Drs. Liliana Menalled, Liz Doherty, and Jonathan Bard for details about antibody production and comments on this manuscript. This study was supported by the CHDIF (AR) and the Dake Family Fund (MD).

## Author contributions

YD, MJ, RC, and ES conducted the studies. YD, AR, ES and MD conducted analysis. AR, YD, MJ, ES and MD prepared the manuscript.

## Ethical considerations

All animal use was carried out in accordance with the National Institutes of Health Guide for Care and Use of Laboratory Animals and Society for Neuroscience Guidelines, and with the approval of the University of Tennessee Health Science Center Animal Care and Use Committee (approval # 21-0308) on November 8, 2021. All human studies were conducted on de-identified postmortem tissue, and thus the University of Tennessee Health Science Center Institutional Review Board waived the requirement for approval (16-04430-NHSR), and no informed consent was required.

## Consent to participate

Not applicable.

## Consent for publication

Not applicable.

## Data availability

Data collected as part of this study will be made available upon reasonable request.

## Supplementary Figures

**Supplementary Figure 1.** Images showing 11G2 immunolabeling in the right CEA of a 4-month old N171-82Q mouse without antigen retrieval (no AR) (A) and with antigen retrieval (AR) (B), in the right GPe with antigen retrieval (C), and in layer 5 of cortex with antigen retrieval (D). The scale bar applies to all images.

**Supplementary Figure 2.** Images showing that 11G2 immunostaining in 2-month old WT and Q175 mouse cortex is blocked by HTT1a target octapeptide (AEEPLHRP), but not with non-target control peptide (YHRLLACLQNVHKVTTC). The first row (A-C) shows results for Q175 mouse, and the second (D-F) shows a WT littermate, with each row showing the same cortical region immunolabeled with 11G2 (A, D), 11G2 plus target octapeptide (B, E), and 11G2 plus non-target octapeptide (C, F). Layer 4 (L4) neurons are labeled in the first and last images of the first row, and layer 5 (L5) neurons in the first and last images of the second row. Insets to the lower left of each of the high magnification images show the section from which the high magnification image came, with asterisks marking the location of the field of view in the high-power image. The scale bar applies to all six main images.

**Supplementary Figure 3.** Images showing that 11G2 immunostaining in 12-month old Q175 mouse striatum and GPe are blocked by HTT1a target octapeptide (AEEPLHRP), but not with non-target control peptide (YHRLLACLQNVHKVTTC). The first row (A-C) shows results for left striatum, and the second for right GPe (D-F), with each row showing the same region immunolabeled with 11G2 (A, D), 11G2 plus target octapeptide (B, E), and 11G2 plus non-target octapeptide (C, F). Insets to the lower left of each of the high magnification images show the section from which the high magnification image came, with asterisks marking the location of the field of view in the high-power image. The scale bar applies to all six main images.

**Supplementary Figure 4.** Images showing that 11G2 immunostaining in 12-month old WT and Q175 mouse cortex is blocked by HTT1a target octapeptide (AEEPLHRP), but not with non-target control peptide (YHRLLACLQNVHKVTTC). The first row (A-C) shows results for Q175 mouse, and the second (D-F) shows a WT littermate, with each row showing the same cortical region immunolabeled with 11G2 (A, D), 11G2 plus target octapeptide (B, E), and 11G2 plus non-target octapeptide (C, F). Layer 5 (L5) and layer 6 (L6) neurons are labeled in the first and last images of the first row, and layer 5 (L5) neurons in the first and last images of the second row. Insets to the lower left of each of the high magnification images show the section from which the high magnification image came, with asterisks marking the location of the field of view in the high-power image. The scale bar applies to all six main images.

**Supplementary Figure 5.** Images showing that 11G2 immunostaining in 12-month old Q175 mouse left hippocampus CA1 and right hypothalamus are blocked by HTT1a target octapeptide (AEEPLHRP), but not with non-target control peptide (YHRLLACLQNVHKVTTC). The first row (A-C) shows results for hippocampus, and the second (D-F) for hypothalamus, with each row showing the same region immunolabeled with 11G2 (A, D), 11G2 plus target octapeptide (B, E), and 11G2 plus non-target octapeptide (C, F). Insets to the lower left of each of the high magnification images show the section from which the high magnification image came, with asterisks marking the location of the field of view in the high-power image. The scale bar applies to all six main images.

**Supplementary Figure 6.** Images showing that 11G2 immunostaining in 71-day old R6/2 mouse striatum is greatly attenuated by HTT1a target octapeptide (AEEPLHRP). The first image (A) shows the 11G2 immunolabeling without block and the second (B) with 11G2 plus target. Insets to the lower left of each of the high magnification images show the section from which the high magnification image came, with asterisks marking the location of the field of view in the high-power image. The scale bar applies to both main images.

**Supplementary Figure 7.** Images showing that 11G2 immunostaining in 71-day old R6/2 and WT mouse cortex is greatly attenuated by HTT1a target octapeptide (AEEPLHRP). The first image (A) shows the 11G2 immunolabeling without block and the second (B) with 11G2 plus target. Insets to the lower left of each of the high magnification images show the section from which the high magnification image came, with asterisks marking the location of the field of view in the high-power image. The scale bar applies to both main images.

**Supplementary Figure 8.** Images showing that 11G2 immunostaining of aggregates in Grade 1 HD striatum is blocked by HTT1a target octapeptide (AEEPLHRP), but not with non-target control peptide (YHRLLACLQNVHKVTTC). Image A shows results for striatum with 11G2, image B for 11G2 plus target octapeptide, and image C for 11G2 plus non-target octapeptide. Arrows indicate some of the 11G2+ aggregates. The scale bar applies to all three images.

**Supplementary Figure 9.** Images showing that 11G2 immunostaining of aggregates in Grade 1 HD BA9 cortex is blocked by HTT1a target octapeptide (AEEPLHRP), but not with non-target control peptide (YHRLLACLQNVHKVTTC). Image A shows results for BA9 with 11G2, image B for 11G2 plus target octapeptide, and image C for 11G2 plus non-target octapeptide. Arrows indicate some of the 11G2+ aggregates. The scale bar applies to all three images.

## Supplementary Table

**Supplementary Table 1.**
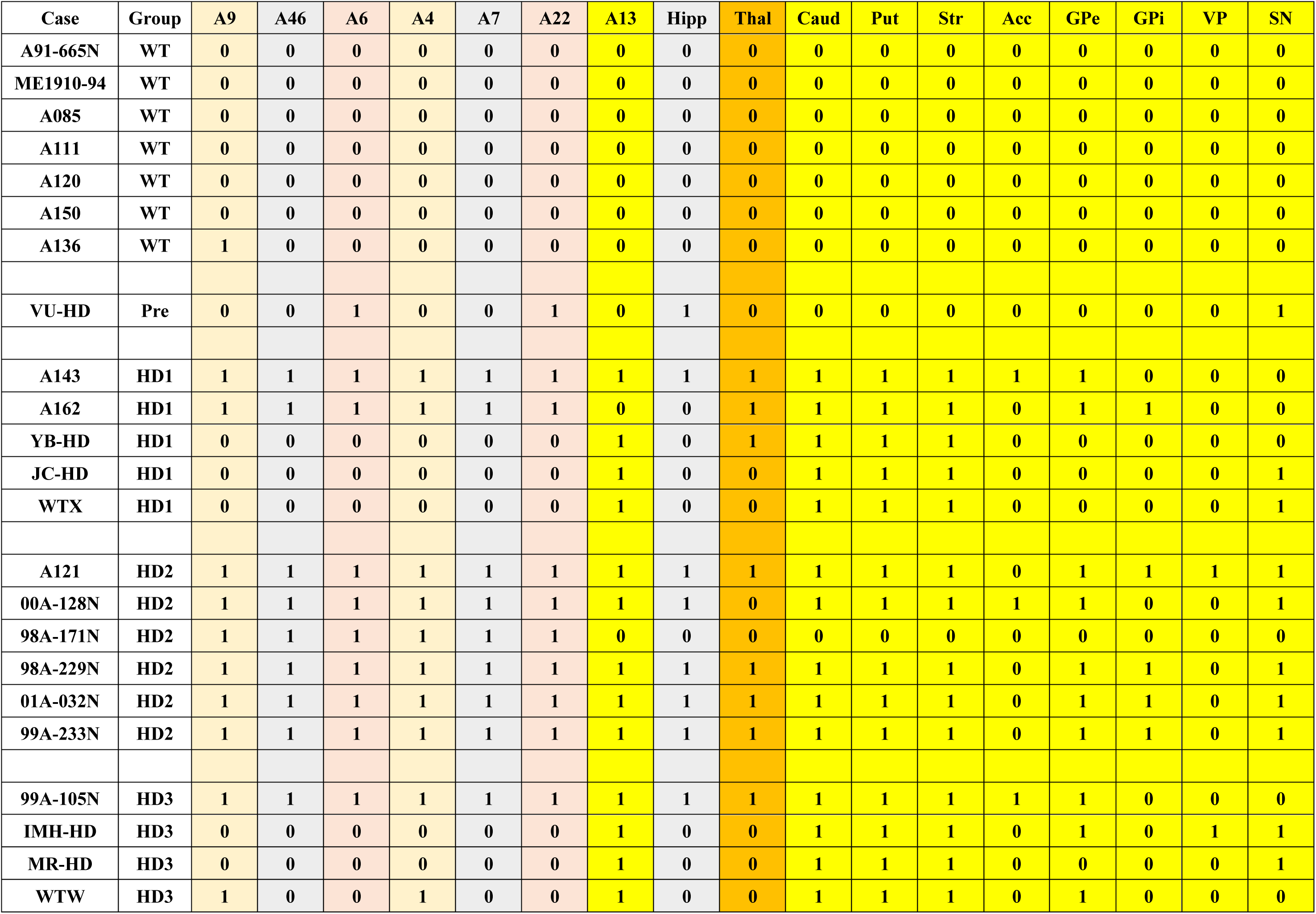
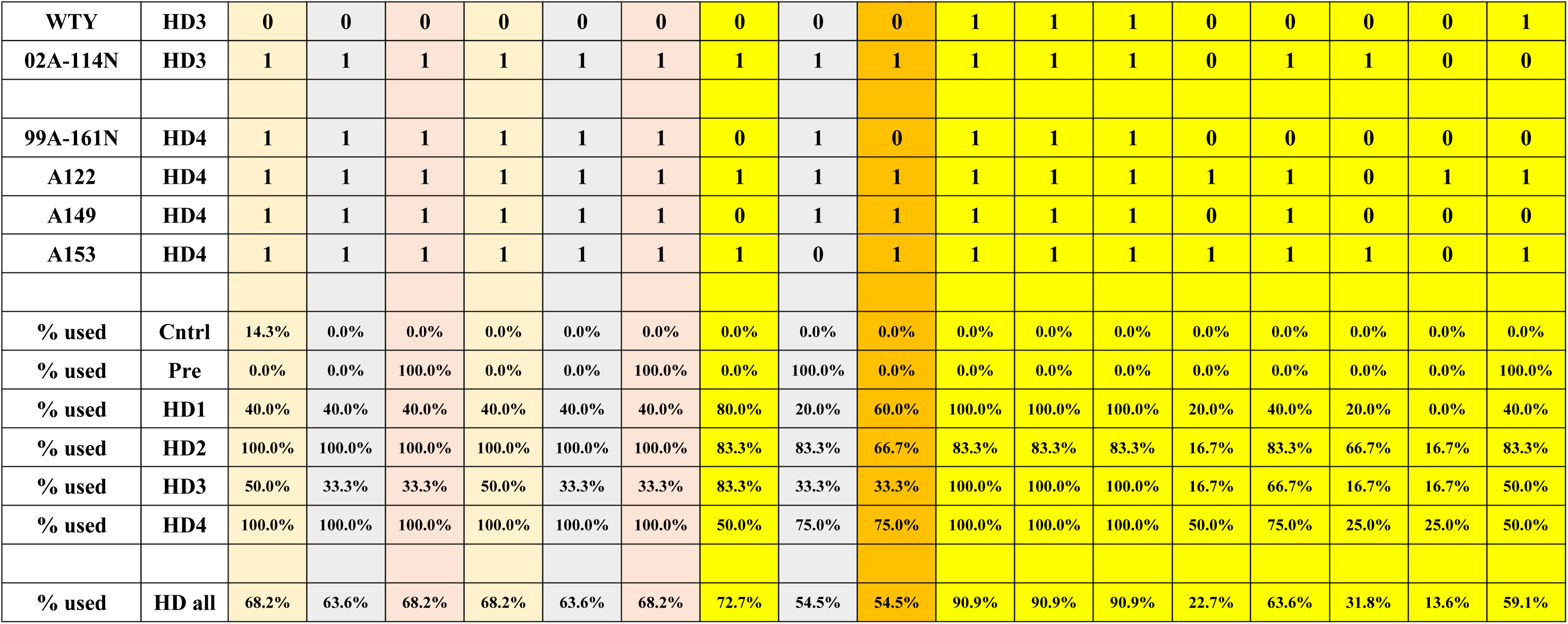
Human case use for each structure examined. See text for abbreviations.

## References

Albin RL, Reiner A, Anderson KD, Penney JB, Young AB. Striatal and nigral neuron subpopulations in rigid Huntington’s disease: implications for the functional anatomy of chorea and rigidity-akinesia. Ann Neurol 1990; 27: 357–365.

Allen Mouse Brain Atlas. Allen Institute for Brain Science 2004; Available from: http://mouse.brain-map.org.

Bakken TE, Jorstad NL, Hu Q, Lake BB, Tian W, Kalmbach BE, Crow M, Hodge RD, Krienen FM, Sorensen SA, Eggermont J, Yao Z, Aevermann BD, Aldridge AI, Bartlett A, Bertagnolli D, Casper T, Castanon RG, Crichton K, Daigle TL, Dalley R, Dee N, Dembrow N, Diep D, Ding SL, Dong W, Fang R, Fischer S, Goldman M, Goldy J, Graybuck LT, Herb BR, Hou X, Kancherla J, Kroll M, Lathia K, van Lew B, Li YE, Liu CS, Liu H, Lucero JD, Mahurkar A, McMillen D, Miller JA, Moussa M, Nery JR, Nicovich PR, Niu SY, Orvis J, Osteen JK, Owen S, Palmer CR, Pham T, Plongthongkum N, Poirion O, Reed NM, Rimorin C, Rivkin A, Romanow WJ, Sedeño-Cortés AE, Siletti K, Somasundaram S, Sulc J, Tieu M, Torkelson A, Tung H, Wang X, Xie F, Yanny AM, Zhang R, Ament SA, Behrens MM, Bravo HC, Chun J, Dobin A, Gillis J, Hertzano R, Hof PR, Höllt T, Horwitz GD, Keene CD, Kharchenko PV, Ko AL, Lelieveldt BP, Luo C, Mukamel EA, Pinto-Duarte A, Preissl S, Regev A, Ren B, Scheuermann RH, Smith K, Spain WJ, White OR, Koch C, Hawrylycz M, Tasic B, Macosko EZ, McCarroll SA, Ting JT, Zeng H, Zhang K, Feng G, Ecker JR, Linnarsson S, Lein ES. Comparative cellular analysis of motor cortex in human, marmoset and mouse. Nature 2021; 598: 111–119.

Bayram-Weston Z, Jones L, Dunnett SB, Brooks SP. Comparison of mHTT antibodies in Huntington’s Disease mouse models reveal specific binding profiles and steady-state ubiquitin levels with disease development. PLoS One 3026; 11: e0155834.

Benn CL, Landles C, Li H, Strand AD, Woodman B, Sathasivam K, Li SH, Ghazi-Noori S, Hockly E, Faruque SM, Cha JH, Sharpe PT, Olson JM, Li XJ, Bates GP. Contribution of nuclear and extranuclear polyQ to neurological phenotypes in mouse models of Huntington’s disease. Human Mol Genetics 2005; 14: 3065–3078.

Bragg RM, Mathews EW, Grindeland A, Cantle JP, Howland D, Vogt T, Carroll JB. Global huntingtin knockout in adult mice leads to fatal neurodegeneration that spares the pancreas. Life Sci Alliance 2024; 7: E202402571.

Brooks SP, Dunnett SB. Mouse models of Huntington’s disease. Curr Top Behav Neurosci 2015; 22: 101–133.

Davies SW, Turmaine M, Cozens BA, DiFiglia M, Sharp AH, Ross CA, Scherzinger E, Wanker EE, Mangiarini L, Bates GP. Formation of neuronal intranuclear inclusions underlies the neurological dysfunction in mice transgenic for the HD mutation. Cell 1997; 90: 537–548.

Deng YP, Penney JB, Young AB, Albin RL, Anderson KD, Reiner A. Differential loss of striatal projection neurons in Huntington’s disease: A quantitative immunohistochemical study. J Chem Neuroanat 2004; 27: 143–164.

Deng, YP, Lei WL, Reiner A. Differential perikaryal localization in rats of D1 and D2 dopamine receptors on striatal projection neuron types identified by retrograde labeling. J Chem Neuroanat 2006; 32: 101–116.

Deng YP, Wong T, Wan JY, Reiner A. Differential early loss of thalamostriatal and corticostriatal input to striatal projection neuron types in the Q140 Huntington’s disease knock-in mouse model. Front Systems Neurosci 2014; 8: 198.

Deng Y, Wang H, Joni M, Sekhri R, Reiner A. Progression of basal ganglia pathology in heterozygous Q175 knock-in Huntington’s disease mice. J Comp Neurol 2021; 529: 1327–1371.

DiFiglia M, Sapp E, Chase KO, Davies SW, Bates GP, Vonsattel JP, Aronin N. Aggregation of huntingtin in neuronal intranuclear inclusions and dystrophic neurites in brain. Science 1997; 277: 1990–1993.

Dragatsis I, Goldowitz D, Del Mar N, Deng YP, Meade CA, Liu L, Sun Z, Dietrich P, Yue J, Reiner A. CAG repeat lengths ≥335 attenuate the phenotype in the R6/2 Huntington’s disease transgenic mouse. Neurobiol Dis 2009; 33: 315–330.

Dragatsis I, Dietrich P, Ren H, Deng YP, Del Mar N, Wang HB, Johnson IM, Jones KR, Reiner A. Effect of early embryonic deletion of huntingtin from pyramidal neurons on the development and long-term survival of neurons in cerebral cortex and striatum. Neurobiol Dis 2018; 111: 102–117.

Franich NR, Hickey MA, Zhu C, Osborne GF, Ali N, Chu T, Bove NH, Lemesre V, Lerner RP, Zeitlin SO, Howland D, Neueder A, Landles C, Bates GP, Chesselet MF. Phenotype onset in Huntington’s disease knock in mice is correlated with the incomplete splicing of the mutant huntingtin gene. J Neuro Res 2019; 97: 1590–1605.

Fusco FR, Chen Q, Lamoreaux WJ, Figueredo-Cardenas G, Jiao Y, Coffman JA, Surmeier DJ, Honig MG, Carlock LR, Reiner A. Cellular localization of Huntingtin in striatal and cortical neurons in rats: Lack of correlation with neuronal vulnerability in Huntington’s disease. J Neurosci 1999; 19: 1189–1202.

Glass M, Dragunow M, Faull RLM. The pattern of neurodegeneration in Huntington’s disease: a comparative study of cannabinoid, dopamine, adenosine and GABAA receptor alterations in the human basal ganglia in Huntington’s disease. Neuroscience 2000; 97: 505–519.

Gu X, Cantle JP, Greiner ER, Lee CY, Barth AM, Gao F, Park CS, Zhang Z, Sandoval-Miller S, Zhang RL, Diamond M, Mody I, Coppola G, Yang XW. N17 modifies mutant huntingtin nuclear pathogenesis and severity of disease in HD BAC transgenic mice. Neuron 2015; 85, 726–741.

Gutekunst CA, Li SH, Yi H, Mulroy JS, Kuemmerle S, Jones R, Rye D, Ferrante RJ, Hersch SM, Li XJ. Nuclear and neuropil aggregates in Huntington’s disease: relationship to neuropathology. J Neurosci 1999; 19: 2522–2534.

Häggkvist J, Tóth M, Tari L, Varnäs K, Svedberg M, Forsberg A, Nag S, Dominguez C, Munoz-Sanjuan I, Bard J, Wityak J, Varrone A, Halldin C, Mrzljak L. Longitudinal small-animal PET imaging of the zQ175 mouse model of Huntington disease shows *in vivo* changes of molecular targets in the striatum and cerebral cortex. J Nucl Med 2017; 58: 617–622.

Handsaker RE, Kashin S, Reed Nm, Tan S, Lee WS, McDonald TM, Morris K, Kamitaki N, Mullally CD, Morakabati NR, Goldman M, Lind G, Kohli R, Lawton E, Hogan M, Ichihara K, Berretta S, McCarroll SS. Long somatic DNA-repeat expansion drives neurodegeneration in Huntington’s disease. Cell 2025; 188: 623–639.

Hausmann FS, Barrett JM, Martin ME, Zhan H, Shepherd GMG. Axonal barcode analysis of pyramidal tract projections from mouse forelimb M1 and M2. J Neurosci 2022; 42: 7733–7743.

Heikkinen T, Lehtimäki K, Vartiainen N, Puoliväli J, Hendricks SJ, Glaser JR, Bradaia A, Wadel K, Touller C, Kontkanen O, Yrjänheikki JM, Buisson B, Howland D, Beaumont V, Munoz-Sanjuan I, Park LC. Characterization of neurophysiological and behavioral changes, MRI brain volumetry and 1H MRS in zQ175 knock-in mouse model of Huntington’s disease. PLoS One 2012; 7: e50717.

Honig MG, Dorian CC, Worthen JD, Micetich AC, Mulder IA, Sanchez KB, Pierce WF, Del Mar NA, Reiner A. Progressive long-term spatial memory loss following repeat concussive and subconcussive brain injury in mice, associated with dorsal hippocampal neuron loss, microglial phenotype shift, and vascular abnormalities. Eur J Neurosci 2021; 54: 5844–5879.

Hoschek F, Natan J, Wagner M, Sathasivam K, Abdelmoez A, von Einem B, Bates GP, Landwehrmeyer GB, Neueder A. Huntingtin HTT1a is generated in a CAG repeat-length-dependent manner in human tissues. Mol Med 2024; 30: 36.

Jansen AH, van Hal M, Op den Kelder IC, Meier RT, de Ruiter AA, Schut MH, Smith DL, Grit C, Brouwer N, Kamphuis W, Boddeke HW, den Dunnen WF, van Roon WM, Bates GP, Hol EM, Reits EA. Frequency of nuclear mutant huntingtin inclusion formation in neurons and glia is cell-type-specific. Glia 65 2017; 50–61.

Jiao Y, Sun Z, Lee T, Fusco FR, Kimble TD, Meade CA, Cuthbertson S, Reiner A. A simple and sensitive antigen retrieval method for free-floating and slide-mounted tissue sections. J Neurosci Methods 1999; 93: 149–162.

Ko J, Ou S, Patterson PH. New anti-huntingtin monoclonal antibodies: Implications for huntingtin conformation and its binding proteins. Brain Res Bull 2001; 56: 319–329.

Ko J, Isas JM, Sabbaugh A, Yoo JH, Pandey NK, Chongtham A, Ladinsky M, Wu WL, Rohweder H, Weiss A, Macdonald D, Munoz-Sanjuan I, Langen R, Patterson PH, Khoshnan A. Identification of distinct conformations associated with monomers and fibril assemblies of mutant huntingtin. Human Mol Gen 2018; 27: 2330–2343.

Kuemmerle S, Gutekunst CA, Klein AM, Li XJ, Li SH, Beal MF, Hersch SM, Ferrante RJ. Huntington aggregates may not predict neuronal death in Huntington’s disease. Ann Neurol 1999; 46: 842–849.

Landles C, Sathasivam K, Weiss A, Woodman B, Moffitt H, Finkbeiner S, Sun B, Gafni J, Ellerby LM, Trottier Y, Richards WG, Osmand A, Paganetti P, Bates GP. Proteolysis of mutant huntingtin produces an exon 1 fragment that accumulates as an aggregated protein in neuronal nuclei in Huntington disease. J Biol Chem 2010; 285: 8808–8823.

Landles C, Milton RE, Jean A, McLarnon S, McAteer SJ, Taxy BA, Osborne GF, Zhang C, Duan W, Howland D, Bates GP. Development of novel bioassays to detect soluble and aggregated huntingtin proteins on three technology platforms. Brain Commun 2021; 3: fcaa231.

Landles C, Osborne GF, Phillips J, Canibano-Pico M, Nita IM, Ali N, Bobkov K, Greene JR, Sathasivam K, Bates GP. Mutant huntingtin protein decreases with CAG repeat expansion: implications for therapeutics and bioassays. Brain Commun 2024; 6: fcae410.

Lei W, Jiao Y, Del Mar N, Reiner A. Evidence for differential cortical input to direct pathway versus indirect pathway striatal projection neurons in rats. J Neurosci 2004; 24: 8289–8299.

Maat-Schieman ML, Dorsman JC, Smoor MA, Siesling S, Van Duinen SG, Verschuuren JJ, den Dunnen JT, Van Ommen GJ, Roos RA. Distribution of inclusions in neuronal nuclei and dystrophic neurites in Huntington disease brain. J Neuropathol Exp Neurol 1999; 58: 129–137.

Mätlik K, Baffuto M, Kus L, Deshmukh AL, Davis DA, Paul MR, Carroll TS, Caron MC, Masson JY, Pearson CE, Heintz N. Cell type specific CAG repeat expansions and toxicity of mutant huntingtin in human striatum and cerebellum. Nature Genetics 2024; 56: 383–394.

Massouh M, Wallman, Pourcher E, MJ, Parent A. The fate of the large striatal interneurons expressing calretinin in huntington’s disease. Neuroscience Research 2008; 62: 216–224.

Meade CA, Deng YP, Fusco FR, Del Mar N, Hersch S, Goldowitz D, Reiner A. Cellular localization and development of neuronal intranuclear inclusions in striatal and cortical neurons in R6/2 transgenic mice. J Com Neurol 2002; 449: 241–269.

Menalled LB, Kudwa AE, Miller S, Fitzpatrick J, Watson-Johnson J, Keating N, Ruiz M, Mushlin R, Alosio W, McConnell K, Connor D, Murphy C, Oakeshott S, Kwan M, Beltran J, Ghavami A, Brunner D, Park LC, Ramboz S, Howland D. Comprehensive behavioral and molecular characterization of a new knock-in mouse model of Huntington’s disease: zQ175. PLoS One 2012; 7: e49838.

Menalled L, Lutz C, Ramboz S, Brunner D, Lager B, Noble S, Park L, Howland D. A Field Guide to Working with Mouse Models of Huntington’s Disease. 2014. The Jackson Laboratory.

Missineo A, Tomei L, Alaimo N, Martufi P, Zavattieri M, Ferrari F, Piai A, Séguin J, Esquina C, Huang N, Wu H-Y, Nicotra E, Pace J, Doherty EM. Highly selective monoclonal antibodies targeting the HTT exon1 neo-epitope. 19^th^ Annual Huntington’s Disease Therapeutics Conference (HDTC), Palm Springs, CA 2024. Poster available at: A. Missineo_anti HTT neo P90 clone 1B12_Poster 23_HDTC 2024.pdf

Nana AL, Kim EH, Thu DC, Oorschot DE, Tippett LJ, Hogg VM, Synek BJ, Roxburgh R, Waldvogel HJ, Faull RL. Widespread heterogeneous neuronal loss across the cerebral cortex in Huntington’s Disease. Journal of Huntington’s Disease 2014; 3:45–64.

Neueder A, Landles C, Ghosh R, Howland D, Myers RH, Faull RLM, Tabrizi SJ, Bates GP. The pathogenic exon 1 HTT protein is produced by incomplete splicing in Huntington’s disease patients. Sci Rep 2017; 7: 1307.

Pressl C, Mätlik K, Kus L, Darnell P, Luo JD, Paul MR, Weiss AR, Liguore W, Carroll TS, Davis DA, McBride J, Heintz N. Selective vulnerability of layer 5a corticostriatal neurons in Huntington’s disease. Neuron 2024; 112: 1–18.

Reiner A, Albin RL, Anderson KD, D’Amato CJ, Penney JB, Young AB. Differential loss of striatal projection neurons in Huntington’s disease. Proc Natl Acad Sci USA 1988; 85: 5733–5737.

Reiner A, Jiao Y, Del mar N, Laverghetta AV, Lei WL. Differential morphology of pyramidal tract-type and intratelencephalically projecting-type corticostriatal neurons and their intrastriatal terminals in rats. J Comp Neurol 2003; 457: 420–440.

Reiner A, Del Mar N, Deng YP, Meade CA, Sun Z, Goldowitz D. R6/2 Neurons with intranuclear inclusions survive for prolonged periods in the brains of chimeric mice. J Comp Neurol 2007; 505: 603–629.

Reiner A, Dragatsis I, Dietrich P. Genetics and neuropathology of Huntington’s disease. Int Rev Neurobiol 2011; 98: 325–372.

Reiner A, Lafferty DC, Wang HB, Del Mar N, Deng YP. The group 2 metabotropic glutamate receptor agonist LY379268 rescues neuronal, neurochemical and motor abnormalities in R6/2 Huntington’s disease mice. Neurobiol Dis 2012; 47: 75–91.

Reiner A, Shelby E, Wang H, DeMarch Z, Deng Y, Guley NH, Hogg V, Roxburgh R, Tippett LJ, Waldvogel HJ, Faull RL. Striatal parvalbuminergic neurons are lost in Huntington’s Disease: Implications for dystonia. Movement Disorders 2013; 28: 1691–1699.

Reiner A, Deng YP. Disrupted striatal inputs and outputs in Huntington’s disease. CNS Neuroscience and Therapeutics 2018; 24: 250–280.

Richfield EK, Vonsattel JP, MacDonald ME, Sun Z, Deng YP, Reiner A. Selective loss of striatal preprotachykinin neurons in a phenocopy of Huntington’s disease. Movement Disorders 2002; 17: 327–332.

Sapp E, Ge P, Aizawa H, Bird E, Penny J, Young AB, Vonsattel JP, DiFiglia M. Evidence for a preferential loss of enkephalin immunoreactivity in the external globus pallidus in low grade Huntington’s disease using high resolution image analysis. Neuroscience 1995; 64: 397–404.

Sapp E, Penney J, Young A, Aronin N, Vonsattel JP, DiFiglia M. Axonal transport of N-terminal huntingtin suggests early pathology of corticostriatal projections in Huntington’s disease. J Neuropathol Exp Neurol 1999; 58: 165–173.

Sapp E, Boudi A, Iwanowicz A, Belgrad J, Miller R, Robertson R, O’Reilly D, Yamada K, Deng Y, Joni M, Li X, Kegel-Gleason K, Khvorova A, Reiner A, Aronin N, DiFiglia M. Mutant huntingtin exon 1 protein detected in mouse brain with neoepitope antibody: effects of CAG repeat expansion, MutS Homolog 3 silencing and aggregation. Brain Commun 2025; 7: fcaf314.

Sathasivam K, Neueder A, Gipson TA, Landles C, Benjamin AC, Bondulich MK, Smith DL, Faull RL, Roos RA, Howland D, Detloff PJ, Housman DE, Bates GP. Aberrant splicing of HTT generates the pathogenic exon 1 protein in Huntington disease. Proc Natl Acad Sci USA 2013; 110: 2366–2370.

Schilling G, Becher MW, Sharp AH, Jinnah HA, Duan K, Kotzuk JA, Slunt HH, Ratovitski T, Cooper JK, Jenkins NA, Copeland NG, Price DL, Ross CA, Borchelt DR. Intranuclear inclusions and neuritic aggregates in transgenic mice expressing a mutant N-terminal fragment of huntingtin. Human Mol Genet 1999; 8: 397–407.

Smith EJ, Sathasivam K, Landles C, Osborne GF, Mason MA, Gomez-Paredes C, Taxy BA, Milton RE, Ast A, Schindler F, Zhang C, Duan W, Wanker EE, Bates GP. Early detection of exon 1 huntingtin aggregation in zQ175 brains by molecular and histological approaches. Brain Commun 2023; 5: fcad010.

Sogorb-Gonzalez M, Landles C, Caron NS, Stam A, Osborne G, Hayden MR, Howland D, van Deventer S, Bates GP, Vallès A, Evers M. Exon 1-targeting miRNA reduces the pathogenic exon 1 HTT protein in Huntington’s disease models. Brain 2024; 147: 4043–4055.

Solem MA, G Pelzel R, Rozema NB, Brown TG, Reid E, Mansky RH, Gomez-Pastor R. Absence of hippocampal pathology persists in the Q175DN mouse model of Huntington’s disease despite elevated HTT aggregation. J Huntington’s Disease 2025; 14: 59–84.

Sorensen SA, Bernard A, Menon V, Royall JJ, Glattfelder KJ, Desta T, Hirokawa K, Mortrud M, Miller JA, Zeng H, Hohmann JG, Jones AR, Lein ES. Correlated gene expression and target specificity demonstrate excitatory projection neuron diversity. Cerebral Cortex 2015; 25: 433–449.

Spargo E, Everall IP, Lantos PL. Neuronal loss in the hippocampus in Huntington’s disease: a comparison with HIV infection. Journal of Neurology, Neurosurgery, and Psychiatry 1993; 56: 487–491.

Thu DC, Oorschot DE, Tippett LJ, Nana AL, Hogg VM, Synek BJ, Luthi-Carter R, Waldvogel HJ, Faull RL. Cell loss in the motor and cingulate cortex correlates with symptomatology in Huntington’s disease. Brain 2010; 133: 1094–1110.

Vitet H, Brandt V, Saudou F. Traffic signaling: new functions of huntingtin and axonal transport in neurological disease. Curr Opin Neurobiol 2020; 63: 122–130.

Vonsattel JP, Myers RH, Stevens TJ, Ferrante RJ, Bird ED, Richardson EP. Neuropathological classification of Huntington’s disease. J Neuropathol Exp Neurol 1985; 44: 559–577.

Vonsattel JP. Huntington disease models and human neuropathology: similarities and differences. Acta Neuropathol 2008; 115: 55–69.

Waelter S, Boeddrich A, Lurz R, Scherzinger E, Lueder G, Lehrach H, Wanker EE. Accumulation of mutant huntingtin fragments in aggresome-like inclusion bodies as a result of insufficient protein degradation. Molecular Biology of the Cell 2001; 12: 1393–1407.

Waldvogel HJ, Kim EH, Tippett LJ, Vonsattel JPG, Faull RLM. The neuropathology of Huntington’s disease. Curr Topics Behav Neurosci 2015; 22: 33–80.

Wang EN, Tydlacka S, Orr AL, Yang SH, Graham RK, Hayden MR, Li S, Chan AWS, Li XJ. Accumulation of N-terminal mutant huntingtin in mouse and monkey models implicated as a pathogenic mechanism in Huntington’s disease. Human Mol Gen 2008; 17: 2738–2751.

Zhao DY, Bäuerlein FJB, Saha I, Hartl FU, Baumeister W, Wilfling F. Autophagy preferentially degrades non-fibrillar polyQ aggregates. Molecular Cell 2024; 84: 1980–1994.

Zuccato C, Cattaneo E. Huntington’s disease. Handbook Experimental Pharmacol 2014; 220: 357–409.

