## Supplemental Table 1 for "Localization of Mutant Huntingtin with HTT Exon1 P90 C-terminal Neoepitope Antibodies in Relation to Regional and Neuronal Vulnerability in Forebrain in Q175 Mice and Human Huntington’s Disease"

### Supplementary Table 1 - Human case use for each structure examined

| Case | Group | A9 | A46 | A6 | A4 | A7 | A22 | A13 | Hipp | Thal | Caud | Put | Str | Acc | GPe | GPi | VP | SN |
|---|---|---|---|---|---|---|---|---|---|---|---|---|---|---|---|---|---|---|
| A91-665N | WT | 0 | 0 | 0 | 0 | 0 | 0 | 0 | 0 | 0 | 0 | 0 | 0 | 0 | 0 | 0 | 0 | 0 |
| ME1910-94 | WT | 0 | 0 | 0 | 0 | 0 | 0 | 0 | 0 | 0 | 0 | 0 | 0 | 0 | 0 | 0 | 0 | 0 |
| A085 | WT | 0 | 0 | 0 | 0 | 0 | 0 | 0 | 0 | 0 | 0 | 0 | 0 | 0 | 0 | 0 | 0 | 0 |
| A111 | WT | 0 | 0 | 0 | 0 | 0 | 0 | 0 | 0 | 0 | 0 | 0 | 0 | 0 | 0 | 0 | 0 | 0 |
| A120 | WT | 0 | 0 | 0 | 0 | 0 | 0 | 0 | 0 | 0 | 0 | 0 | 0 | 0 | 0 | 0 | 0 | 0 |
| A150 | WT | 0 | 0 | 0 | 0 | 0 | 0 | 0 | 0 | 0 | 0 | 0 | 0 | 0 | 0 | 0 | 0 | 0 |
| A136 | WT | 1 | 0 | 0 | 0 | 0 | 0 | 0 | 0 | 0 | 0 | 0 | 0 | 0 | 0 | 0 | 0 | 0 |
| VU-HD | Pre | 0 | 0 | 1 | 0 | 0 | 1 | 0 | 1 | 0 | 0 | 0 | 0 | 0 | 0 | 0 | 0 | 1 |
| A143 | HD1 | 1 | 1 | 1 | 1 | 1 | 1 | 1 | 1 | 1 | 1 | 1 | 1 | 1 | 1 | 0 | 0 | 0 |
| A162 | HD1 | 1 | 1 | 1 | 1 | 1 | 1 | 0 | 0 | 1 | 1 | 1 | 1 | 0 | 1 | 1 | 0 | 0 |
| YB-HD | HD1 | 0 | 0 | 0 | 0 | 0 | 0 | 1 | 0 | 1 | 1 | 1 | 1 | 0 | 0 | 0 | 0 | 0 |
| JC-HD | HD1 | 0 | 0 | 0 | 0 | 0 | 0 | 1 | 0 | 0 | 1 | 1 | 1 | 0 | 0 | 0 | 0 | 1 |
| WTX | HD1 | 0 | 0 | 0 | 0 | 0 | 0 | 1 | 0 | 0 | 1 | 1 | 1 | 0 | 0 | 0 | 0 | 1 |
| A121 | HD2 | 1 | 1 | 1 | 1 | 1 | 1 | 1 | 1 | 1 | 1 | 1 | 1 | 0 | 1 | 1 | 1 | 1 |
| 00A-128N | HD2 | 1 | 1 | 1 | 1 | 1 | 1 | 1 | 1 | 0 | 1 | 1 | 1 | 1 | 1 | 0 | 0 | 1 |
| 98A-171N | HD2 | 1 | 1 | 1 | 1 | 1 | 1 | 0 | 0 | 0 | 0 | 0 | 0 | 0 | 0 | 0 | 0 | 0 |
| 98A-229N | HD2 | 1 | 1 | 1 | 1 | 1 | 1 | 1 | 1 | 1 | 1 | 1 | 1 | 0 | 1 | 1 | 0 | 1 |
| 01A-032N | HD2 | 1 | 1 | 1 | 1 | 1 | 1 | 1 | 1 | 1 | 1 | 1 | 1 | 0 | 1 | 1 | 0 | 1 |
| 99A-233N | HD2 | 1 | 1 | 1 | 1 | 1 | 1 | 1 | 1 | 1 | 1 | 1 | 1 | 0 | 1 | 1 | 0 | 1 |
| 99A-105N | HD3 | 1 | 1 | 1 | 1 | 1 | 1 | 1 | 1 | 1 | 1 | 1 | 1 | 1 | 1 | 0 | 0 | 0 |
| IMH-HD | HD3 | 0 | 0 | 0 | 0 | 0 | 0 | 1 | 0 | 0 | 1 | 1 | 1 | 0 | 1 | 0 | 1 | 1 |
| MR-HD | HD3 | 0 | 0 | 0 | 0 | 0 | 0 | 1 | 0 | 0 | 1 | 1 | 1 | 0 | 0 | 0 | 0 | 1 |
| WTW | HD3 | 1 | 0 | 0 | 1 | 0 | 0 | 1 | 0 | 0 | 1 | 1 | 1 | 0 | 1 | 0 | 0 | 0 |

| | | | | | | | | | | | | | | | | | | |
|---|---|---|---|---|---|---|---|---|---|---|---|---|---|---|---|---|---|---|
| **WTY** | **HD3** | **0** | **0** | **0** | **0** | **0** | **0** | **0** | **0** | **0** | **1** | **1** | **1** | **0** | **0** | **0** | **0** | **1** |
| **02A-114N** | **HD3** | **1** | **1** | **1** | **1** | **1** | **1** | **1** | **1** | **1** | **1** | **1** | **1** | **0** | **1** | **1** | **0** | **0** |
| **99A-161N** | **HD4** | **1** | **1** | **1** | **1** | **1** | **1** | **0** | **1** | **0** | **1** | **1** | **1** | **0** | **0** | **0** | **0** | **0** |
| **A122** | **HD4** | **1** | **1** | **1** | **1** | **1** | **1** | **1** | **1** | **1** | **1** | **1** | **1** | **1** | **1** | **0** | **1** | **1** |
| **A149** | **HD4** | **1** | **1** | **1** | **1** | **1** | **1** | **0** | **1** | **1** | **1** | **1** | **1** | **0** | **1** | **0** | **0** | **0** |
| **A153** | **HD4** | **1** | **1** | **1** | **1** | **1** | **1** | **1** | **0** | **1** | **1** | **1** | **1** | **1** | **1** | **1** | **0** | **1** |
| **% used** | **Cntrl** | **14.3%** | **0.0%** | **0.0%** | **0.0%** | **0.0%** | **0.0%** | **0.0%** | **0.0%** | **0.0%** | **0.0%** | **0.0%** | **0.0%** | **0.0%** | **0.0%** | **0.0%** | **0.0%** | **0.0%** |
| **% used** | **Pre** | **0.0%** | **0.0%** | **100.0%** | **0.0%** | **0.0%** | **100.0%** | **0.0%** | **100.0%** | **0.0%** | **0.0%** | **0.0%** | **0.0%** | **0.0%** | **0.0%** | **0.0%** | **0.0%** | **100.0%** |
| **% used** | **HD1** | **40.0%** | **40.0%** | **40.0%** | **40.0%** | **40.0%** | **40.0%** | **80.0%** | **20.0%** | **60.0%** | **100.0%** | **100.0%** | **100.0%** | **20.0%** | **40.0%** | **20.0%** | **0.0%** | **40.0%** |
| **% used** | **HD2** | **100.0%** | **100.0%** | **100.0%** | **100.0%** | **100.0%** | **100.0%** | **83.3%** | **83.3%** | **66.7%** | **83.3%** | **83.3%** | **83.3%** | **16.7%** | **83.3%** | **66.7%** | **16.7%** | **83.3%** |
| **% used** | **HD3** | **50.0%** | **33.3%** | **33.3%** | **50.0%** | **33.3%** | **33.3%** | **83.3%** | **33.3%** | **33.3%** | **100.0%** | **100.0%** | **100.0%** | **16.7%** | **66.7%** | **16.7%** | **16.7%** | **50.0%** |
| **% used** | **HD4** | **100.0%** | **100.0%** | **100.0%** | **100.0%** | **100.0%** | **100.0%** | **50.0%** | **75.0%** | **75.0%** | **100.0%** | **100.0%** | **100.0%** | **50.0%** | **75.0%** | **25.0%** | **25.0%** | **50.0%** |
| **% used** | **HD all** | **68.2%** | **63.6%** | **68.2%** | **68.2%** | **63.6%** | **68.2%** | **72.7%** | **54.5%** | **54.5%** | **90.9%** | **90.9%** | **90.9%** | **22.7%** | **63.6%** | **31.8%** | **13.6%** | **59.1%** |
