## Supplemental Figures 1-9 for "Localization of Mutant Huntingtin with HTT Exon1 P90 C-terminal Neoepitope Antibodies in Relation to Regional and Neuronal Vulnerability in Forebrain in Q175 Mice and Human Huntington’s Disease"

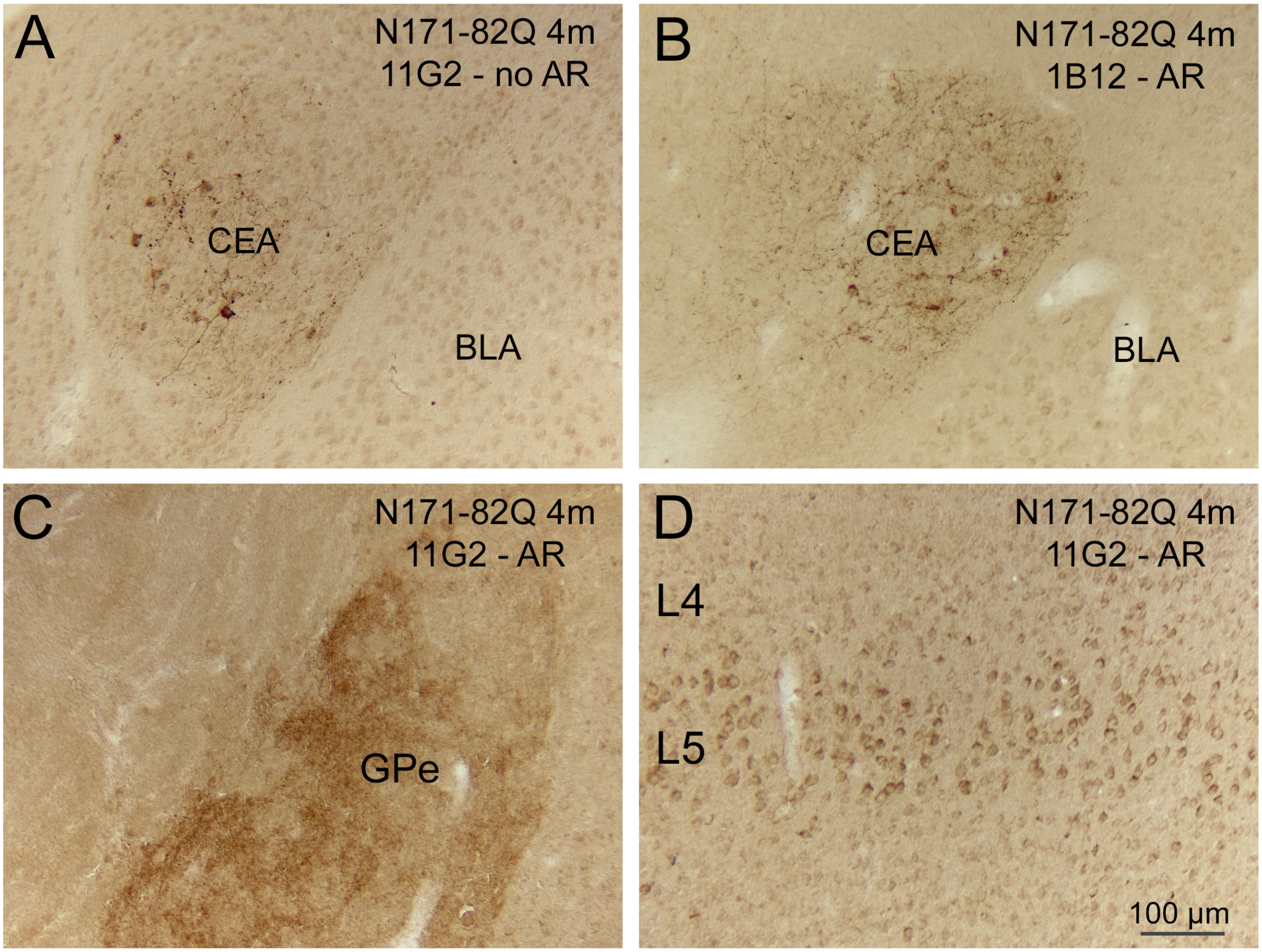


Supplementary Figure 1

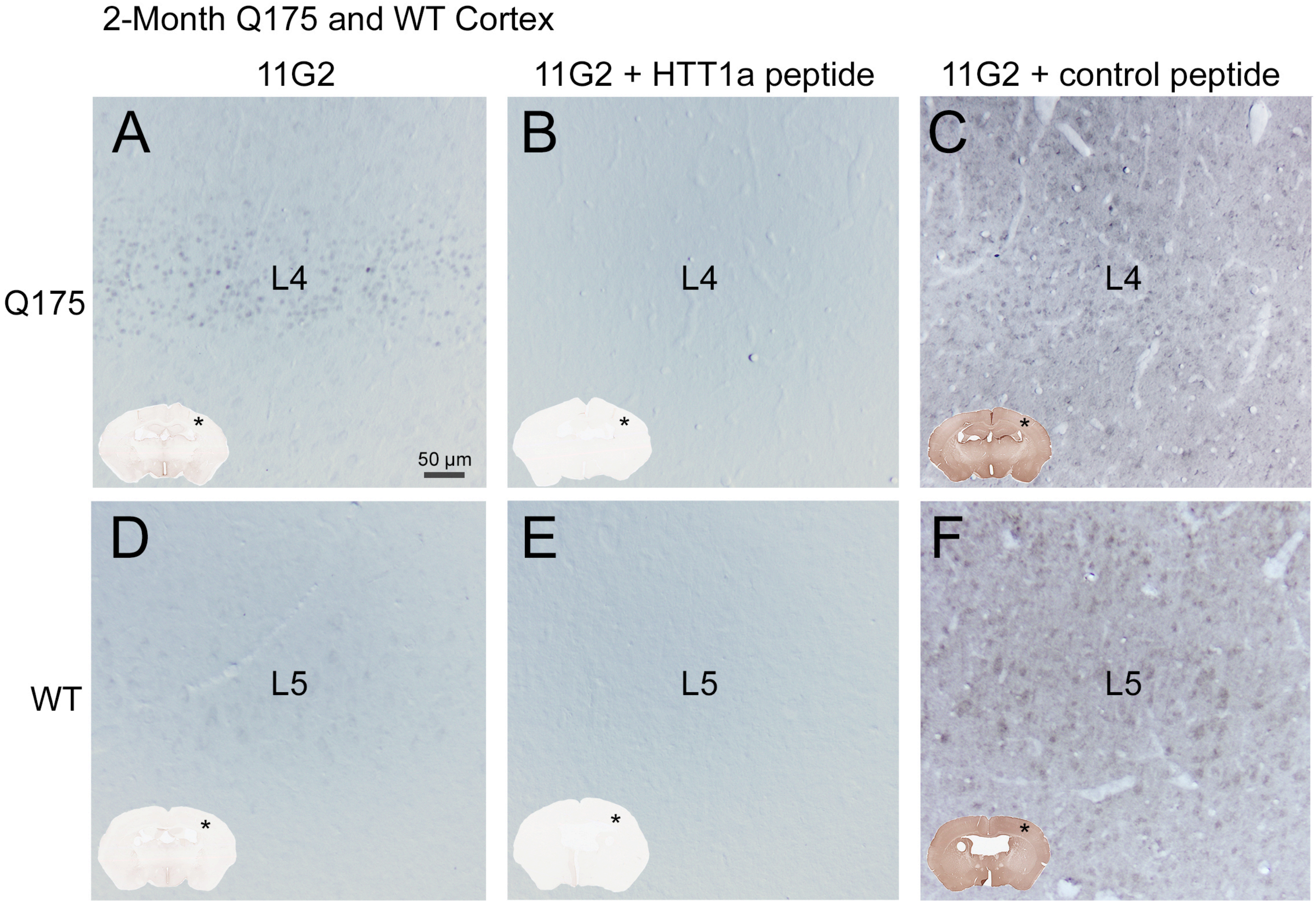


Supplementary Figure 2

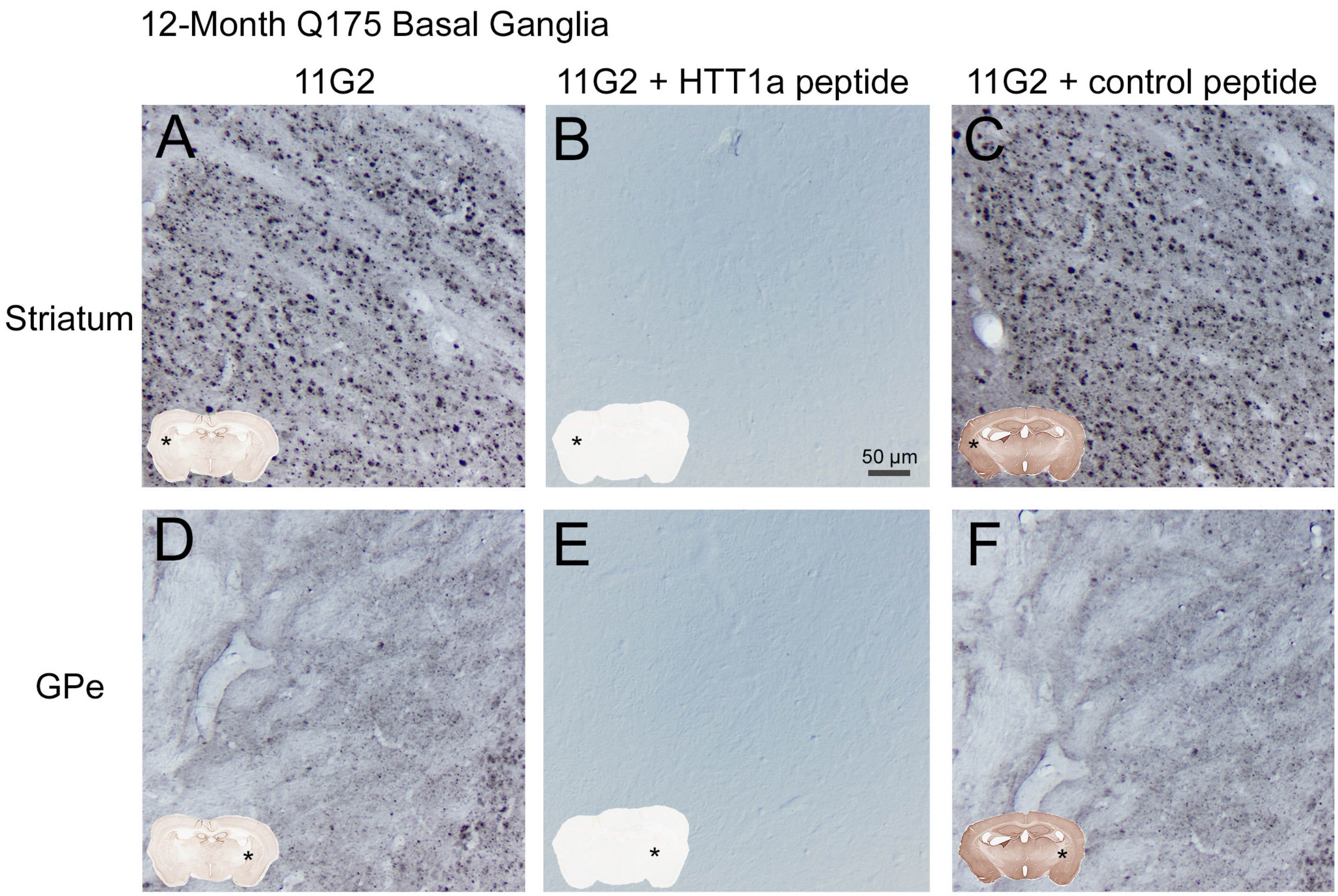


Supplementary Figure 3

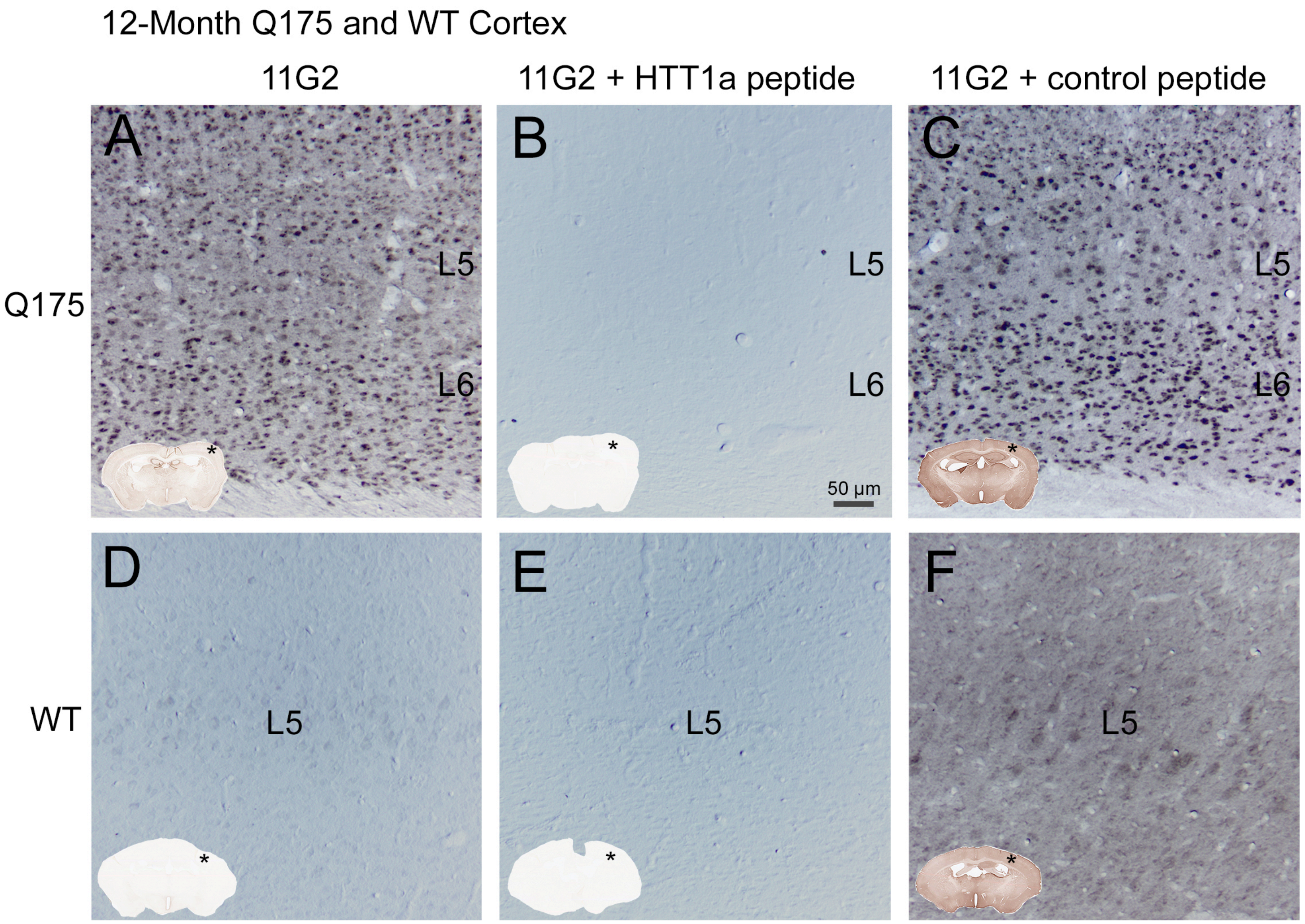


Supplementary Figure 4

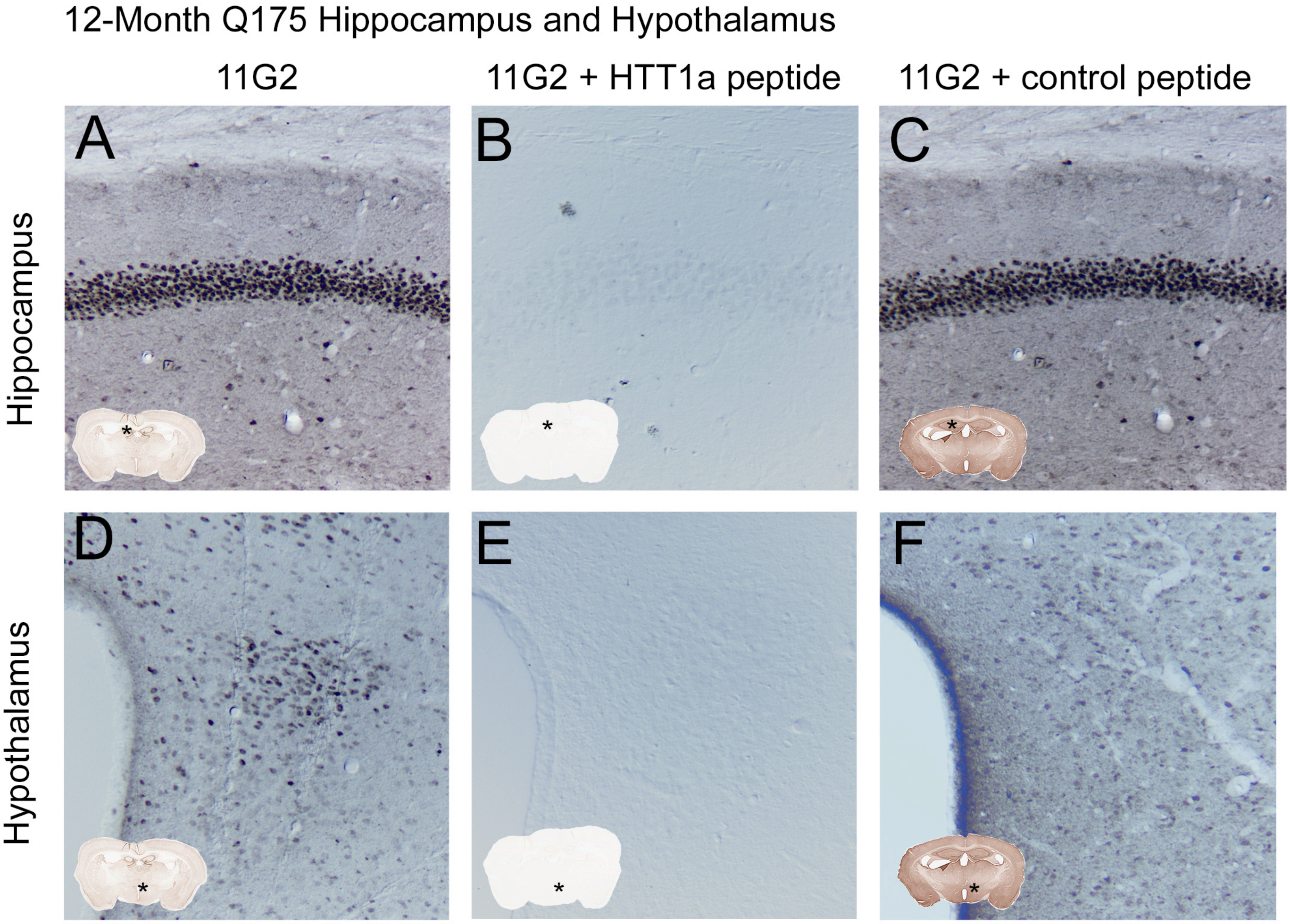


### Supplementary Figure 5

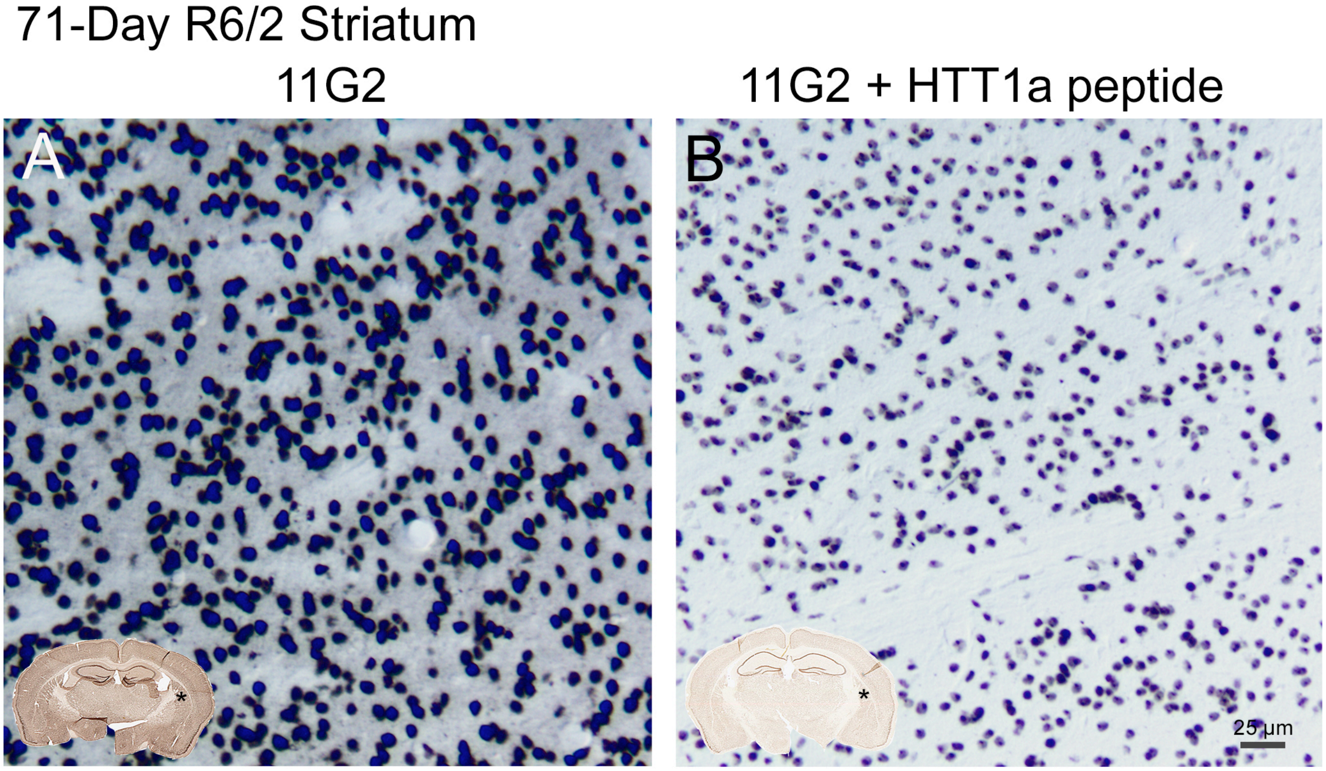


Supplementary Figure 6

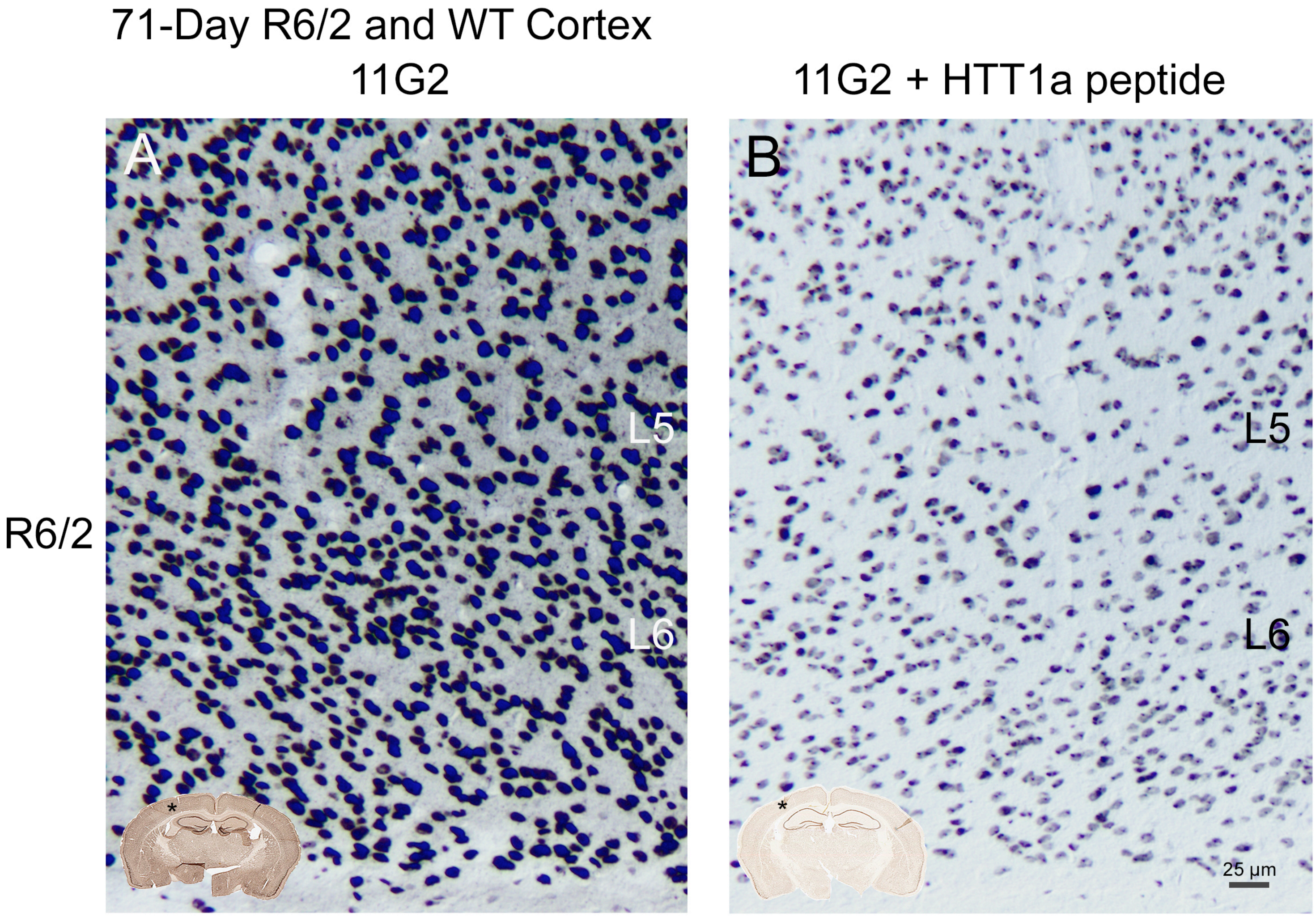


Supplementary Figure 7

Grade 1 Caudate

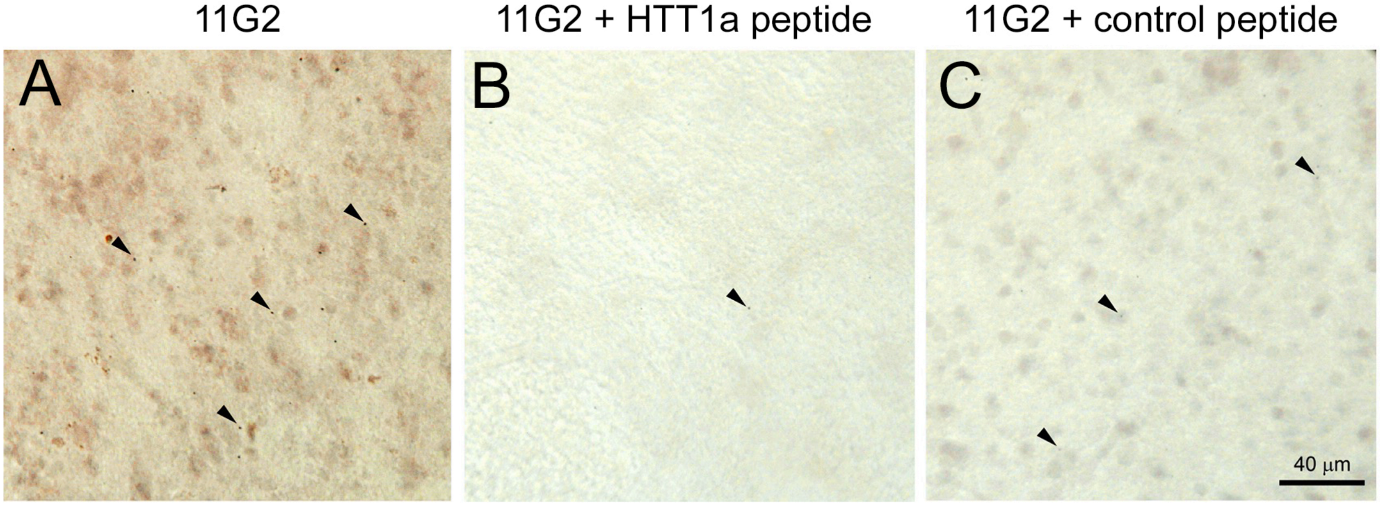


Supplementary Figure 8

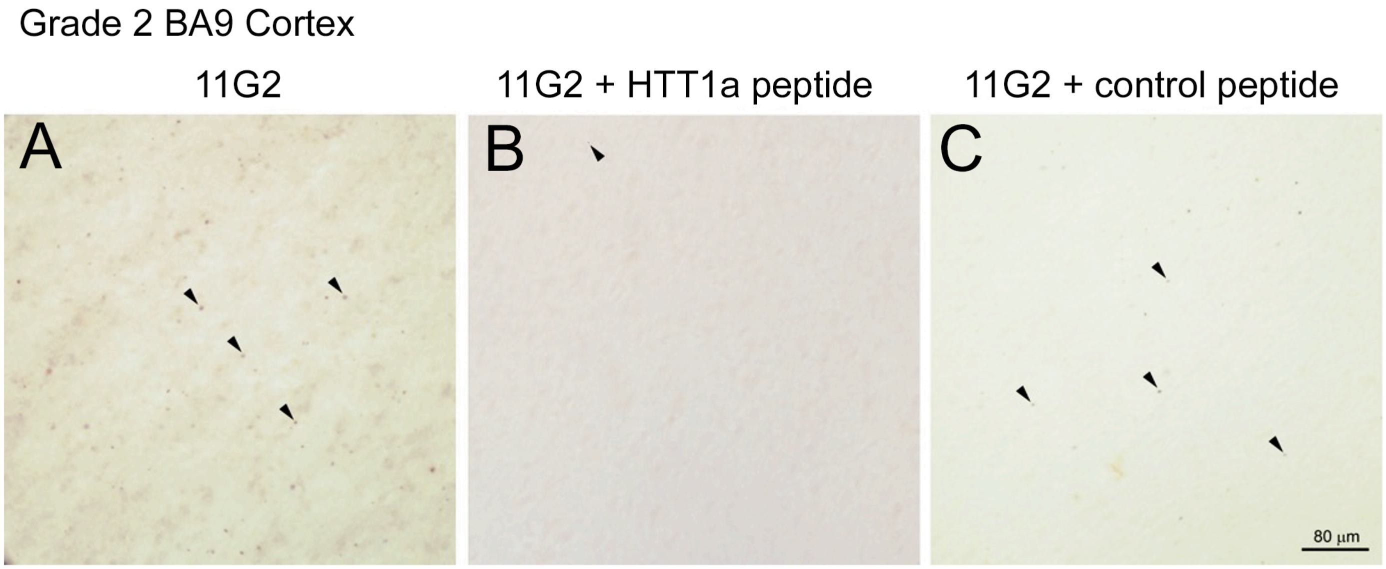


Supplementary Figure 9
